# STAMP-LME: A spatiotemporal surface-mapping framework for population-level iEEG

**DOI:** 10.64898/2026.09.26.754608

**Authors:** Alejandro O. Blenkmann, Sabine L. Leske, Robert T. Knight, Anne-Kristin Solbakk, Robert Oostenveld

**Affiliations:** RITMO Centre for Interdisciplinary Studies in Rhythm, Time and Motion, Department of Psychology, University of Oslo, Noway; Department of Neuropsychology, Clinic for Mental Health and Substance Use, Helgeland Hospital, Mosjøen, Norway; Departments of Neuroscience and Psychology, Helen Wills Neuroscience Institute, UC Berkeley, USA; Department of Neurosurgery, Oslo University Hospital, Norway; Donders Centre for Cognitive Neuroimaging, Radboud University Nijmegen, The Netherlands

**Keywords:** stereoelectroencephalography, group-level analysis, linear mixed-effects models, threshold-free cluster enhancement, population-level inference, cortical mapping

## Abstract

Intracranial EEG (iEEG) affords a unique opportunity to assess human neural activity with millisecond temporal resolution and millimeter spatial precision. However, group-level statistical analyses remain challenging due to sparse and heterogeneous electrode coverage, anatomical variability, and the difficulty of integrating spatiotemporal responses across individuals. Most iEEG studies therefore define brain regions of interest for the group analysis, sacrificing a spatial resolution strength of iEEG and excluding data outside those regions of interest.

Here, we present a spatiotemporal analysis and mapping with permutation-based linear mixed-effects models (STAMP-LME), a cortical-mapping framework for population-level iEEG analysis independent of predefined cortical regions of interest. STAMP-LME is a complete processing pipeline spanning electrode localization, mapping to a standardized surface of the cortical mantle —including medial, orbitofrontal, and insular regions— and statistical inference at the cortical level. Electrode-level data from each participant is mapped to a standardized cortical surface using weights that account for electrode-to-cortex distance and confidence in cortical sampling. Brain signals are then combined with a spatial smoothing kernel and entered into LME models at each vertex-node and time point. Data-derived null distributions are generated through controlled permutations and used for spatiotemporal inference with threshold-free cluster enhancement (TFCE) or false-discovery-rate (FDR) correction.

We validated STAMP-LME using simulations with known cortical ground truth while varying effect size, interparticipant source-location variability, electrode coverage, and spatial smoothing. The framework reliably detected simulated effects and recovered their spatiotemporal distribution while maintaining low localization error and limiting false-positive detections across different sampling conditions. We then applied the method to intracranial recordings from nine individuals performing an auditory oddball task. The empirical analysis revealed distributed auditory cortical responses consistent with conventional channel-level analyses, while providing more fine-grained group-level activation maps across sampled cortical regions.

STAMP-LME provides a flexible, open-source framework for data-driven population-level cortical iEEG analysis that preserves the spatiotemporal strengths of intracranial recordings while accounting for sparse and heterogeneous electrode coverage. The structured workflow enables reproducible and anatomically precise group-level inference on a standardized cortical surface and facilitates comparison with surface-based fMRI and source-localized M/EEG results, therefore providing a practical tool for cognitive and clinical neuroscience research.

**Highlights:**

- STAMP-LME generates population-level cortical maps from sparse SEEG recordings.
- Projection weighting accounts for electrode-to-cortex distance and confidence in cortical sampling.
- Mixed-effects models capture participant- and channel-level variability.
- Data-derived permutations enable ROI-free spatiotemporal inference using TFCE or FDR correction.
- Simulations quantify detection, spatial recovery, and robustness to reduced electrode coverage.

## 1. Introduction

Intracranial EEG (iEEG), including electrocorticography (ECoG) and stereoelectroencephalography (SEEG), provides direct access to human brain activity with exceptional temporal resolution and high spatial precision (Parvizi & Kastner, 2017). This renders iEEG a powerful tool for studying the neural basis of cognitive functions, especially given traditional non-invasive methods (such as scalp EEG or fMRI) temporal resolution limitations. However, a key constraint of iEEG is the lack of a standardized method for group-level statistical analysis (Dubarry et al., 2022; Kadipasaoglu et al., 2014; Mercier et al., 2022; Oehrn, 2023; Trotta et al., 2018). Unlike non-invasive imaging modalities, where data are recorded in consistent spatial layouts, iEEG recordings are highly variable across patients in electrode count, anatomical placement, and coverage. This poses major challenges for statistical inference at the population level.

iEEG have often relied on analyses at the single-subject level, not accounting for inter-subject variability in electrode sampling limiting population-level inference (Kadipasaoglu et al., 2015; Trotta et al., 2018; Mercier et al., 2022). Most studies move beyond case-level analysis using standardized frameworks for group-level inference, including predefined brain regions of interest (ROIs) or aggregating results from functionally selected electrodes (e.g., from a localizer task) (Combrisson et al., 2022; König et al., 2024; Magnotti et al., 2020; Mercier et al., 2022). While informative, these strategies compromise spatial resolution, a major strength of iEEG, in addition to neglecting data sampled outside the defined ROIs.

Statistical group inference also poses challenges besides the spatial organization of the data. Linear mixed-effects (LME) models have been increasingly adopted as a suitable statistical framework for neurophysiological data analysis (Combrisson et al., 2022; Gelman & Hill, 2006; Laird & Ware, 1982; Lv & Wang, 2023; Magnotti et al., 2020; Mercier et al., 2022; Woolnough et al., 2023). LME models allow researchers to simultaneously account for both fixed effects (e.g., stimulus or task condition) and random effects (e.g., subject-level variation), making them suitable for data with unbalanced designs or hierarchical structure, such as those commonly found in iEEG cognitive experiments (Lachaux et al., 2012; Lv & Wang, 2023). LME modeling has been successfully combined with ROIs for statistical inference in the time and frequency domains (Magnotti et al., 2020) and information-theory-derived metrics (Combrisson et al., 2022).

Surface-based mixed-effects multilevel analysis provided an alternative for population-level ECoG mapping by projecting electrode-level effect and precision estimates onto a standardized cortical surface before group inference (Kadipasaoglu et al., 2014). More recently, this approach was extended by constructing for each participant trial-by-trial surface maps and applying vertex-wise LME models to SEEG (Woolnough et al., 2023). However, electrode-level variability is not explicitly represented in the statistical model.

Despite these advances, current approaches remain limited in their ability to handle the spatial characteristics of SEEG data. SEEG is increasingly prevalent in clinical epilepsy monitoring and surgical planning (Abou-Al-Shaar et al., 2018; Dasgupta et al., 2022; Joswig et al., 2018), and there is a need for methods designed to accommodate its properties. Unlike subdural electrodes positioned directly on the cortical surface, SEEG contacts may lie within sulci, gray matter, or near gray–white matter boundaries. Mapping their recordings onto a cortical surface requires consideration of electrode-to-cortex distance and uncertainty regarding which cortical locations contribute to the measured activity (Mercier et al., 2017; von Ellenrieder et al., 2021). These spatial representations must also be integrated with statistical models that account for hierarchical sampling and retain the temporal resolution of intracranial recordings.

Here, we introduce STAMP-LME (spatiotemporal analysis and mapping with permutation-based linear mixed-effects models), a cortical surface-mapping framework for population-level iEEG analysis that is designed for SEEG and readily extendable to ECoG recordings. The pipeline maps electrode-level data from each individual onto a standardized cortical surface (Dale et al., 1999; Fischl, Sereno, Tootell, et al., 1999). The electrode-cortex mapping penalizes plausible cortical sources according to their distance from the recording location and also accounts for the likelihood that several sources may contribute to the recorded activity (i.e., their confidence). At each spatial point (surface’s vertex) and temporal sample, an LME model is fitted to assess the effects of experimental task conditions while accounting for variability across participants and channels nested within participants. In addition, a distance-based spatial weighting is used in the LME computation, which influences how nearby cortically mapped data contribute to the model at each surface vertex. This procedure yields spatially continuous, time-resolved statistical maps on the cortical surface.

Statistical significance is assessed using non-parametric permutation testing. Permutation testing of LME models across all vertices and time points is computationally prohibitive. To address this, we perform permutations on random-effect-corrected (marginal) iEEG data using computationally simpler linear models (Visalli et al., 2024a). The resulting data-derived null distributions are used to obtain permutation-based statistics and to correct for multiple comparisons using either False Discovery Rate (FDR; (Benjamini & Hochberg, 1995a)) or Threshold-Free Cluster Enhancement (TFCE; (Smith & Nichols, 2009)).

High-frequency activity (HFA) is the physiological measure with the highest spatial specificity available via routine iEEG monitoring in epilepsy patients (McCarthy et al., 1999; Parvizi & Kastner, 2017). It is a reliable correlate of neuronal firing (Leszczyński et al., 2020; Mukamel, 2005; Ray & Maunsell, 2011) and has been widely used to study cognitive functions (Johnson et al., 2020; Lachaux et al., 2012; Parvizi & Kastner, 2017; Stolk et al., 2018). We validated STAMP-LME using simulated HFA responses with known cortical ground truth. Simulations were performed across multiple cortical regions while varying effect size, interparticipant source-location variability, electrode coverage, and spatial smoothing. Comparison with the known ground truth simulations allowed quantification of detectability, sensitivity, precision, classification performance, spatiotemporal map similarity, and localization error. These analyses showed that STAMP-LME reliably recovered population-level cortical effects across a range of source and sampling conditions.

To demonstrate the utility of the approach, we applied it to SEEG data from nine patients who performed an auditory oddball task. This paradigm elicits well-characterized neural responses serving as a proof of principle for the method’s ability to reveal group-level spatiotemporal dynamics. Our results showed spatially extended clusters of HFA responses across auditory and associative cortices, confirming the ability of STAMP-LME to detect task condition-specific activations at the group level.

This paper introduces STAMP-LME as a cortical surface-based framework for group-level iEEG analysis, leveraging diffuse cortical mapping, confidence- and distance-weighted signal integration, and vertex-wise linear mixed-effects modeling. The framework enables population-level inference from sparse intracranial recordings while preserving spatial specificity where sampling is sufficient, providing a data-driven approach for cognitive neuroscience and clinical research.

The remainder of this paper is organized as follows: In the Methods section, we describe the STAMP-LME modeling framework, including cortical mapping techniques, group-level spatial smoothing, statistical modeling, and correction procedures, as well as the simulation framework used for validation. The Results section first presents simulation-based evaluations against ground truth, followed by findings from a real case application. Finally, the Discussion situates STAMP-LME within current iEEG analysis approaches and addresses practical considerations, limitations, and possible extensions of the methodological framework.

## 2. Methods

The surface-based linear mixed-effects (STAMP-LME) framework combines anatomical normalization, signal cortical mapping, and statistical modeling into a unified pipeline for group-level iEEG analysis. The key steps of this pipeline are illustrated in Figs. 1 and 2. Broadly, the procedure starts with within-subject processing of brain imaging and electrophysiological data (Fig. 1), followed by group-level statistical modeling on the normalized cortical surface (Fig. 2). Electrode-level signals are diffusely mapped onto the cortex using distance- and confidence-weighted projections, combined across subjects through a spatial smoothing kernel, and analyzed using LME models with permutation-based inference and multiple-comparison correction. The framework is evaluated using both simulated datasets with known ground truth and empirical SEEG recordings.

**Figure 1.**
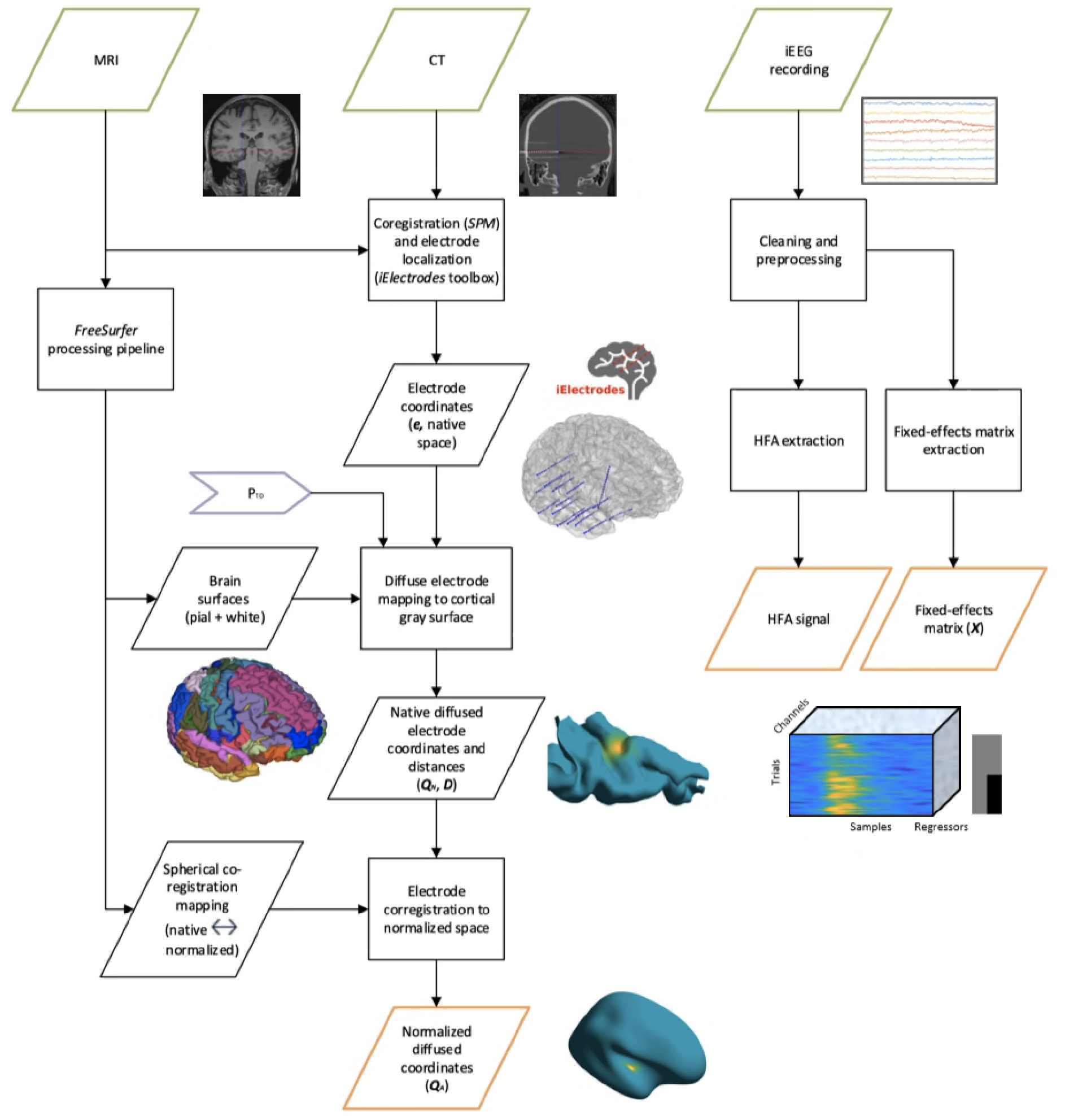
Overview of the within-subject preprocessing pipeline. This figure summarizes the preprocessing steps performed independently for each participant, integrating anatomical imaging and intracranial electrophysiology. The pipeline transforms individual anatomical and electrophysiological data into normalized surface-diffused activity. Structural MRI and postoperative CT images are first coregistered in SPM to localize electrode contacts within native anatomical space. The MRI images are then processed using the FreeSurfer pipeline, which reconstructs the pial and white-matter surfaces and computes spherical registration to a normalized brain template space. Electrode coordinates are extracted from the coregistered MRI–CT data using the iElectrodes toolbox. Each electrode contact is diffusely mapped onto the cortical surface in native space using distance-based weighting (see Fig. 3 for details). Contacts farther than a predefined Threshold Distance (*T_Dist_*) from the cortex are excluded to avoid unreliable projections. The resulting diffuse cortical mappings are subsequently transformed to normalized space via spherical registration, enabling cross-subject alignment. In parallel, intracranial EEG recordings are preprocessed, and high-frequency activity (HFA; 64-245 Hz) is extracted for subsequent statistical group analysis. Diagram notation: parallelogram = data, rectangle = processing step, and arrowhead = method’s parameter.

**Figure 2.**
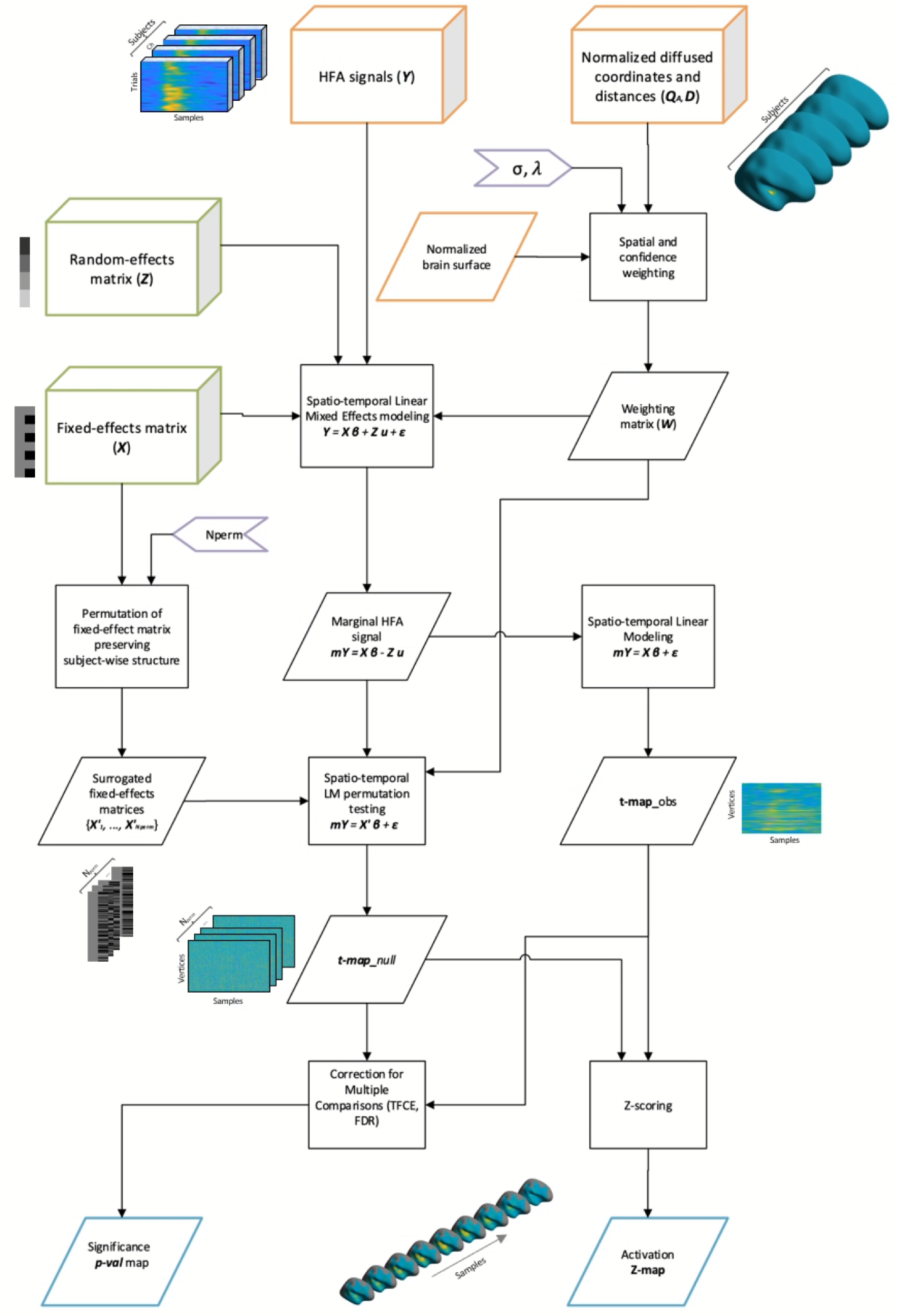
Overview of group-level analysis pipeline. This figure outlines the steps used to integrate channel-level activity across participants and compute vertex-wise (i.e., surface-level) statistical maps. Diffuse cortical mappings and cortical surfaces, aligned to normalized space, are combined to generate a weighting matrix that quantifies each channel’s contribution to each cortical vertex. These weights incorporate both anatomical distance penalization and confidence weighting, reflecting the likelihood that recorded activity originates from a given cortical region. At each vertex and time point, linear mixed-effects (LME) models are fitted to high-frequency activity (HFA), modeling fixed experimental effects and subject-level random effects. Removing the random effects yields marginal HFA signals (mY), which are used to compute the observed vertex-wise t-statistic map (*t-map_obs_*). To assess statistical significance, trial-level labels in the fixed-effects design matrix (X) are randomly permuted *N_perm_* times within subjects, preserving within-subject dependencies. The resulting surrogate design matrices ({*X’_1,_…,X’_Nperm_*}) generate a null distribution of t-maps (*t-map_null_*) through permutation-based linear modeling. Finally, the observed and null distributions are combined to compute Z-scored spatiotemporal activation maps (*Z-map*). Correction for multiple comparisons is performed using TFCE or FDR, yielding the final group-level significance maps. Diagram notation: parallelogram = data, rectangle = processing step, 3D block = multiple-subject dataset, and arrowhead = method’s parameter.

We first describe how MRI and CT scans are coregistered and used to obtain electrode coordinates, which are then mapped onto each individual’s cortical surface and normalized to a standardized anatomical space. We then detail the construction of a vertex-wise weighting scheme that integrates electrode contributions across nearby cortex, incorporating distance penalization, confidence weighting, and group-level spatial smoothing. The statistical modeling stage follows, where vertex- and time-resolved LME models are estimated. Random effects are removed to obtain marginal brain signals, which are then modeled and compared against null distributions derived from within-subject permutations to generate group-level statistical maps. Finally, we describe the procedures for multiple-comparison correction (FDR and TFCE) and for constructing significance maps. Table 1 summarizes the key parameters used in the pipeline.

**Table 1.** Key parameters of the analysis pipeline.

| Step | Parameter Name | Parameter Description | Used value |
| --- | --- | --- | --- |
| Electrode-surface mapping | $T_{Dist}$<br>(threshold distance) | Maximum allowed distance between an electrode and the cortical surface (pial or white) for inclusion in the analysis. | 4 mm |
| | $\lambda$<br>(distance decay constant) | Exponential decay constant controlling distance-based weighting of electrode contributions. Value selected based on the frequency band and reference scheme (bipolar HFA). | 0.36195 |
| | $R_p$<br>(projection radius) | Maximum radius around each electrode that defines the boundary for diffuse projections to be defined. | 8.25 mm |
| Statistical inference | $\sigma$<br>(spatial smoothing constant) | Group-level spatial smoothing parameter. For each vertex, it controls the distance-weighted contribution of the surrounding channels applied via a Gaussian kernel. | 5 mm |
| | $M_{sc}$<br>(minimum subject count) | The minimum number of contributing participants to perform LME modeling at each vertex. | 4 |
|  | E, H<br>(TFCE extent and height parameter) | Threshold Free Cluster Enhancement parameter E weights the extent and parameter H weights the height of clusters. | 0.66, 2 |
| | $N_{perm}$<br>(number of permutations) | Number of permutations used for non-parametric permutation testing. | 2000 |

Together, these steps provide a structured workflow that enables reproducible and anatomically precise group-level inference from heterogeneous iEEG datasets. The subsequent parts of the section introduce the simulation framework used to generate controlled datasets and quantitatively assess method performance. Finally, we describe the empirical SEEG dataset and analysis used to demonstrate the method’s application under realistic recording conditions.

Signal processing and statistical analysis were performed using MATLAB (MathWorks, USA), SPM (Friston et al., 2007), the iElectrodes toolbox (Blenkmann et al., 2017, 2024), FreeSurfer (Fischl, 2012), the FieldTrip toolbox (Oostenveld et al., 2011), the ept-TFCE toolbox (Mensen & Khatami, 2013), iso2mesh toolbox (Fang & Boas, 2009), and custom code. AI assistance (ChatGPT, OpenAI and Github Copilot, Microsoft) was used in developing simulation scripts.

### 2.1. Extraction of cortical surfaces

FreeSurfer is used to extract individual cortical surface models from T1-weighted pre-implantation MRI images. Standard preprocessing includes skull stripping, segmentation of white and gray matter, and the generation of a 3D brain model (Dale et al., 1999; Fischl, Sereno, & Dale, 1999). The last is implemented as a tessellated mesh composed of vertices and triangular faces representing the gray–white matter boundary and the pial surface (the outer boundary of the cortical surface). A mid-gray surface, equidistant between the pial and gray-white matter boundaries, is computed. For the sake of notation simplicity, we refer to these surfaces as white, gray, and pial. Importantly, vertices representing the same cortical column or cortical patch are identically indexed across the three surfaces (i.e., for any coordinate *x* on the surface ***x_N_****__white_* ↔ ***x_N_****__gray_* ↔ ***x_N_****__pial_*).

For each participant, surfaces are aligned to the fsaverage template brain using FreeSurfer’s surface-based spherical registration, enabling cortical alignment across individuals. Native inflated and spherical surface representations, as well as the registration to normalized spherical surface representations, are thus obtained (Fischl, Sereno, Tootell, et al., 1999).

### 2.2. Electrode localization

Intracranial electrode coordinates are localized using post-implantation CT images coregistered to pre-implantation MRI images (SPM12; Studholme et al., 1999). Electrode coordinates ***e*** are extracted for each patient in native space using the iElectrodes toolbox (Blenkmann et al., 2017). Individual electrode artifacts are identified from the coregistered CT using a clustering algorithm (Blenkmann et al., 2017) or by applying the GridFit model-based localization algorithm to all CT artifacts corresponding to an SEEG or ECoG electrode array, thereby improving accuracy (Blenkmann et al., 2024). Coordinates are extracted in the participant’s native MRI space and require visual confirmation.

Bipolar re-referencing is a common approach in iEEG research that enhances the spatial resolution of the recorded signals (McCarty et al., 2022). Virtual coordinates of the bipolar channels may be computed by finding the midpoint between the original electrode locations. The following procedures can be applied to both real and virtual channels. For simplicity, we use the terms electrode or channel in the following to refer to either of them.

### 2.3. Mapping electrodes to the cortical surface

Electrodes capture brain activity from surrounding cortical areas via volume conduction, even when located within white matter (McCarty et al., 2022; Mercier et al., 2022; von Ellenrieder et al., 2021). To avoid unreliable data in the STAMP-LME analysis, only electrodes within the cortical tissue or in its proximity are included. Electrodes farther than the threshold distance (*T_Dist_*) of 4 mm from the white or pial surfaces are excluded.

Then, each single electrode i with coordinates e ^i^ is diffusely mapped to all gray surface vertices within a sphere of radius R_P_, resulting in a set of N_P_ surface-registered coordinates Q ^i^_N_gray_ = {q ^i^_N_gray 1_, …, q ^i^_N_gray Np_ }, with a corresponding set of Euclidean distances D ^i^ = {d ^i^_1_, …, d ^i^_Np_ }, where d ^i^_p_ = || e ^i^ - q ^i^_N_gray p_ || for p = 1, …, N_P_.

A concrete example of this procedure is illustrated in Fig. 3 A-F, which shows how individual electrode contacts are mapped onto the cortical surface in native space and how distance-dependent weights (section 2.5) are distributed across vertices.

**Figure 3.**
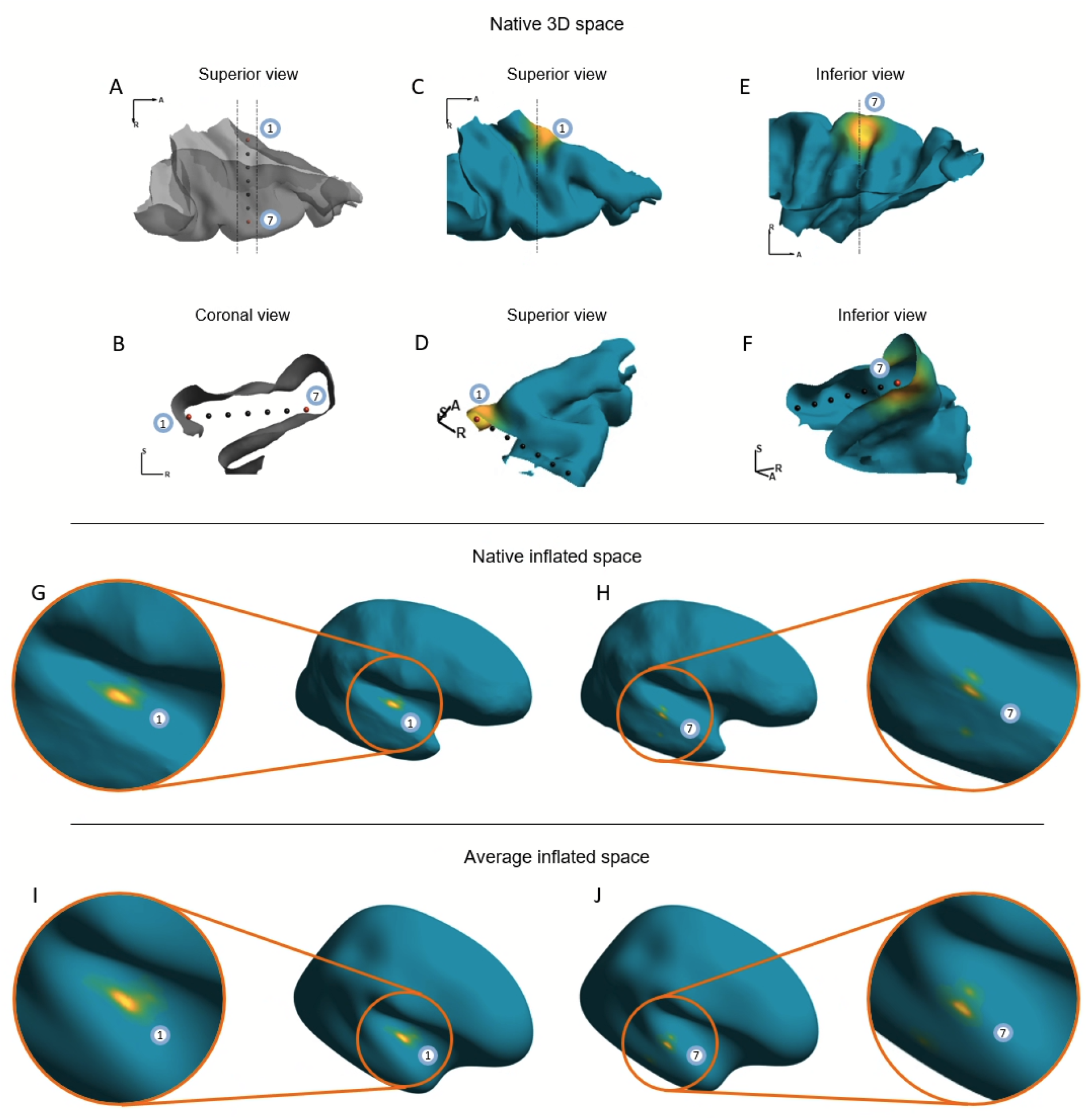
Electrode diffuse cortical mapping in native and normalized space. This figure illustrates how individual intracranial electrodes are diffusely projected onto the vertices of the cortical surface (***Q*** *^i^*). The color coding represents the magnitude of the distance-penalization weights (***W*** *^i^_dist_*, section 2.5.1) at each cortical vertex (warmer colors indicate stronger influence near the electrode). Confidence weighting has been omitted for clarity. Panel A shows a semi-transparent 3D reconstruction of the superior temporal plane in native space (superior view). Seven electrode contacts traverse the region, ordered from deepest (electrode 1) to most lateral (electrode 7). The inner contacts lie immediately beneath the medial aspect of Heschl’s gyrus. Panel B presents a coronal slice indicating the anatomical proximity between the electrode contacts and the surrounding cortical surfaces. Panels C–F demonstrate the diffuse cortical mapping (***Q*** *^i^*_*N_gray*_, for i = 1 and 7) for two representative electrodes. Panels C and D show the mapping for the deepest contact (electrode 1), located beneath the medial aspect of Heschl’s gyrus. Panels E and F illustrate the mapping for the most lateral contact (electrode 7), positioned near the intersection of the superior temporal gyrus (STG) and superior temporal sulcus (STS). Panels G and H display the same distance penalization maps for electrodes 1 and 7 on the participant’s native inflated surface (***Q*** *^i^*_*N_inflatesd*_), which improves visualization by unfolding sulcal anatomy while preserving native geometry. The mapping of electrode 1 remains largely confined to a contiguous medial region of Heschl’s gyrus, reflecting its focal anatomical proximity. In contrast, the mapping of electrode 7 spans across non-contiguous regions of the STG and extends into portions of the middle temporal gyrus (MTG, not shown in previous panels), illustrating how cortical folding and local geometry influence the spatial distribution of weights. Panels I and J show the corresponding mappings (***Q*** *^i^*_*A_inflated*_) after spherical registration to the average cortical surface. This transformation preserves the spatial pattern of weights while enabling alignment across participants for group-level analysis.

As each electrode’s proximity to the cortex varies, N_P_ may differ for each electrode i. Additionally, mappings may distribute across anatomically non-contiguous regions due to cortical folding, as illustrated in Fig. 3 E-F.

Electrodes located within subcortical structures – defined by the anatomical labels obtained earlier – are excluded from the surface mapping pipeline. However, classical ROI-based methods can still be applied to these cases.

Because native gray, inflated, and spherical surfaces share a one-to-one correspondence between vertices, electrode mapping locations can be seamlessly transformed between these representations (***Q*** *^i^*_*N_gray*_ ↔ ***Q*** *^i^*_*N_inflated*_ ↔ ***Q*** *^i^*_*N_spherical*_), as exemplified in Fig. 3G-H.

Given the effects of volume conduction and the spatial spread of iEEG signals (McCarty et al., 2022; von Ellenrieder et al., 2021), we interpret the diffuse cortical mapping locations as potential generators of neural activity, rather than exact signal origins.

### 2.4. Coregistration to normalized cortical surfaces

To enable group-level analysis, the electrodes’ diffused cortical mapping coordinates are remapped from each participant’s native cortical anatomy to a standardized surface-based coordinate system. This transformation is illustrated in Fig. 3G–J, which shows the diffuse mapping of two representative electrodes after spherical registration to the average cortical surface.

FreeSurfer provides such a template —fsaverage— generated by averaging cortical folding patterns from multiple individuals (Fischl, Sereno, Tootell, et al., 1999). This surface-based normalization achieves higher anatomical correspondence across participants than conventional volumetric registration techniques (Coalson et al., 2018; Kadipasaoglu et al., 2014; Trotta et al., 2018). To transfer electrode coordinates to the fsaverage space, FreeSurfer’s spherical surface registration is used, which aligns the native spherical surface to the average spherical curvature pattern extracted from the folding of sulci and gyri (Fischl, Sereno, Tootell, et al., 1999). Therefore, the electrode diffuse mapping coordinates in native space (***Q*** *^i^*_*N_spherical*_) are corregistered to the normalized spherical space (***Q*** *^i^*_*N_spherical*_). Because the individual and the normalized surfaces are sampled (i.e., tessellated) differently, the normalized coordinates (***Q*** *^i^*_*N_spherical*_) are assigned to the nearest vertices on the average brain surface (Trotta et al., 2018). As a result, the new coordinates preserve the one-to-one vertex correspondences across surfaces in the average brain space. Overall, the procedure allows the diffused mappings to be localized across average brain surfaces while preserving anatomical specificity (***Q***^*i*^_*A_white*_ ⟷ ***Q***^*i*^_*A_pial*_ ⟷ ***Q***^*i*^_*A_inflated*_ ⟷ ***Q***^*i*^_*A_spherical*_).

Surface-based LME modeling is performed on the average pial surface (section 2.5), integrating data from electrodes within the neighborhood of each vertex. While the original FreeSurfer surfaces provide very high spatial resolution (at the submillimeter scale), this can lead to prohibitive computational demands for LME. To address this, the pial surfaces are downsampled using the iso2mesh toolbox (Fang & Boas, 2009) to achieve an average inter-vertex spacing of approximately 5 mm. This spacing criterion is user-defined and can be adjusted according to the requirements of each study. Importantly, the spatial downsampling only restricts the vertices on which the LME is modeled, and is not applied to the diffused mapping coordinates, which are kept at the original high resolution.

### 2.5. Surface-based statistical inference

#### 2.5.1. Spatial and confidence weighting

A weighting matrix is needed to describe individual electrodes’ data contributions and aggregate data from different participants at each vertex coordinate in normalized space. Three complementary weightings are applied to address different aspects of the information that electrodes contribute to LME modeling.

- **Distance penalization weighting**. Electrodes farther from the cortical surface are expected to provide less reliable information than those closer to it. Their influence is modeled via an exponential penalization function. A set of weights ***W*** *^i^*_*dist*_ = {*w ^i^_*dist 1*_, …, w ^i^*_*dist Np*_ } is computed for each of the diffused cortical mapping locations, where *w ^i^*_*dist p*_ = exp (-*λ d ^i^_*p*_)*, *d ^i^*_*p*_ is the distance between the electrode i and the mapping coordinate, and *λ* is a decay constant. Fig. 3 C-F shows two examples of distance penalization weights mapped onto the cortex. Based on the spatial correlation observed in HFA, we set *λ* = 0.36195 (McCarty et al., 2022). This decay constant should be adjusted based on the oscillatory frequency band under study and the re-referencing scheme used.
- **Confidence weighting**. If an electrode is closer to a smaller or larger extent of cortex, or several non-contiguous cortices, the confidence in assigning the collected information at the electrode to each cortex point should be attenuated accordingly. Moreover, confidence weighting also depends on mapping distance, assigning higher confidence to closer than to distant cortical areas. Therefore, confidence weights ***W*** *^i^_conf_* = {*w^i^_conf 1_*, …, *w^i^_conf Np_*} are computed for all mapping locations as *w^i^_conf p_* = *w ^i^_dist p_ / W_D_,* where *W_D_ = ∑^Np^_p=1_ w ^i^_dist p_*. Thus, ***W*** *^i^_conf_* is a normalized version of ***W*** *^i^_dist_* (*∑^Np^_p=1_ w^i^_conf p_* = 1).
- **Group-level spatial smoothing.** To integrate data from multiple participants across space, a spatial smoothing kernel weights the contributions of diffuse cortical mappings around a vertex. Given that the LME modeling operates at the average pial surface, the Euclidean distance between each electrode diffused mapping coordinate within ***Q*** ^i^ *_A_pial_* and each pial vertex *v* is computed as *d ^i^_*v-p*_ = || **x**_*v*_ - **q** ^i^_*p*_ ||*. Then, a kernel function is applied to them. In this study, we employed a Gaussian function with a default σ = 5 mm to obtain a smoothing weighting *w ^i^ = exp(- d ^i^ ^2^ / 2*σ^2^). However, other kernel functions can also be implemented. Cortical mapping distances *d ^i^*_*v-p*_> 3σ are ignored as their contribution is negligible. For additional comparison, other values of σ are evaluated (section 2.6).

Overall, each diffused mapped coordinate has the following weights: the distance penalization *w ^i^*_*dist p*_, the confidence *w^i^*_*conf p*_, and the spatial smoothing *w ^i^*_*smooth v-p*_ in connection with all the vertices to which it contributes. To aggregate all the diffused mapping contributions of each electrode *i* into a vertex *v* for LME modeling, weights are combined as *w ^i^*_*v*_ = ∑*^Np^*_*p*=1_ *w ^i^_smooth v-p_ w ^i^_dist p_ w^i^_dist p_*. Finally, matrix ***W*** (N_elec_ x N_vert_) contains all weighting parameters relating electrodes and vertices.

In addition, we define the *cumulative confidence* of an electrode *i* to a vertex *v* as *c^i^_v_* = ∑_*p*_ *w^i^_conf p_* where *p* indexes the diffused mapped points from electrode *i* within a radius of 3σ around vertex *v* (i.e., *d ^i^_v-p_ <* 3σ). This quantity reflects the degree of overlap between the diffuse cortical mapping of electrode *i* and the neighborhood of vertex *v,* and is used to determine which vertices LME is computed on (see section 2.5.2).

The three weighting components described above are illustrated schematically in Fig. 4. Fig. 4A–D depict how different anatomical configurations shape the effective contribution of a single electrode via distance penalization (*w ^i^_dist_*, dashed blue curves) and distance penalization adjusted by confidence weighting (*w ^i^_dist_* ⋅ *w^i^_conf_,* solid orange curves). Fig. 4E illustrates the additional group-level spatial smoothing term (*w _smooth v_,* purple dashed curve), which distributes each electrode’s weighted contribution across nearby cortical vertices.

**Figure 4.**
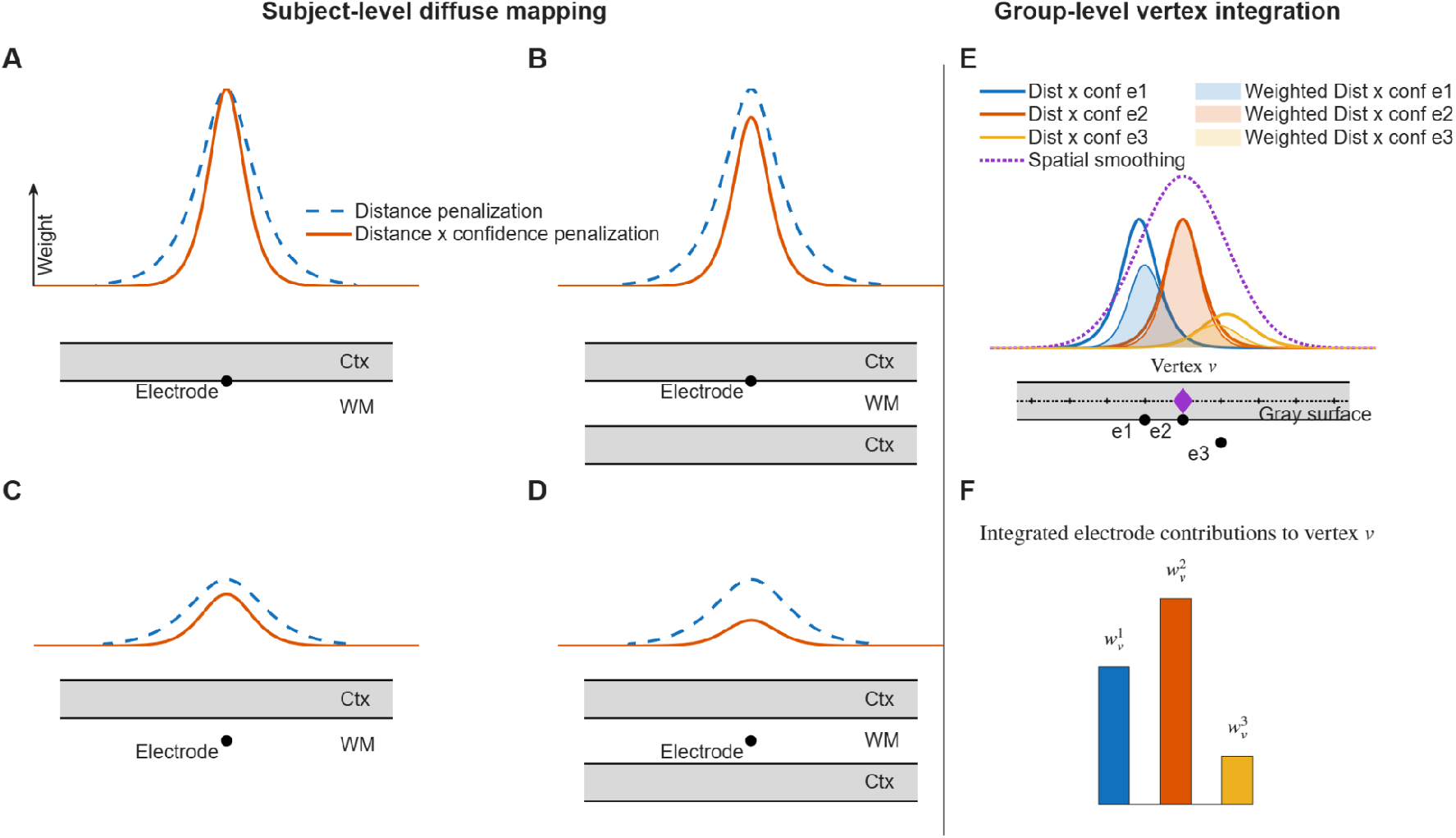
Conceptual illustration of diffuse electrode-to-cortex mapping and group-level integration. This schematic illustrates how activity recorded from intracranial electrodes contributes to cortical vertex LME modeling. The three weighting terms defined in section 2.5.1 — distance penalization (*w_dist_*), confidence normalization (*w_conf_*), and spatial smoothing (*w_smooth_)* — are combined to determine the final electrode-to-vertex weight *w ^i^_v_*used in LME modeling. Panels A–D illustrate how electrode activity is diffusely projected onto the cortical surface under different anatomical configurations. The top curves show two quantities computed along the cortical surface: (i) the distance penalization weight (*w_dist_*, dashed blue line), modeled as an exponential decay with increasing electrode–cortex distance, and (ii) the combined distance × confidence weight (*w_dist_* ⋅ *w_conf_*, solid orange line). Confidence reflects how precisely the recorded signal can be attributed to a given cortical surface and is defined as a normalized version of the distance penalization. Mappings to the lower cortex are omitted for clarity. A. A single electrode lies adjacent to the cortical surface, resulting in strong and unambiguous mapping. B. An additional cortical surface is present beneath the electrode. Although the distance to the upper cortex remains unchanged (and thus *w_dist_* is identical to panel A), confidence is reduced because activity could plausibly originate from either surface. C. The panel shows the effect of increased electrode–cortex distance, leading to reduced distance penalization. D. The panel combines increased distance and anatomical ambiguity (two surrounding cortices), producing the weakest effective mapping to the upper cortex.E. This panel illustrates how multiple electrodes contribute to a cortical vertex *v*. For each electrode, the distance × confidence weights are multiplied by a spatial smoothing kernel (Gaussian function, purple dashed line) centered at vertex *v*. This smoothing kernel models group-level variability and allows spatial integration across nearby cortical locations. The shaded regions represent the spatially weighted contribution of each electrode. F. This panel shows the final integrated weights obtained by summing the shaded areas in panel E. These values quantify the relative contribution of each electrode *i* to vertex *v*. Together, these weights form the matrix ***W*** *(N_elec_* × *N_vert_)* used in the vertex-wise LME models. *Note:* For visualization purposes, curves are shown as continuous functions; in practice, all computations are performed on a discretely sampled cortical surface.

Finally, Fig. 4F represents the integrated vertex-wise weight *w ^i^_v_*, obtained by summing the joint contributions of all diffused mapping points, and forming the entries of matrix **W** used for vertex-wise LME modeling.

#### 2.5.2. Linear-mixed effect modeling

At each vertex coordinate *v,* LME models are estimated using the weighted contributions from surrounding cortically mapped activity. To ensure sufficient statistical support, only pial vertices with a minimum amount of reliable information are included in the analysis. A single electrode is considered to make a *substantial contribution* if the cumulative confidence exceeds 50% (*c^i^*_*v*_ > 0.5, see definition in section 2.5.1). Ultimately, only vertices with *substantial contributions* from at least a *minimum subject count* (MSC) are considered. The MSC threshold is user-defined and should be adjusted according to the study context.

LME is defined for each vertex *v* and sample *t* as:

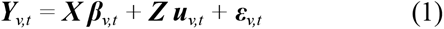

where:

- ***Y****_v,t_* is the vector of observed values (brain activity) at vertex *v* at time *t*. Each observation corresponds to trial-level data from all contributing channels.
- **X** is the fixed-effects design matrix, and ***β****_v,t_* represents the fixed-effects coefficients or regressors at vertex *v* at time *t*.
- ***Z*** is the random-effects design matrix, with ***u****_v,t_* representing the subject-level random regressors at vertex *v* at time *t*.
- ***ε****_v,t_* are the corresponding residual errors.

The dimensions of ***Y****_v,t_, **X**, **Z**,* and ***ε****_v,t_* depend on the number of channels and observations (trials) contributing to each specific vertex. In contrast, the size of ***β*** is constant across the cortical surface vertices and time samples.

Importantly, each observation in ***Y****_v,t_* is weighted by its electrode-to-vertex weight *w ^i^_v_* (i.e., the weighted contribution of data from electrode *i* to vertex *v*; section 2.5.1). These weights are incorporated as observation-level precision weights in the LME estimation, effectively scaling each sample’s contribution to the restricted maximum likelihood (REML) objective function (Gelman & Hill, 2006; Pinheiro & Bates, 2000).

Consequently, observations with higher confidence and closer anatomical proximity exert greater influence on the estimation of both fixed and random effects. Conceptually, this corresponds to a weighted linear mixed-effects model in which contributions with lower mapping certainty are proportionally down-weighted rather than discarded, reducing the impact of anatomically ambiguous or distal mappings while preserving all available information for inference.

Although LME models can incorporate random slopes (i.e., subject-specific variations in experimental condition effects), estimating them requires many additional parameters and may lead to convergence issues and high computational cost. Due to these limitations, and following the approach of Visalli and colleagues (Visalli et al., 2024a), we restricted our model to random intercepts only.

#### 2.5.3. Permutation-based inference using marginal models

Permutation testing is one of the most robust and widely used techniques for statistical inference in neuroimaging research (Maris & Oostenveld, 2007; Nichols & Holmes, 2002; Winkler et al., 2014). By repeatedly shuffling the task-condition labels in the trial data and recomputing test statistics, a null distribution is generated against which the observed results can be compared to assess significance. Permutation testing is a non-parametric approach that does not rely on distributional assumptions, requires that observations are exchangeable under the null hypothesis, and is particularly effective for addressing the multiple comparisons problem when combined with cluster-based statistics (we expand on this idea in section 2.5.4) (Maris & Oostenveld, 2007; Nichols & Holmes, 2002; Winkler et al., 2014).

However, permutation testing is computationally intensive, especially for LME models, which must be refitted for each permutation. To address this limitation, we adopted a marginal model approach as described by Visalli et al. (2024). This approach avoids refitting the full LME model at each permutation, substantially improving computational efficiency. The procedure consists of two main steps:

First, we isolate marginal activity data (***mY***) by removing the random effects from the full model:

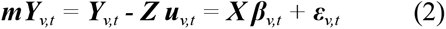

which is computed numerically by adding the fixed-effects predictions and the residuals (obtained by solving eq. 1). Previous work has shown that solving the multiple linear regression problem in equation (2) yields results comparable to those obtained by solving the full LME model (eq. 1) (Visalli et al., 2024a).

By solving the linear regression for each vertex *v = 1,.. ,N_v_* and time point *t = 1,..,N_s_* of ***mY****_v,t_* we obtain:

- a statistical ***t-map****_obs_* (*N_v_* x *N_s_*), and
- a coefficient map ***β***-***map****_obs_* (*N_v_* x *N_s_*) for each fixed effect.

Similar to the LME modeling, the linear regressions are estimated using the weighted contributions from surrounding channels within 3σ around the vertex.

Second, we perform mass-univariate permutation testing using OLS on the marginal activation data (***mY****, N_perm_* permutations). This approach reduces computational cost by several orders of magnitude—approximately 300 times faster than re-fitting LME models for each permutation (Visalli et al., 2024a).

For each permutation, task-condition labels are randomly assigned across trials within each subject, and the corresponding test statistic is recomputed. Thus, within-subject dependence is preserved and exchangeability constraints satisfied (Winkler et al., 2014). This choice ensures valid exchangeability in the presence of hierarchical data and yields more conservative inference in sparse and heterogeneous study designs.

Importantly, for each permutation iteration (***X’****_1_, …, **X’**_Nperm_*), the same randomization of ***X*** is used for all vertices and timepoints to support cluster-based inference (TFCE, section 2.5.4).

The resulting null distribution of t-values for each fixed effect obtained from *Nperm* permutations forms a 3D matrix ***t-map****_null_* (*N_p_* x *N_v_* x *N_s_*). Then, uncorrected p-value maps ***p-val****_uncorr_* (*N_v_* x *N_s_*) are derived as the proportion of null statistics exceeding the observed values (Maris & Oostenveld, 2007).

Using the null distribution of t-values (***t-map****_null_*), we compute, for each vertex and time point, a Z-scored normalized value of the observed statistic, yielding a ***Z-map*** (*N_v_* x *N_s_*) (Cohen, 2014). This normalization expresses each observed effect relative to the distribution expected under the null hypothesis, enabling robust assessment of spatiotemporal activity patterns within a non-parametric, permutation-based statistical method.

#### 2.5.4. Multiple-comparison correction

Given the mass-univariate nature of the STAMP-LME framework, statistical inference requires appropriate control of multiple comparisons across time and space. To address this, we provide two complementary approaches to correcting for multiple comparisons: False Discovery Rate (Benjamini & Hochberg, 1995) and Threshold-Free Cluster Enhancement (Smith & Nichols, 2009).

The FDR method (Benjamini & Hochberg, 1995a) yields FDR-corrected p-value maps ***p-val***_FDRcorr_ (*N_v_* x *N_s_*) from the uncorrected p-value maps ***p-val****_uncorr_* (*N_v_* x *N_s_*), under the assumption of positive or weakly dependent tests, which is commonly satisfied in neuroimaging data (Benjamini & Yekutieli, 2001; Genovese et al., 2002).

While FDR controls the expected false discoveries at the level of individual tests, TFCE provides Family-Wise Error Rate (FWER) control by leveraging the spatiotemporal structure of the data. Neuroimaging data often exhibit strong spatial and temporal correlations, and cluster-based statistics can leverage this structure to improve sensitivity and control for multiple comparisons. A common approach involves identifying clusters of contiguous points (e.g., in time or space) that exceed a fixed statistical threshold, then comparing their size or mass to a null distribution obtained by permuting task-condition labels (Maris & Oostenveld, 2007). While the approach efficiently controls the FWER, it depends critically on the choice of cluster-forming threshold, which can bias results and limit sensitivity to weak or spatially diffuse effects (Mensen & Khatami, 2013; Pernet et al., 2015). To overcome this limitation, the TFCE (Smith & Nichols, 2009) computes an enhanced statistic at each testing point (i.e., vertex and time) that reflects the local support of cluster-like structure, without requiring a predefined threshold. TFCE integrates the height and temporal extent of the statistical signal to elevate real effects in correlated structures.

This method has been shown to improve sensitivity for detecting a wide range of effects in electrophysiological data (Mensen & Khatami, 2013).

TFCE inference is implemented as a maximum-statistic permutation approach. TFCE is computed for each t-statistic map generated during permutation testing in ***t-map****_null_* using surface-based spatial adjacency derived from mesh connectivity and temporal adjacency across consecutive time points. For each permutation surrogate, the maximum TFCE value is extracted and stored in a null distribution vector ***maxTFCE****_null_* (vector of length *N_p_*). The significance threshold *TFCE_thres_*is then defined as the percentile of this distribution corresponding to the chosen significance alpha level. Then, TFCE is applied to the observed statistical map ***t-map****_obs_,*. Vertices and timepoints are considered statistically significant where the TFCE values exceed ***TFCE_t_****_hres_*.

We use the function *ept_mex_TFCE2D* from the ept_TFCE toolbox (http://github.com/Mensen/ept_TFCE-matlab) (Mensen & Khatami, 2013) to compute TFCE maps with extent parameter *E* = 0.66 and height parameter *H* = 2. This configuration enhances sensitivity to both broad and focal effects, making it particularly well suited to heterogeneous iEEG activation patterns.

### 2.6. Performance evaluation via simulations

To evaluate the performance of the proposed STAMP-LME framework under controlled conditions, we conducted a comprehensive simulation study. We generated realistic, task-related high-frequency activity (HFA) signals on the cortical surface across anatomical regions and effect size (Δ) levels. The Group-level activity was then reconstructed for a simulated scenario using the STAMP-LME framework and quantitatively compared against the known ground-truth solutions.

#### 2.6.1. Simulation of cortical surface-level high-frequency activity (HFA)

For the simulation evaluation, we defined nine regions (gyri and sulci) as possible sources of task-related evoked neural activity (see Supp. Table 2 and Supp. Fig 1). Within each simulation case, surface-level HFA was simulated independently for N = 8 subjects on the right-hemisphere inflated FreeSurfer template surface (fsaverage), with an average vertex spacing of approximately 1.5 mm. Neural activity was defined at the level of surface vertices as an additive data generation model, i.e., a signal of interest plus background activity or noise.

**Table 2.** Performance of STAMP-LME using TFCE across spatial smoothing kernels. Detectability, precision, Matthews correlation coefficient (MCC), similarity to the ground truth and centroid-localization error are shown for each combination of source-location jitter, electrode coverage, and simulated effect size. “Best σ” indicates the smoothing kernel or kernels that were not significantly inferior in paired comparisons (corrected for multiple testing). Each table entry was computed after 18 simulations.

| Scenario |  |  | Detectability [%] |  |  |  | Precision [%] |  |  |  | MCC [%] |  |  |  | Similarity [%] |  |  |  | Centroid Error [mm] |  |  |  |
| --- | --- | --- | --- | --- | --- | --- | --- | --- | --- | --- | --- | --- | --- | --- | --- | --- | --- | --- | --- | --- | --- | --- |
| Source Jitter [mm] | Ch | $\Delta$ effect size | $\sigma = 3$ | $\sigma = 5$ | $\sigma = 7$ | Best $\sigma$ | $\sigma = 3$ | $\sigma = 5$ | $\sigma = 7$ | Best $\sigma$ | $\sigma = 3$ | $\sigma = 5$ | $\sigma = 7$ | Best $\sigma$ | $\sigma = 3$ | $\sigma = 5$ | $\sigma = 7$ | Best $\sigma$ | $\sigma = 3$ | $\sigma = 5$ | $\sigma = 7$ | Best $\sigma$ |
| 4 | Full | 0.2 | 88.9 | 83.3 | 83.3 | 3/5/7 | 81.6 | 72.8 | 64.5 | 3 | 58.6 | 60.2 | 47.4 | 3/5 | 66.1 | 56.2 | 53.7 | 3 | 2.96 | 3.30 | 4.98 | 3/5 |
|  |  | 0.3 | 94.4 | 100 | 88.9 | 3/5/7 | 79.6 | 72.3 | 65.5 | 3 | 66.0 | 64.1 | 64.4 | 3/5 | 75.1 | 66.6 | 63.2 | 3 | 2.54 | 3.23 | 3.62 | 3/5 |
|  | 50% | 0.2 | 77.8 | 61.1 | 55.6 | 3/5/7 | 81.4 | 70.5 | 62.4 | 3 | 55.7 | 59.0 | 53.3 | 3/5 | 63.2 | 52.6 | 48.9 | 3 | 2.82 | 3.16 | 3.71 | 3/5 |
|  |  | 0.3 | 88.9 | 94.4 | 72.2 | 3/5/7 | 79.0 | 71.7 | 67.1 | 3/5 | 56.6 | 56.9 | 63.4 | 3/5/7 | 72.4 | 63.8 | 60.1 | 3 | 3.72 | 3.88 | 3.11 | 3/5/7 |
| 8 | Full | 0.2 | 66.7 | 55.6 | 50.0 | 3/5/7 | 58.6 | 58.9 | 60.1 | 3/5/7 | 40.9 | 52.1 | 53.4 | 3/5/7 | 57.4 | 47.7 | 46.8 | 3 | 4.40 | 3.99 | 4.74 | 3/5/7 |
|  |  | 0.3 | 88.9 | 88.9 | 83.3 | 3/5/7 | 69.6 | 64.5 | 57.6 | 3 | 52.0 | 53.8 | 52.0 | 3/5 | 61.1 | 54.6 | 53.5 | 3 | 5.56 | 6.25 | 7.05 | 3/5 |
|  | 50% | 0.2 | 50.0 | 55.6 | 44.4 | 3/5/7 | 59.0 | 59.2 | 51.7 | 3/5/7 | 44.2 | 43.9 | 41.0 | 3/5/7 | 53.5 | 44.4 | 41.9 | 3 | 4.40 | 5.34 | 5.54 | 3/5/7 |
|  |  | 0.3 | 77.8 | 83.3 | 66.7 | 3/5/7 | 72.0 | 66.3 | 58.8 | 3 | 46.8 | 46.0 | 51.7 | 3/5 | 58.1 | 53.0 | 50.5 | 3 | 5.58 | 7.68 | 6.15 | 3/5/7 |

##### Task-related neural activity

Task-related HFA responses were modeled as spatially localized cortical sources centered on predefined vertices. Sources were simulated as disc-shaped patches with a 15 mm diameter, accounting for less than 0.5% of the surface vertices (Cosandier-Rimélé et al., 2008a; Depuydt et al., 2024; Erhardt et al., 2012). Temporally, responses were modeled as smooth, transient envelopes representing trial-locked broadband power modulations (Krol et al., 2018; Visalli et al., 2024b). The temporal profile followed a Gaussian function, with a peak latency of 200 ms, a standard deviation of 50 ms, and variable peak amplitude. Trial length was set to 600 ms, where the first 100 ms correspond to the baseline period. Two experimental conditions were simulated. The peak amplitude of the task-related response was set to 1 μV for condition A and 1 + Δ μV for condition B, where Δ ranged from 0.1 to 0.4 μV, covering effect sizes commonly observed in intracranial broadband HFA. Sampling rate was set to 200 Hz.

##### Background neural activity

Background brain activity is the physiological neural activity that is not part of the experimentally defined source of interest. It has previously been simulated using physiologically plausible models distributed over the cortical surface and linked to dipole generators, thereby producing spatiotemporally correlated activity (A. Blenkmann, 2012; A. Blenkmann et al., 2011; Cosandier-Rimélé et al., 2007, 2008b). Here, we used a simplified approach in which background activity was first simulated independently at each vertex as temporally autocorrelated noise using a first-order autoregressive process. The autocorrelation time constant was set to 20 ms, consistent with reported HFA timescales (Gao et al., 2020), and the noise amplitude was drawn from a zero-mean distribution with a standard deviation of 0.34 μV. Spatial correlations were then introduced by applying a Gaussian surface-smoothing kernel (standard deviation of 3 mm, FWHM ∼7 mm) to both task-related and background signals (McCarty et al., 2022).

##### Subject-level variability

To approximate realistic inter-subject variability, several random effects were introduced:

- **Source location jitter**: the center of each source was randomly displaced tangentially along the cortical inflated surface according to a normal distribution (mean = 0, σ_source_ = 4 mm) (Allen et al., 2012).
- **Subject intercept**: a random additive offset was applied to the entire HFA signal (normal distribution, mean = 0, standard deviation = 0.2 μV).
- **Subject slope**: a random multiplicative scaling factor was applied to task-related responses (normal distribution, mean = 1, standard deviation = 0.2) (Visalli et al., 2024b).

For each condition, 80 trials were generated by adding task-related and background neural activity. Overall, this procedure yielded cortical-level HFA signals capturing key empirical properties of intracranial recordings, including spatially focal activity, temporal smoothness, and trial-to-trial variability.

#### 2.6.2. Simulation of intracranial electrode implantation

To mimic clinically realistic iEEG coverage, virtual intracranial electrodes were implanted on the folded cortical surface following plausible SEEG configurations (Blenkmann et al., 2022). For each subject, we randomly simulated between 4 and 7 SEEG arrays, each consisting of 10 contacts spaced 5 mm apart. Electrode array entry points were randomly distributed within an 80 mm diameter window centered on the closest smooth cortical envelope (SCE) point relative to the simulated brain source. Each array penetrated the cortex approximately orthogonal to the local cortical surface. To mimic variability in implantation trajectories, the surface-normal direction was perturbed by isotropic Gaussian noise, resulting in angular deviations with a mean of 7.17° and a standard deviation of 3.78°.

#### 2.6.3. Simulation of HFA recordings at the electrode level

Electrode-level HFA recordings were reconstructed by combining surface-level activity onto each electrode contact using a distance-weighted spatial integration approach. Specifically, the signal recorded at each electrode was computed as a weighted sum of nearby surface vertices, with weights decaying exponentially with Euclidean distance from the contact point. This procedure approximates the spatial “listening zone” of intracranial electrodes and captures local field averaging effects (McCarty et al., 2022). We implemented a decay constant kₑ = 0.36195, which, at 8 mm, results in an attenuation of ∼95%.

The resulting electrode signals were low-pass filtered at 12.5 Hz, mimicking the temporal smoothing introduced by HFA extraction pipelines that use band-pass filtering and Hilbert transformation for envelope estimation (typical bandwidths can range between 7 and 25 Hz, (Dürschmid et al., 2018; Giroud et al., 2024; Weber et al., 2024)).

Finally, temporally correlated recording-level noise was added to reflect variability not explained by cortical sources (Scheer et al., 2006). Noise was modeled as a zero-mean Gaussian process (standard deviation = 0.15 μV) with a 10 ms temporal autocorrelation, reflecting the temporal smoothing introduced by HFA extraction (band-pass filtering and Hilbert envelope estimation).

The final simulated datasets consisted of trial-resolved HFA time series with dimensions (N_ch_ × N_s_ × N_tr_), preserving the structure required for mass-univariate and mixed-effects analyses.

#### 2.6.4. Ground-truth (GT) activations

To validate the STAMP-LME framework’s performance in simulation scenarios, we must first define the Ground Truth (GT) neural activations. GT activations were defined by performing statistical inference directly on the simulated cortical-level HFA (i.e., independent of electrode-recording reconstruction). Thus, GT reflects the idealized scenario of full and uniform observation at the cortical level.

At each cortical vertex and time point within the trial, an LME model was fitted to the simulated surface-level trial data, using the same fixed- and random-effects structure (condition as fixed effect; subject-level random intercepts and random slopes). In contrast to the STAMP-LME approach, no group-level spatial smoothing was used for inference.

Statistical significance was assessed using within-subject permutation testing on marginal data, yielding vertex- and time-resolved null distributions. Moreover, GT statistical maps were converted to Z-scores for later comparison with STAMP-LME results. Z-scores were obtained by assessing the observed statistics against the permutation-based null distributions. Statistical significance was adjusted for multiple comparisons using FDR and TFCE, yielding spatiotemporally corrected GT activation maps. An activation threshold of a minimum cluster size of seven spatiotemporal connected vertices was applied to avoid sparse detections. This procedure provides the reference against which STAMP-LME reconstructions were evaluated.

#### 2.6.5. Performance metrics and comparisons

##### Comparison between STAMP-LME and GT results

For each region and effect size Δ, we performed 2 simulation runs, yielding a total of 72 simulations for the main analysis. The STAMP-LME analysis of simulated data followed exactly the framework described in the sections above, including diffuse cortical mapping, vertex-wise linear mixed-effects modeling, permutation-based inference, and correction for multiple comparisons. For simulations, we used MSC = 3.

The high-resolution right-hemisphere pial surface (163,842 vertices) was downsampled to 2,459 vertices, increasing the mean inter-vertex spacing to 5.16 mm (median = 5.01 mm). This reduced spatial resolution substantially lowered the computational demands for STAMP-LME, without compromising the ability to generate continuous spatiotemporal activation maps across the cortical surface.

The performance of STAMP-LME was assessed by comparing statistical maps resulting from STAMP-LME with GT statistical maps. All comparisons were restricted to vertices with sufficient electrode sampling, ensuring that evaluation was limited to regions where STAMP-LME inference was possible.

Several complementary performance metrics were computed:

- **Similarity** was computed as the Pearson correlation coefficient between the STAMP-LME and the GT statistical maps across vertices and time points, capturing correspondence in spatial and temporal activation patterns.
- **True Positive Rate (TPR) and False Positive Rate (FPR)** were computed over time and space by comparing STAMP-LME detections with the GT-defined active vertices, thereby quantifying statistical sensitivity and specificity (Mensen & Khatami, 2013). Descriptions were obtained along the time-course of the trial, and along the Euclidean distance to the GT centroid of the neural activity.
- **Centroid localization error** was computed as the Euclidean distance (mm) between the centroids of the STAMP-LME and GT activation clusters to quantify spatial accuracy.

Together, these metrics enable disentangling of detection sensitivity, spatial precision, and overall pattern recovery, providing a comprehensive evaluation of STAMP-LME performance relative to known ground truth.

To assess whether the group-level smoothing coefficient σ affects the spatial distribution of STAMP-LME results and the detection of effects across different scenarios, we evaluated STAMP-LME with smoothing σ = 3, 5, and 7 mm, combined with simulated source-location jitter drawn from zero-mean Gaussian distributions with σ_source_ = 4 and 8 mm, the number of electrodes set to either 100% or 50% of the original electrode coverage, and an effect size Δ = 0.2 and 0.3 μV. For this analysis, 2 simulation runs across the 9 regions were conducted, resulting in a total of 432 additional simulations.

Paired analyses assessed the effects of simulated effect size, electrode coverage, and source-location jitter on STAMP-LME performance. Detectability was compared using exact McNemar tests, whereas continuous performance measures TPR, FPR, precision, Matthews correlation coefficient (MCC; <u>(Chicco & Jurman, 2023)</u>), similarity, and centroid localization error were compared using paired Wilcoxon signed-rank tests and p-values were adjusted using FDR <u>(Benjamini & Hochberg, 1995b)</u>.

##### Comparison between STAMP-LME and classical channel-based group-level detection performance

After validating the ability of the STAMP-LME framework to recover ground-truth activations, we aimed to compare its performance with conventional channel-based statistical analysis. Therefore, we analyzed the simulated recordings using conventional channel-level statistics. For each subject independently, task-condition effects were tested at each channel and time sample using an unpaired two-sample t-test across trials (condition B vs A). Statistical significance was assessed using non-parametric permutation testing by randomly relabeling trials between conditions (N = 2000 permutations), yielding empirical two-sided permutation p-values.

Within each subject, the multiple comparisons problem was addressed using two approaches. First, permutation p-values were FDR-corrected across all channel×time tests. Second, TFCE was applied to the channel×time t-maps using the same parameters (E/H) as in the STAMP-LME cortical surface analysis. As in STAMP-LME analysis, TFCE inference employed a maximum-statistic permutation framework, in which the null distribution was constructed from the maximum absolute TFCE value across the channel×time map for each permutation, thereby controlling the family-wise error rate within each subject. In both cases, a minimum cluster size (nClus = 7) was defined using a 7 mm radius-based channel adjacency and temporal adjacency between neighboring samples, and was used to remove small and spurious detections.

Group-level inference was based on a subject-level detection approach (Ince et al., 2021; Mercier et al., 2022). A subject was considered to show an effect if any channel×time sample within a predefined detection window (100–300 ms) was significant after within-subject correction. The number of subjects showing an effect was then evaluated using a one-sided binomial test against the nominal false positive rate (p = 0.05). Detection rates were summarized as the fraction of simulations in which this group-level test reached significance at each SNR.

### 2.7. Real case application of STAMP-LME

#### 2.7.1. Patients

We recorded data from normal-hearing (self-reported) adults with drug-resistant epilepsy. Patients underwent invasive stereoelectroencephalography (SEEG) recordings as part of their pre-surgical evaluation for resective surgery of the epileptic focus area. Intracranial depth electrodes were temporarily implanted to localize the epileptogenic zone and eloquent cortex. The number and placement of electrodes were guided exclusively by clinical requirements. Data were collected at the Department of Neurosurgery at Oslo University Hospital. Patients were implanted with depth electrodes having 8-18 contacts and an inter-electrode distance of 1.5-5 mm (AdTech, USA, and DIXI Medical, France), as determined by clinical needs. Pre-implantation structural MRI and post-implantation CT scans were acquired for each patient in accordance with standard clinical procedures.

For this study, we selected nine patients (mean age 36 years, range 18 to 50 years, 5 female) from our database who had electrodes covering the temporal and frontal cortices. Frontal and temporal areas were expected to show HFA activations in response to the auditory experimental stimuli used in this validation (see experimental details below). Supp. Table 1 shows additional demographic information. On average, 160 electrodes (range: 101-247) were implanted per patient. Part of the data used in this study were previously reported in (Blenkmann et al., 2019; Fuhrer et al., 2022, 2023, 2025).

This study was approved by the Regional Committees for Medical and Health Research Ethics, Region North Norway (REK, project 2014/381). The research was conducted in accordance with relevant guidelines and regulations. All patients provided written informed consent before participation, in accordance with the World Medical Association Declaration of Helsinki (World Medical Association, 2025).

#### 2.7.2. Experimental design

To validate our framework, we used data collected during an auditory task. We employed the Optimum-1 multi-feature auditory oddball paradigm (A. O. Blenkmann et al., 2019; Näätänen et al., 2004), which included a standard tone and eight different deviant tone types. The standard tone had a duration of 75 ms, with 7 ms linear onset and offset ramps, and consisted of three sinusoidal partials at 500, 1000, and 1500 Hz. Deviants differed relative to the standard in the perceived sound-source location, intensity, frequency, or temporal profile. Standard and deviant tones were interleaved and presented isochronously at 500 ms intervals. Deviant types were pseudo-randomly selected without repetitions. Each block lasted 5 minutes and consisted of 300 standard and 300 deviant tones, preceded by 15 standard tones. Participants were instructed to ignore the sounds while reading a book or magazine. They completed 3 to 10 blocks, yielding at least 1800 trials per participant. Tones were presented through headphones using Psychtoolbox-3 (Kleiner M, Brainard D, 2007) at a comfortable intensity level.

#### 2.7.3. Intracranial EEG data acquisition and signal preprocessing

Intracranial EEG data were recorded using a NicoletOne (Nicolet, Natus Neurology Inc., USA), an ATLAS (Neuralynx, USA), or a BrainQuick (Micromed S.p.A., Italy) system with sampling frequencies of 512, 16000, and 1024/2048 Hz, respectively.

Monopolar iEEG recordings were visually inspected, and channels or epochs showing epileptiform activity or other abnormal signals were removed. Electrodes in lesional or later resected tissue, or those in white matter (except those within one contact of gray matter), were excluded. Bipolar channels were computed as the difference between signals recorded from pairs of neighboring electrodes in the same electrode array. Signals were low-pass filtered at 245 Hz, and line noise was removed using bandstop filters at 50, 100, 150, 200, and 250 Hz (zero-phase filters). Data were then segmented into 2200 ms epochs (950 ms before and 1250 ms after tone onset), demeaned, detrended, and manually inspected for residual artifacts after re-referencing.

To reject remaining outlier trials, we applied an amplitude and spectral rejection criterion within each experimental condition: trials exceeding ±10 median absolute deviations (MAD) from the median amplitude for more than 100 ms, or those with spectral power deviations exceeding 10 MAD across 7 consecutive frequency bins, were excluded. Data were then resampled to 1000 Hz for further analysis.

#### 2.7.4. Extraction of High-Frequency Activity

To obtain the HFA time series, segmented and preprocessed data were bandpass filtered into 12 non-overlapping frequency bands of exponentially increasing bandwidth, ranging from 64 to 245 Hz. Using the default settings in Fieldtrip (ft_preprocessing), the Hilbert transform was applied to each filtered signal, yielding the squared modulus of the complex-valued analytic time series, i.e., the instantaneous power time series. Power signals were then downsampled to 200 Hz and re-epoched between -500 and 1000 ms relative to tone onset to discard edge artifacts.

Each power time series was then z-score normalized relative to a bootstrapped baseline distribution (−100 to 0 ms) (Flinker et al., 2010; Helfrich et al., 2018; Kam et al., 2018). A bootstrapped distribution of baseline values was created by randomly choosing N (number of trials) baseline samples and averaging across these values (the procedure was repeated for 1000 iterations). The power time series were normalized (per frequency band) by subtracting the mean and dividing by the standard deviation (SD) of the baseline distribution. This bootstrapping approach minimizes bias and enhances robustness to statistical outliers. Finally, the 12 power time series were averaged to compute the HFA time series. This procedure implicitly accounts for the 1/f spectral profile of neural signals.

To further control for outliers, trials with values exceeding ±5 MAD from the median for over 50 ms were removed within each experimental condition.

#### 2.7.5. Contrast testing

To demonstrate our framework, we compared responses to Intensity Up deviants (*IntUP*; +6 dB, probability of occurrence 0.05) against Standard tone (*Std*) responses. To reduce computational load, we randomly subsampled *Std* tone responses to match the number of *IntUp* responses per participant. We evaluated differences in the 0 to 400 ms range after tone onset, and used the following LME model:

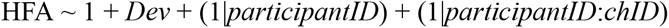

where *Dev* is a binary independent variable encoding *IntUp* deviants (1) or *Std* tones (0). Random intercepts were included for each participant (*participantID*) and for each channel nested within participant (*chID*), capturing between-participant and within-participant (between-channel) variability, respectively. The dependent variable is the rankit-normalized HFA. The rankit transformation maps values to a standard normal distribution based on their rank (Hoy et al., 2023), thereby accommodating the distributional assumptions of the LME regression models (as defined in Section 2.7). Note that permutation-based inference does not require assumptions about the underlying probability distribution of the data.

To simplify the analysis, electrodes on the normalized left cerebral hemisphere were projected onto the corresponding right-hemisphere pial surface. We used an MSC of 4, assuming a small effect variability across subjects, and evaluated results for σ = 5 and 7 mm. For permutation-based statistics, we used *N_perm_* = 2000. The analysis required 9 and 15.1 hours for σ = 5 and 7 mm respectively on a 16-core Linux computer (CPU @ 3.00 GHz × 16; 64 GB RAM).

#### 2.7.6. Comparing STAMP-LME with channel-level t-test

To further assess the validity of our framework, we compared the STAMP-LME results with those obtained from a conventional channel-level statistical approach. For each channel, we performed unpaired-samples t-tests at each time point, comparing responses to Intensity Up deviants (IntUp) versus Standard tones (Std). Multiple comparisons over time were controlled using FDR correction (Benjamini & Hochberg, 1995).

For visualization and comparison, the resulting t-values from each channel were projected onto the cortical surface and averaged using the same distance-weighted kernel function employed in the STAMP-LME framework. This analysis allowed us to assess the extent to which STAMP-LME preserves or enhances spatial HFA activity patterns observed in channel-level analyses that do not account for random effects.

## 3. Results

We evaluated the proposed STAMP-LME framework using both controlled simulations with known ground truth (GT) and empirical iEEG data. Simulations allowed direct quantification of sensitivity, specificity, spatial accuracy, and detection performance across effect sizes, while empirical data illustrated the practical behavior of the analytical approach under realistic recording conditions. Results are presented first for the simulation framework and subsequently for the real data.

### 3.1. Simulation-based performance evaluation of STAMP-LME

Simulated task-related HFA responses were defined at the cortical surface level as spatially localized patches with smooth temporal envelopes. These surface activations were projected onto simulated SEEG contacts via distance weighting, generating realistic channel-level signals with trial-level variability due to noise. Statistical inference was then performed either at the cortical surface level using STAMP-LME or at the electrode level using conventional channel-wise testing. GT activations were defined independent of electrode projections, enabling performance evaluation at the inference level (spatial and temporal correspondence) and at the detection level (sensitivity and specificity).

#### 3.1.1. Simulated ground-truth activations and HFA data

Simulated GT activations consisted of localized cortical patches exhibiting transient HFA increases peaking at 200 ms. At the single-subject level, activations were centered within predefined anatomical regions and varied in position and amplitude due to simulated random effects. Fig. 5A shows a representative simulation instance.

**Figure 5.**
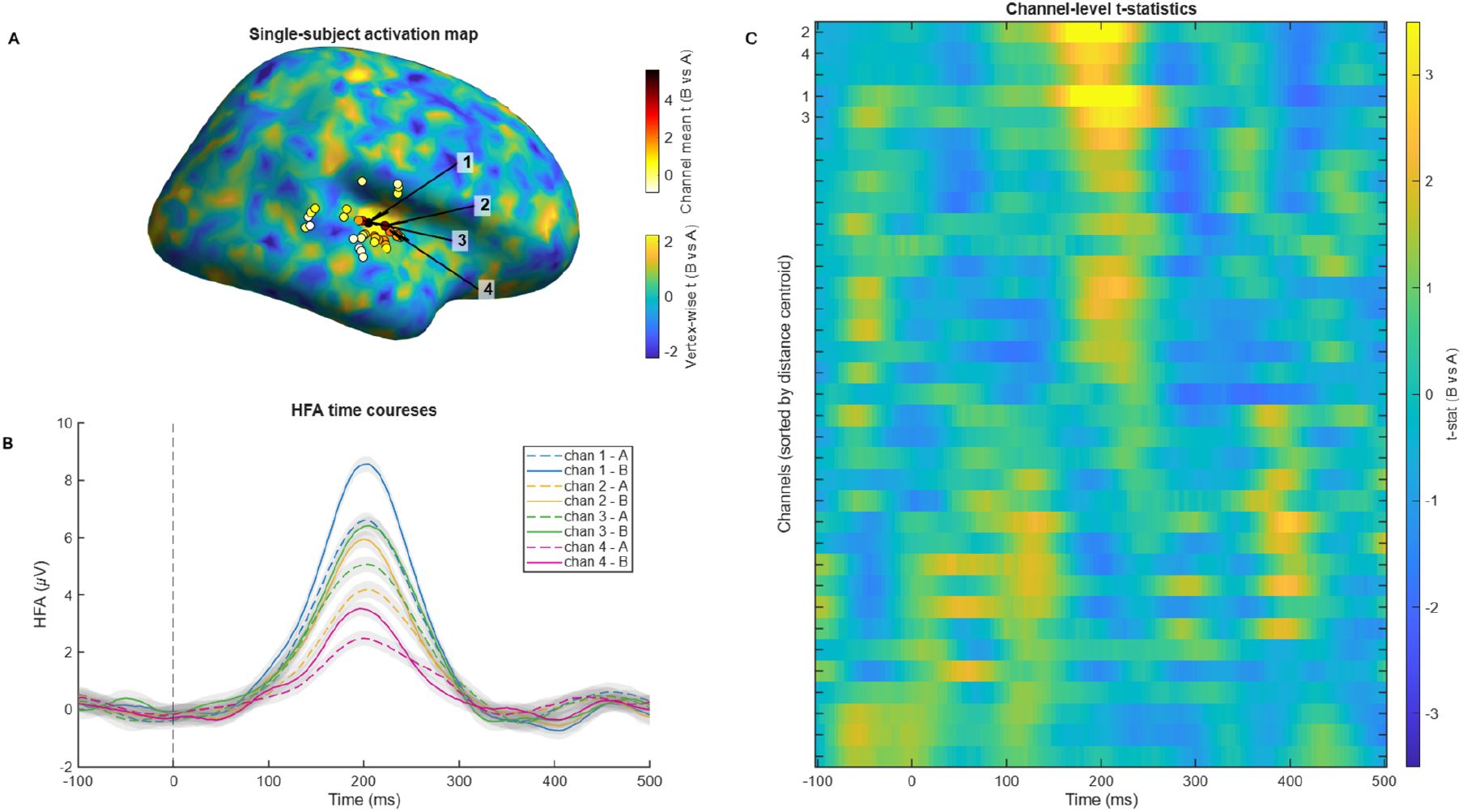
Representative single-subject simulation instance illustrating ground-truth (GT) HFA activity and corresponding channel-level recordings. A. Single-subject cortical activation map derived from vertex-wise t-tests (condition B vs A, effect size Δ = 0.2 μV) on simulated surface-level HFA activity within the 100–300 ms window. Electrodes (projected to the closest cortical surface) are overlaid and color-coded according to each channel’s mean t-statistic. Arrows indicate the four highest-ranked channels (1–4). B. HFA time courses (mean ± SE across trials) for the four highest-ranked channels. Condition A is shown with dashed lines and condition B with solid lines of matching color. C. Channel-level t-statistics (B vs A) across time for all contacts sorted by distance to GT centroid. Channels proximal to the simulated source (top) exhibit robust HFA condition differences centered around 200 ms, whereas distant channels predominantly (bottom) reflect noise-driven activity fluctuations.

Projection of cortical surface activity to SEEG contacts yielded channel-level HFA time series. Channels located proximal to the source exhibited clear experimental condition differences (Fig. 5B). In contrast, distant channels showed predominantly noise-driven fluctuations, as revealed by channel-wise statistical testing across time (i.e., simple t-test contrast, Fig. 5C). In line with these results, GT group-level inference reconstructed continuous cortical activation maps that closely matched the underlying GT simulation parameters within the sampled regions (Fig. 6A, Supp. Fig. 1).

**Figure 6.**
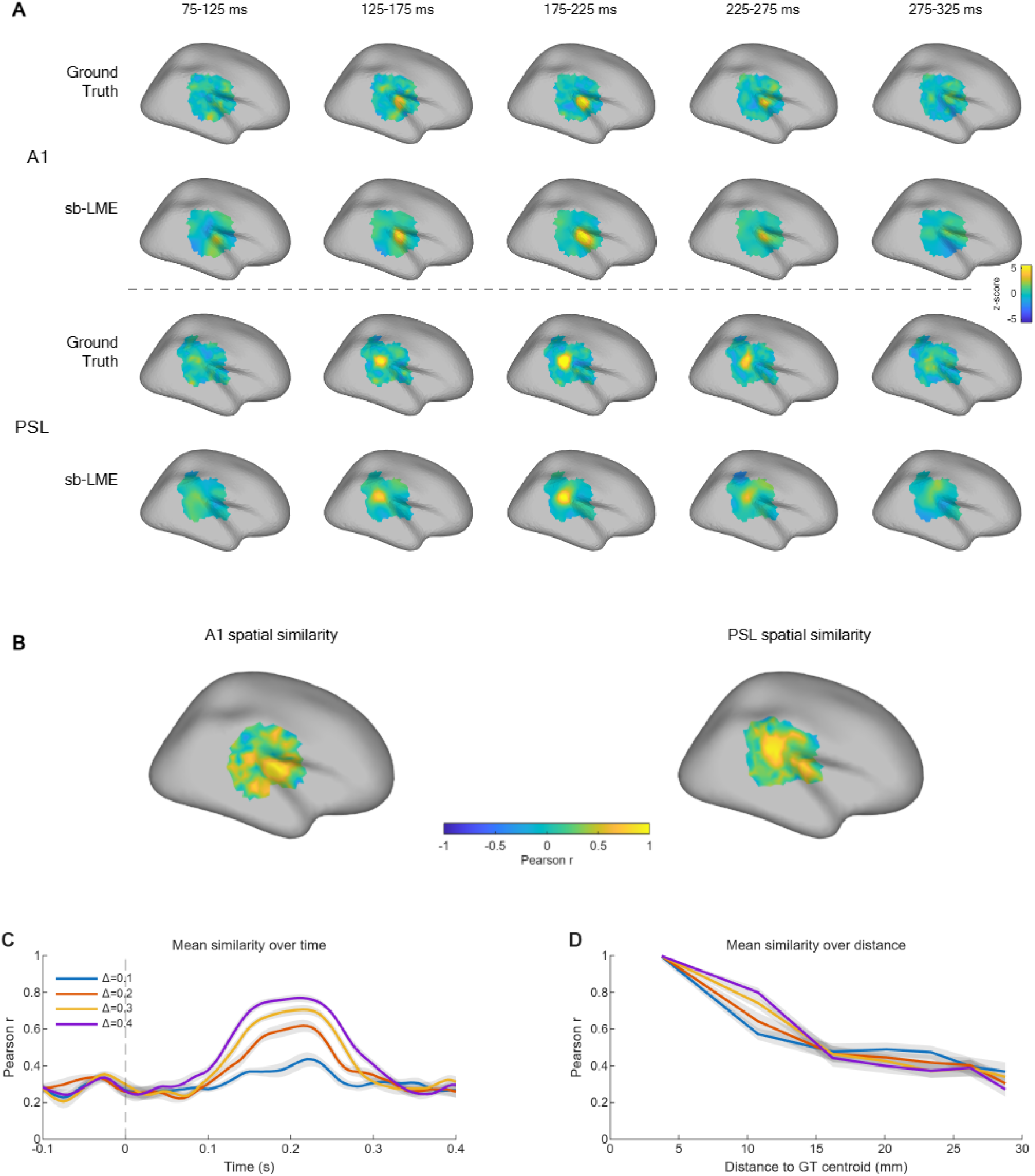
Exemplary cases of STAMP-LME (sb-LME) reconstructions, ground-truth (GT) activation maps, and their corresponding spatiotemporal similarity. A. Example activation Z-score maps for simulated sources in primary auditory cortex (A1) and the PeriSylvian area (PSL) (effect size Δ = 0.2 μV). For each region, GT inference (top rows) and STAMP-LME reconstruction (bottom rows) are shown for successive 50 ms windows between 100 and 300 ms. Despite sparse and heterogeneous electrode sampling across participants, STAMP-LME accurately reconstructs both the spatial location and temporal evolution of the simulated sources. Non-evaluated vertices beyond the sampled areas are depicted in gray color. B. Vertex-wise spatial similarity between the STAMP-LME and GT maps. For each cortical vertex, Pearson’s correlation coefficient between STAMP-LME and GT activity time courses was computed across the task-related interval (0–400 ms) and averaged across simulations. High correlations are localized around the simulated source regions, indicating recovery of the spatial activation pattern. C. Mean similarity between the STAMP-LME and GT maps over time for different simulated effect sizes (Δ). Curves represent the mean Pearson correlation across vertices, averaged across simulations (shaded areas indicate SE). Similarity increases during the task-related interval and peaks near the response maximum (200 ms). In addition, the similarity metric reflects higher effect size (Δ) values. D. Spatial similarity as a function of distance from the GT activation centroid. Pearson correlation between the STAMP-LME and GT maps decreases progressively with increasing distance from the source location, demonstrating that reconstruction accuracy is highest near the true activation center and gradually declines toward the surrounding cortex, consistent with the patterns observed in panel B.

#### 3.1.2. STAMP-LME spatiotemporal similarity to GT

STAMP-LME reconstructed continuous cortical surface-based activation maps that closely matched the GT spatial pattern within the sampled regions, despite sparse and heterogeneous electrode coverage on the individual basis (Supp. Fig. 1, Supp. Fig 2). Fig. 6A shows GT inference of two simulation instances with sources in A1 and SPL and their respective STAMP-LME reconstructions.

Spatiotemporal correspondence between STAMP-LME and group-level GT maps was assessed using Pearson’s correlation coefficient across vertices and time points (Fig. 6B). Similarity increased sharply during the task-related interval and peaked near the response maximum (Fig. 6C). Similarity peaked at the activation centroid location and decayed with distance from it (Fig. 6D).

#### 3.1.3. STAMP-LME detection performance relative to Ground Truth

We next evaluated the ability of STAMP-LME to detect the simulated condition effect relative to the GT significance maps. Figure 7A illustrates representative examples of GT detections and those obtained from TFCE- and FRD-corrected STAMP-LME maps. The spatial location of significant vertices and clusters closely matched the GT source region, with minimal detections outside the activation patch.

**Figure 7.**
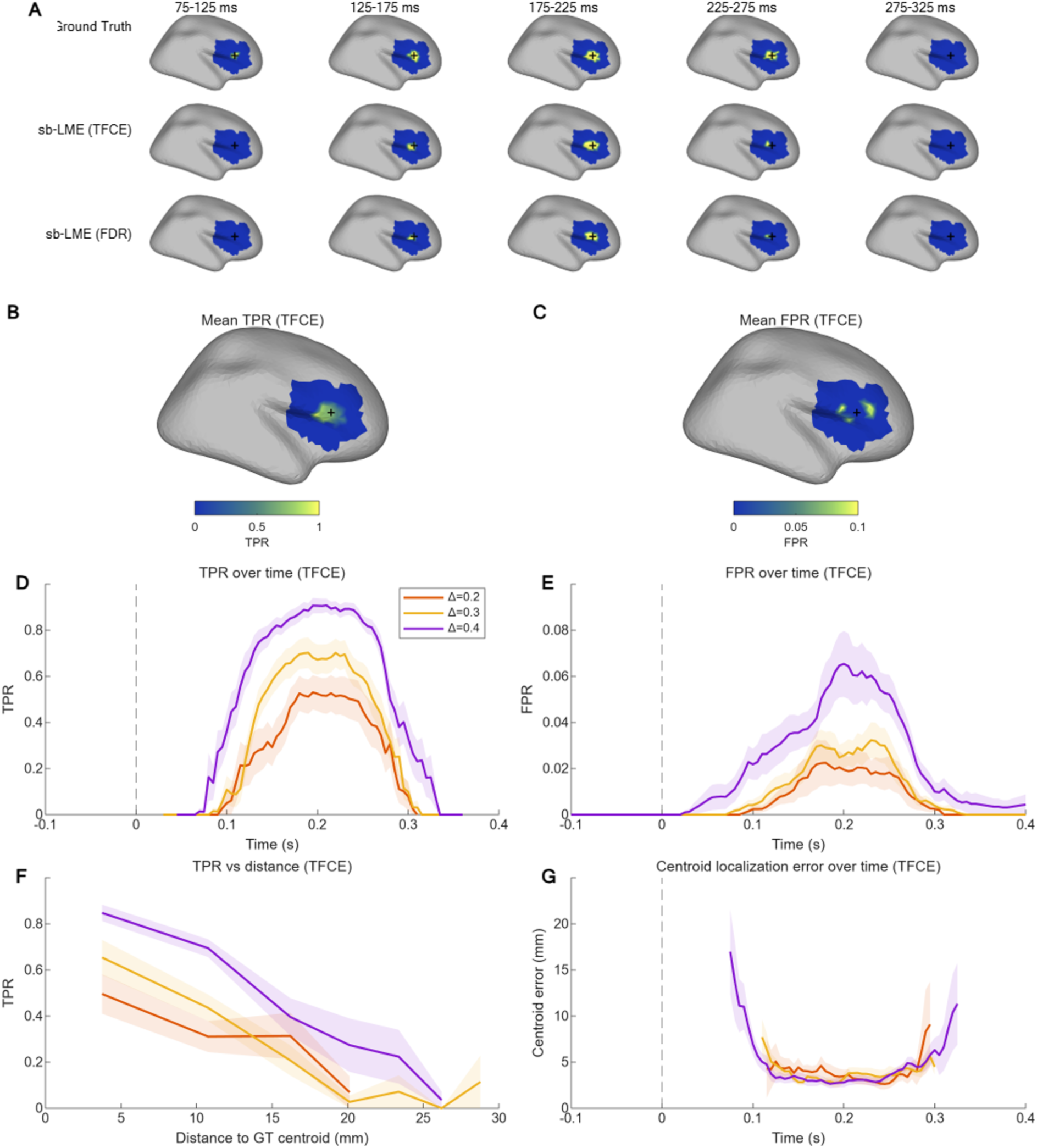
Temporal and spatial detection performance of STAMP-LME (sbLME) relative to ground truth (GT). A. Example detection maps for a simulated effect in the inferior frontal gyrus (IFG, Δ = 0.3 μV). GT TFCE-corrected detections (top row), STAMP-LME TFCE-corrected detections (middle row), and STAMP-LME FDR-corrected detections (bottom row) are shown for successive 50 ms windows. Only time windows spanning 100–300 ms are shown, as detections were absent outside this interval. The red cross denotes the GT source centroid. Both TFCE- and FDR-corrected STAMP-LME maps recovered the same spatiotemporal pattern of the simulated effect. Positive detection is color-coded yellow, and no detection is color-coded blue. Non-evaluated vertices beyond the sampled areas are depicted in gray color. B. Mean true positive rate (TPR) surface for the same IFG example (Δ = 0.3 μV), averaged across 0–400 ms using TFCE-corrected detections. The highest TPR values were concentrated near the GT centroid (red cross). C. Mean false positive rate (FPR) surface for the same IFG example (Δ = 0.3 μV), averaged across 0–400 ms using TFCE-corrected detections. FPR remained low and was primarily confined to the periphery of the source patch. D. TPR over time for different simulated effect sizes (Δ), averaged across all simulations. Shaded bands indicate ±SE. TPR tended to peak around the simulated response peak time (200 ms). E. FPR over time for different simulated effect sizes (Δ), averaged across all simulated datasets. Shaded bands indicate ±SE. F. TPR as a function of Euclidean distance from the GT centroid for different simulated effect sizes (Δ). Curves represent averages across all simulated datasets; shaded bands indicate ±SE. G. Centroid localization error over time for different simulated effect sizes (Δ). Shaded bands indicate ±SE. Note: Estimates for Δ = 0.1 μV were excluded because too few simulations produced detectable effects to obtain stable summary statistics.

Temporal detection performance was quantified using true positive rates and false positive rates. Higher TPR values were concentrated around the GT centroid (Fig. 7B), whereas FPR values were higher around the edge of the activation patch and dropped abruptly at 20 mm from the centroid (Fig. 7C and Supp. Fig. 3 G-H). From a temporal perspective, TPR increased sharply following stimulus onset and peaked within the simulated response interval (peak over 80% for Δ = 0.4, Fig. 7D), indicating reliable detection of the effect during the task-related period. FPR increased during the same period but remained low overall (average below 7%; Fig. 7E), demonstrating that STAMP-LME did not produce widespread spurious detections.

The choice of multiple-comparison correction affected detection sensitivity. TFCE correction consistently yielded higher TPR than FDR, reflecting increased sensitivity to spatially clustered effects, at the cost of a modest increase in FPR. In contrast, FDR produced more conservative detections with reduced sensitivity (Supp. Fig. 3).

Spatial specificity of effect detection was examined as a function of Euclidean distance from the GT activation centroid. TPRs were highest near the source location and decreased monotonically with increasing distance (Fig. 7F), indicating that STAMP-LME detections were spatially concentrated around the simulated effect.

Finally, spatial accuracy was quantified by measuring the Euclidean distance between the centroids of the STAMP-LME and GT detection clusters over time. Localization errors decreased rapidly after response onset and remained around 5 mm during peak activation (Fig. 7G), demonstrating accurate recovery of the dominant cortical source. It should be noted that the spatial resolution (i.e., average distance between vertices) of the STAMP-LME activation maps was approximately 5 mm.

#### 3.1.4. Factors influencing STAMP-LME detection performance

To assess how the simulation parameters affected STAMP-LME detection and recovery of effects, we evaluated simulation performance under a variety of scenarios and smoothing kernels. First, we tested whether effect size, electrode sampling, and source-location variability systematically influenced STAMP-LME performance.

Across outcomes, all three simulation factors significantly influenced most performance measures after correction for multiple comparisons. Increasing the simulated effect size improved detectability, TPR, and MCC. In contrast, reducing electrode coverage and increasing source-location jitter impaired these measures. Centroid localization error increased for increased source jitter. These effects were observed for both TFCE- and FDR-corrected results, although performance was generally better for TFCE in the weakest simulation scenarios. Full statistical summaries are shown in Supplementary Figures 4 and 5.

To assess the practical consequences of selecting the spatial smoothing kernel, σ = 3, 5, and 7 mm were compared directly within each scenario (Table 2 and Supp. Table 3 for TFCE- and FDR-corrected results respectively). Smoothing had little reliable influence on case-level detectability. For both TFCE- and FDR-corrected results, no pairwise difference in detectability remained significant in any simulated condition, indicating that σ did not consistently determine whether an STAMP-LME effect was detected. In contrast, metrics reflecting spatial overlap with the GT were more sensitive to the smoothing kernel. Precision generally favored smaller kernels, particularly σ = 3 mm, but 5 and 7 showed as equally valid alternatives in a few scenarios. MCC, a metric used to evaluate the quality of binary unbalanced classifications (Chicco & Jurman, 2023), showed a uniform pattern: σ = 3 and 5 mm often retained among the supported alternatives, whereas σ = 7 mm showed an equivalent performance in about 50% of the scenarios.

The most consistent σ effect was observed for spatiotemporal similarity (which is independent of the multiple comparison correction method). Across simulated conditions, σ = 3 mm produced higher similarity to the ground truth than σ = 5 or 7 mm, indicating that broader smoothing reduced the fidelity of the unthresholded STAMP-LME maps. Centroid localization error also tended to favor smaller kernels when computable, although FDR-corrected comparisons were less stable because difficult scenarios often produced no valid centroid estimate.

Overall, no single smoothing kernel optimized every performance measure. However, σ = 3 mm provided the most reliable spatial fidelity, with the highest similarity and generally higher precision, while σ = 5 mm was often retained as a supported alternative. The largest kernel, σ = 7 mm, provided no consistent benefit and was less often supported by metrics sensitive to spatial overlap or localization.

#### 3.1.5. STAMP-LME versus channel-based group analysis

To assess the practical sensitivity of STAMP-LME relative to conventional electrode-wise analysis, we compared detection performance between surface-based STAMP-LME inference and classical channel-level statistics applied independently at each electrode contact, followed by group-level testing.

Channel-level inference yielded scarce detections across simulation conditions. Even at the largest simulated effect size (Δ = 0.4 μV), only a small number of electrode contacts showed significant condition effects, with approximately three channels per subject on average (Fig. 9B). At lower effect sizes, detections were rare, often falling below one channel on average. Correspondingly, the proportion of subjects exhibiting any significant channel-level result increased with effect size but remained limited overall, ranging from approximately 10% to 60% depending on Δ (Fig. 9A).

**Figure 8.**
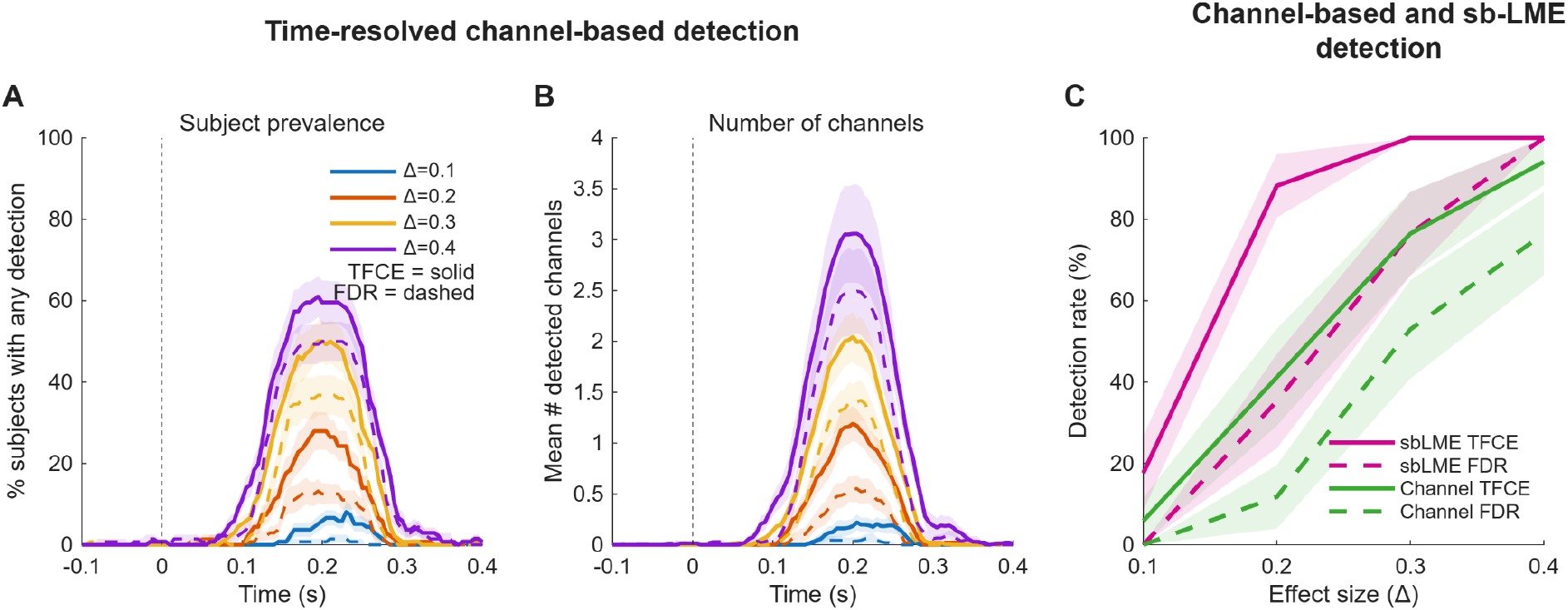
STAMP-LME (sbLME) versus channel-based group inference. A. Percentage of subjects with any significant channel-level detection for different effect sizes (Δ) plotted over time (s). Curves are averaged across all simulations. TFCE-corrected results are shown with solid lines and FDR-corrected results with dashed lines. Shaded bands denote ±SE. B. Mean number of significant channels detected with channel-level inference for different effect sizes (Δ) over time (s), averaged across simulations. TFCE-corrected results are shown with solid lines and FDR-corrected results with dashed lines. Shaded bands denote ±SE. C. Group-level detection rate (%) as a function of effect size (Δ), comparing channel-based group inference and STAMP-LME inference. Detection rate is defined as the percentage of observations classified as positive within the detection window across all simulations. Curves are shown separately for FDR- and TFCE-corrected channel-level inference (green) and for FDR- and TFCE-corrected STAMP-LME inference (pink). Shaded bands denote ±SE.

**Figure 9.**
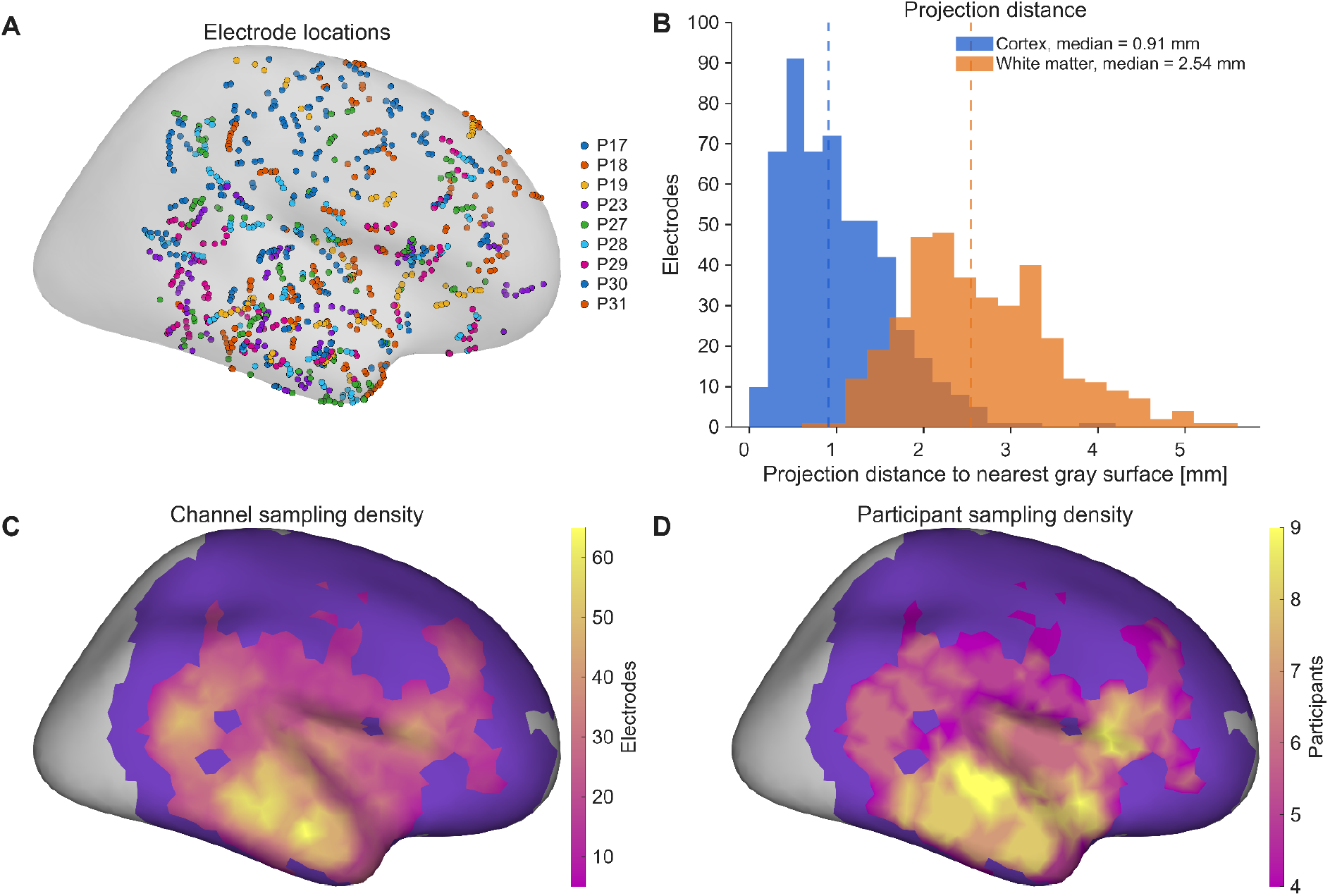
Electrode distribution, projection distance, and cortical sampling. A. Electrode locations projected onto the closest gray cortical surface, color-coded by participant (N = 9). B. Distribution of projection distance to the nearest gray cortical surface for electrodes within cortex (blue) and within white matter (orange). The dashed lines indicate the median values of each distribution. Gray surface was defined as the average of the pial and white surfaces. C. Channel sampling density, computed at each vertex as the number of electrodes within a 15 mm radius that contributed with more than 50% cumulative confidence. D. Participant sampling density, computed at each vertex as the number of unique participants contributing electrodes with more than 50% cumulative confidence within a 15 mm radius. Note: For (C) and (D), color-coded areas represent vertices where the minimum data criterion (i.e., minimum subject count *M_SC_*) was fulfilled, and STAMP-LMEs were evaluated. Purple regions were sampled, but did not fulfill the criterion. Gray areas were not sampled.

In contrast, STAMP-LME consistently achieved higher detection rates compared to channel-level based inferences across all effect sizes (Fig. 9C). This advantage was most pronounced at smaller effect sizes, where channel-based group analyses frequently failed to detect the simulated effect. By aggregating spatially distributed electrode information on the cortical surface and modeling variability across subjects within a unified framework, STAMP-LME improved sensitivity to distributed cortical sources that were only partially sampled by individual electrodes.

Across both STAMP-LME and channel-based group statistics, TFCE-based correction yielded higher detection rates than FDR. Importantly, STAMP-LME combined with TFCE achieved the highest overall detection rates while maintaining spatially localized detections and effectively leveraging the spatial structure of the reconstructed activity.

Together, these results demonstrate that under realistic iEEG sampling conditions, classical channel-based group-level testing provides limited sensitivity to distributed cortical effects, whereas STAMP-LME substantially improves detection performance by integrating spatial information across electrodes and subjects.

### 3.2. Application of STAMP-LME to empirical iEEG data

Having validated STAMP-LME under controlled simulation conditions, we next applied the framework to an empirical iEEG dataset to evaluate its practical use under realistic conditions. This analysis was not intended as a characterization of auditory deviance processing, but as a real-data demonstration of the full pipeline, accounting for electrode sampling, group-level inference, comparison with channel-level statistics, and sensitivity to the spatial smoothing parameter.

We analyzed intracranial EEG data from 9 participants, comprising 958 artifact-free channels (Supp. Table 1 summarizes patient demographics and implantation details). After excluding electrodes located in the hippocampus or amygdala, as well as those with a distance to the cortical surface greater than 4 mm (*T_Dist_*), 886 channels remained for analysis (Fig. 9A). The distribution of distances to the nearest gray cortical surface is shown in Fig. 9B for electrodes within the cortex and white matter. Figures 9C-D show the electrode and participant density at each vertex calculated within a 15 mm radius. Electrodes on the normalized left hemisphere were projected onto their homologous locations on the right-hemisphere pial surface. Participants contributed an average of 166 trials (range: 118–344), balanced between *IntUp* deviants and a randomly selected subset of *Std* trials (50% in each condition).

We then applied STAMP-LME to compare cortical responses to *IntUp* deviants and *Std* tones across vertices and time samples. Only vertices satisfying the minimum subject count criterion (*M_SC_*) were included. Figure 10A shows the mean Z-score STAMP-LME statistics for four consecutive post-stimulus time windows. The unthresholded activation maps revealed a structured spatiotemporal pattern, with effects emerging over the superior temporal plane and extending into posterior and anterior insular, frontal, and temporoparietal regions. Then, permutation-based FDR correction identified effects primarily along the superior temporal plane, posterior insula, temporoparietal junction, and minimally into the frontal cortex (Fig. 10B). TFCE correction yielded a similar spatiotemporal pattern (Fig. 10C).

**Figure 10.**
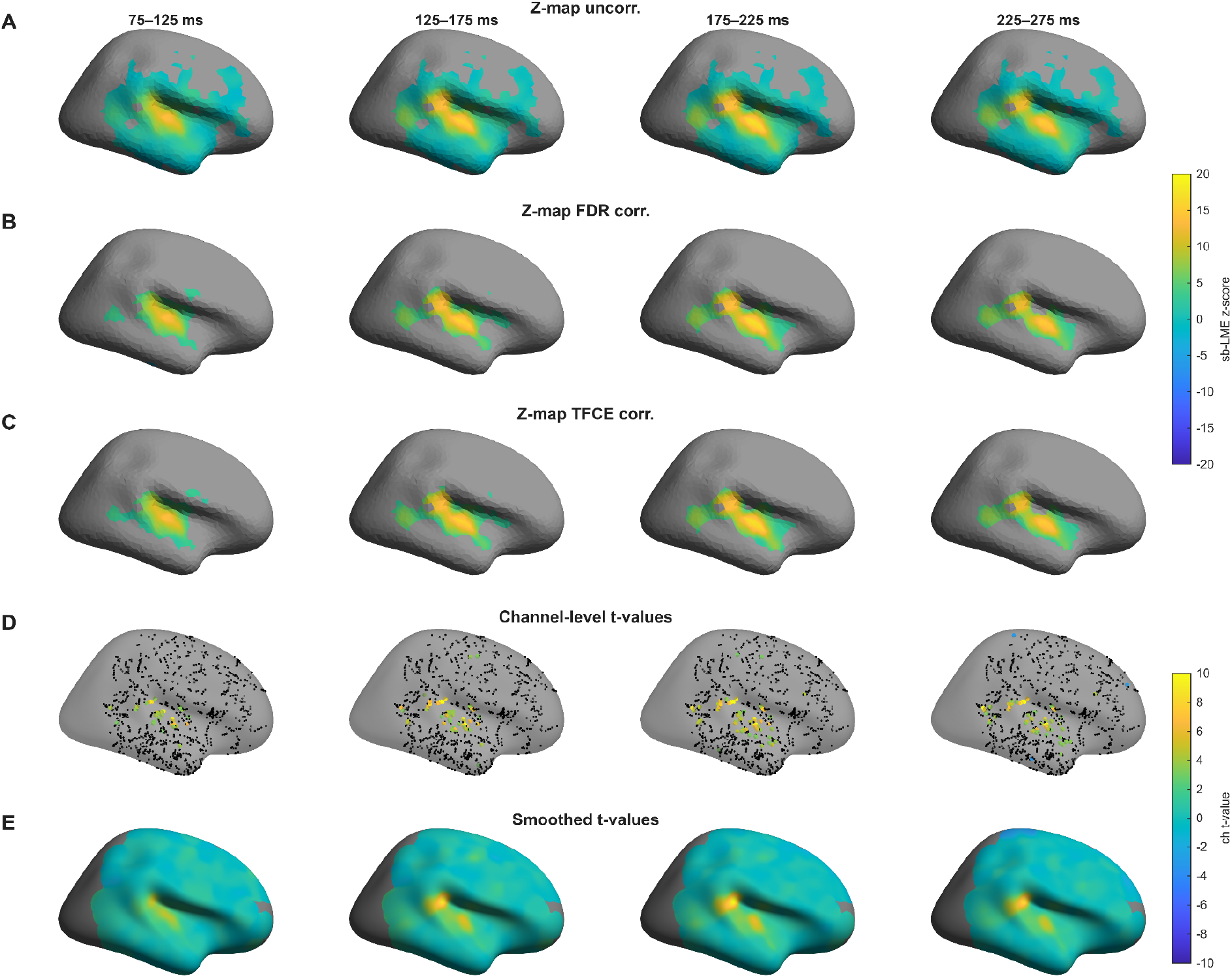
Z-scored STAMP-LME results and channel-level statistics for the IntUp versus Std contrast. Comparison of cortical response patterns obtained with STAMP-LME and channel-level analyses across four post-stimulus time windows. A. Mean Z-score STAMP-LME statistics for the IntUp versus Std contrast, shown for all vertices meeting the STAMP-LME inclusion criteria. Gray vertices indicate locations where STAMP-LME was not evaluated because the sampling criteria were not met. B. Same Z-score maps as in (A), masked by permutation-derived p-values after FDR correction for multiple comparisons. Vertices are color-coded only when significance was present for more than 50% of the samples within the corresponding time window; all other vertices are shown in gray. C. Same Z-score maps as in (A), masked using threshold-free cluster enhancement (TFCE)-corrected permutation statistics. D. Channel-level t-values for the IntUp versus Std contrast shown at individual electrode locations. Significant channels are color-coded, whereas non-significant channels are shown in black. Channels were projected to the nearest point on the cortical surface for visualization. E. Surface maps obtained by Gaussian smoothing of the channel-level t-values shown in (D) (σ = 5 mm). These maps reproduced the large-scale spatial organization seen in the STAMP-LME results, providing a qualitative comparison between channel-level and surface-based analyses. Unlike STAMP-LME, this visualization does not incorporate diffuse cortical projections or mixed-effects modeling.

Across both correction strategies, the strongest and most consistent activations were localized to superior temporal and adjacent perisylvian regions.

In addition, we evaluated the role of the smoothing parameter σ = 5 and 7 mm. The empirical observations closely mirrored the simulation results (section 3.1.4; Supp. Fig. 7), and highlight a trade-off between spatial precision and spatial coherence.

For comparison with a more conventional analysis, we computed channel-level t-values for the *Std* versus *IntUp* contrast. We projected them to the nearest cortical surface location (Fig. 10D). These channel-wise statistics were then averaged at each vertex using Gaussian spatial smoothing (σ = 5 mm; Fig. 10E), with participant-level results shown in Supp. Fig. 6. The resulting spatial patterns were broadly similar to those obtained with STAMP-LME, particularly along the superior temporal plane. However, unlike STAMP-LME, channel-level testing does not explicitly model inter-subject variability and does not incorporate diffuse mapping, distance penalization, or confidence weighting. Accordingly, these maps should be interpreted primarily as a qualitative reference and sanity check rather than as an equivalent group-level inferential framework.

Overall, the empirical application demonstrated that STAMP-LME recovered a coherent auditory deviance-related response pattern, dominated by superior temporal, posterior insular, and temporoparietal regions.

#### 3.2.1. Local neural dynamics extracted from STAMP-LME

The STAMP-LME framework enables characterization of neural response dynamics at specific cortical locations. To illustrate this capability, we extracted temporal profiles from two representative vertices: one in the primary auditory region (A1) and one in the planum temporale//temporoparietal junction (TPJ). For the A1 vertex, data were contributed by 36 channels across 6 participants (Fig. 11A). For the TPJ vertex, data were contributed by 29 channels across 7 participants (Fig. 11B).

**Figure 11.**
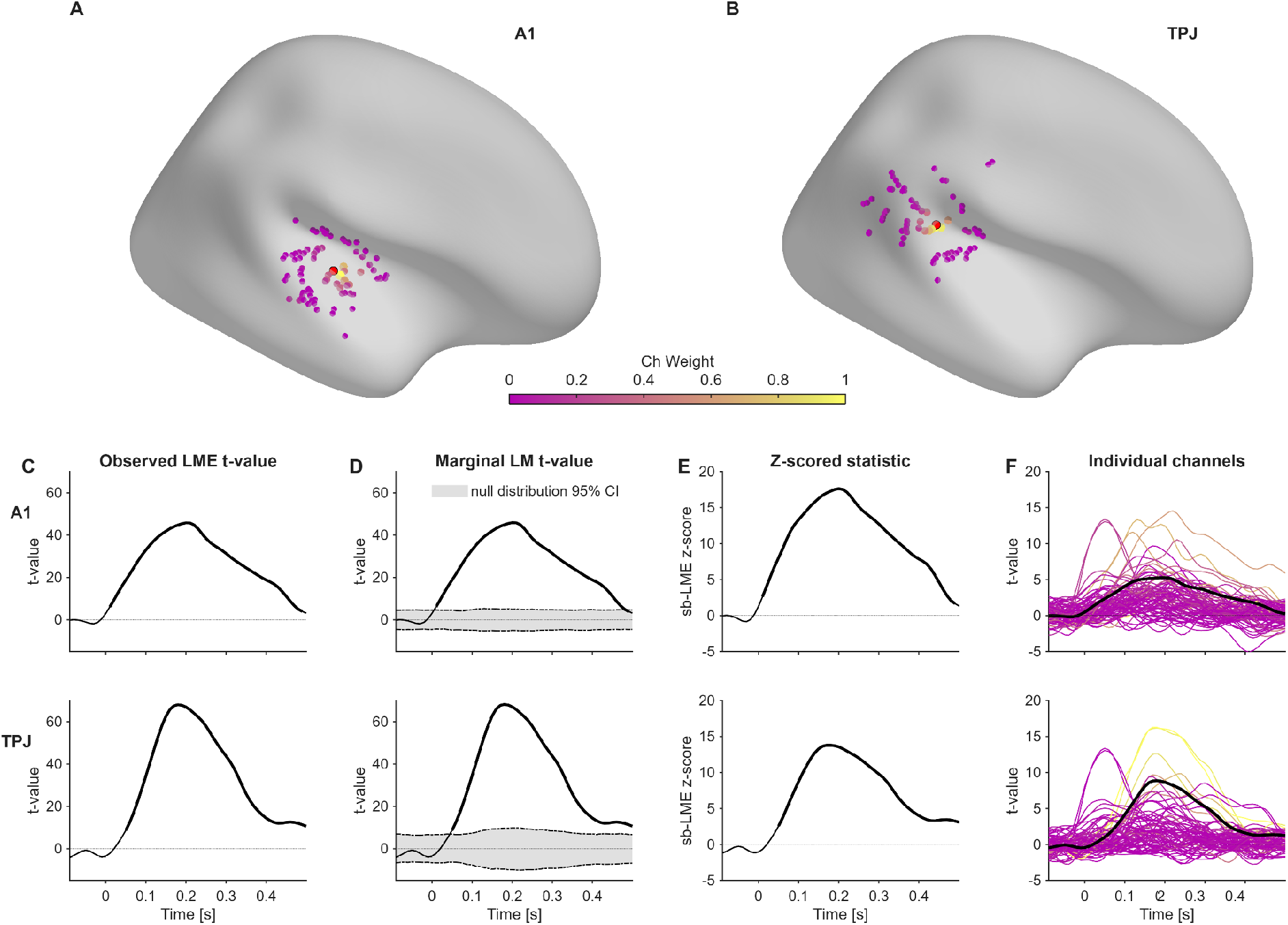
Local neural dynamics at representative A1 and TPJ vertices. A. Spatial distribution of channels contributing to the LME computation at the A1 vertex. Channel color indicates the channel weight in the LME. Red dots mark the selected vertex coordinate (local maximum activation A1 vertex). B. Same as (A), for the TPJ vertex. C. Observed LME-derived t-value time series for the IntUp versus Std contrast. These traces show the temporal evolution of the estimated vertex-level condition effect using the model. D. Marginal LM-derived t-value time series with the 99% interval of the permutation-derived null distribution. Deviations from the null interval indicate strong evidence of a task-condition effect. E. Z-score statistics computed from the observed marginal statistic and the permutation-derived null distribution. Significant periods (bold line) indicate time points surviving the corresponding permutation-based inference (TFCE-corrected). F. Individual channel-wise t-value time series for the electrodes contributing to each vertex, color-coded by their corresponding electrode-vertex weight, together with their weighted average (black line). These traces show the heterogeneity of single-channel HFA responses and their relationship to the vertex-level STAMP-LME estimate. The comparison illustrates that STAMP-LME captures a locally consistent response pattern while accounting for the hierarchical structure of the data.

Fig. 11C shows the LME-derived t-value time series (parametric approach). Fig. 11D displays the marginal LM-derived t-values together with the 99% confidence intervals from the null distribution obtained via permutation testing. Fig. 11E illustrates the Z-scored responses and corresponding periods of statistical significance. Finally, Fig. 11F shows the t-value time series for each contributing channel (based on independent t-tests across trials), along with their weighted average (an approach that neglects random effects but allows comparison of mixed-effects model results with a standard univariate test).

These examples illustrate how local temporal dynamics can be inferred at the cortical surface level while accounting for inter-subject variability and sparse electrode coverage. They also show how STAMP-LME results can be traced back to the underlying electrode contributions while preserving the high-temporal resolution of iEEG.

## 4. Discussion

### 4.1. STAMP-LME in the context of population-level iEEG analysis

Inferring population-level cortical effects from sparse and heterogeneous electrode coverage while retaining high spatial and temporal resolution is a central challenge of cognitive iEEG research. The STAMP-LME framework addresses this challenge by transforming irregular electrode-level observations into spatiotemporally resolved statistical maps on a standardized cortical surface. Its main contribution is the integration of cortical projection, spatial weighting, hierarchical modeling, and permutation inference into a common-space representation. Electrode-to-cortex distance and projection confidence determine how observations are mapped onto the cortical surface, while mixed-effects models account for repeated measurements and variability across participants and channels. STAMP-LME thereby estimates condition effects at each surface vertex and time point and enables data-driven analyses within sufficiently sampled cortical regions without requiring predefined ROIs.

ROI-based analyses remain a valuable approach for hypothesis-driven studies. The sparse and heterogeneous data from multiple participants are combined within predefined ROIs and use either group-level statistical models that account for both fixed effects and random effects (participant, electrode, session), or summary statistics approaches that computed a statistic for each participant and then combine these at the group-level testing (Lv & Wang, 2023; Mercier et al., 2022). Their strength depends on the relevant anatomical regions being defined a priori and independently of the observed effects (Axmacher, 2023; Combrisson et al., 2022; Magnotti et al., 2020; Mercier et al., 2022). However, well-defined a priori assumptions about the location, extent, and spatial boundaries of the regions are not always available, particularly when there is limited previous invasive evidence or when evidence comes from studies with less spatial precision (such as M/EEG source imaging or fNIRS). Under these circumstances, ROI definitions can influence statistical sensitivity: overly extensive ROIs may dilute focal effects, whereas narrower definitions may miss inter-individual spatial variability (König et al., 2024; Oehrn, 2023). Moreover, ROIs are typically defined by anatomical landmarks (Desikan et al., 2006; Destrieux et al., 2010), and may not correspond to functional subdivisions relevant for the particular experiment (but see (Glasser et al., 2016; Yeo et al., 2011)). Moreover, time-varying activations may cross ROI boundaries. Thus, although ROI-based analyses remain valuable, they can reduce one of the main advantages of iEEG, which is its ability to localize neural activity with high spatial precision. In contrast, STAMP-LME can estimate the spatial distribution of effects, as supported by the available sampling.

An alternative strategy is to perform ROI-free analyses within individual participants and subsequently assess the consistency of these effects across the population. For example, single-subject whole-brain or whole-implant analyses can be followed by population prevalence inference (Ince et al., 2021), or frequentist binomial testing (König et al., 2024; Mercier et al., 2022). Such approaches are useful for establishing whether an effect is reliably observed across individuals, even when electrode coverage is heterogeneous. However, consistency of detection does not by itself establish that the effects arise from a common anatomical location or time window. Follow-up analyses using statistically independent procedures may be required to support temporal and anatomical claims (Oehrn, 2023).

STAMP-LME builds on previous approaches that have shown that sparse intracranial recordings can be aligned to a common cortical surface and analyzed at the group level. Kadipasaoglu et al. (2014) introduced surface-based mixed-effects multilevel analysis (SB-MEMA) for ECoG. Electrode-level effect estimates were projected onto participants’ cortical surfaces, and then mapped into standardized surfaces where group-level inference was computed at each vertex. Trotta et al., (2018) followed with a similar approach, but instead aggregated electrodes within standardized reduced cortical parcels for group analysis. More recently, Woolnough et al. (2023) extended this strategy by constructing trial-by-trial participants’ maps and applying vertex-wise LME models for statistical inference (sbLME). These studies established the feasibility and value of transforming sparse intracranial recordings into population-level cortical maps.

STAMP-LME differs from these approaches in several fundamental respects. In previous implementations, the contributions of multiple electrodes were first combined into participant-level surface maps, which then served as the observations used by the group model. STAMP-LME instead retains channel-level observations within the hierarchical model and uses projection weights to determine their contribution at each vertex, allowing explicit representation of channel-level variability.

The projection procedure in STAMP-LME is also adapted to SEEG geometry. It distinguishes the initial mapping of a depth recording to plausible cortical locations from the subsequent group-level spatial integration of observations across the surface. Electrode-to-cortex distance and the relative confidence of candidate projection sites determine the cortical projection weights, whereas the group-level smoothing parameter determines the spatial scale over which mapped observations contribute to each vertex-level model. Separating these operations is particularly relevant for SEEG contacts, which may lie away from the cortical surface or plausibly sample activity from several nearby cortical locations (von Ellenrieder et al., 2012; 2021). To our knowledge none of the previous surface-based approaches implemented such a strategy to deal with ambiguous electrode positions.

The inferential procedures also differ. Earlier surface-based approaches used cluster-based correction based Monte Carlo simulations of spatially smoothed white noise (Kadipasaoglu et al. 2014; Woolnough et al., 2023), whereas STAMP-LME generates null statistical distributions through controlled permutations of the observed data (Maris & Oostenveld, 2007). Monte Carlo cluster correction assumes that synthetic spatially smoothed noise fields adequately approximate the distribution and dependence structure of the statistical maps under the null hypothesis. Direct comparisons between these approaches have shown that permutation-based inference provides more reliable control of false-positive rates (Eklund et al., 2016; Greve & Fischl, 2018). Permutation-based inference is nevertheless not assumption-free: its validity depends on exchangeability under the null hypothesis and on a permutation scheme that respects the relevant experimental dependencies (Maris & Oostenveld, 2007; Nichols & Holmes, 2002; Winkler et al., 2014). STAMP-LME therefore reduces reliance on an assumed synthetic null-field model by deriving its null distributions from the observed data under a permutation scheme matched to the experimental design. The controlled false-positive rates obtained in the present ground-truth simulations provide empirical support for this procedure.

The known-ground-truth simulations used in the current study provide an additional methodological contribution to validate the surface-based method developed here. Previous surface-based approaches were evaluated through their correspondence with individual electrode responses, expected functional anatomy, or alternative cortical mapping procedures (Kadipasaoglu et al. 2014; Trotta et al., 2018; Woolnough et al., 2023). Such comparisons establish empirical plausibility but cannot determine the true location, extent, or temporal profile of the underlying effect. Here, performance was evaluated directly against known cortical activation patterns, via simulations of cortical surface HFA, while varying effect size, interparticipant source-location variability, electrode coverage, and spatial smoothing. Quantifying detectability, sensitivity, precision, MCC, map similarity, and localization error allowed us to characterize not only whether STAMP-LME produced plausible maps, but also how accurately those maps recovered the simulated effects and how performance changed under less favorable source and sampling conditions.

Although the present validation focuses on SEEG, the general strategy should also be applicable to ECoG and mixed SEEG–ECoG datasets, provided that the electrode-to-cortex projection and weighting parameters are adapted and validated for the recording configuration. Finally, because the analysis operates on a normalized cortical surface, STAMP-LME facilitates comparison with surface-based fMRI and source-localized M/EEG results. The accompanying open-source pipeline (Figs. 1 and 2) further supports reproducibility and integration with commonly used neuroimaging and electrophysiology tools (e.g., FieldTrip, EEGLAB, FreeSurfer, iElectrodes), facilitating their incorporation into existing workflows.

### 4.2. Ground-truth simulations validate STAMP-LME spatiotemporal recovery of effects

The simulation framework provided a fully controlled ground-truth scenario for validating the proposed surface-based LME approach. Simulated sources were spatially localized yet extended over cortical patches, varied across participants, and followed temporally structured response profiles, thereby approximating key features of realistic task-related iEEG activity. Because the expected statistical outcome was clearly defined, this design permitted direct assessment of STAMP-LME at the vertex- and sample-level. Such validation is often unattainable in empirical iEEG datasets, where the true cortical origin, spatial extent, and temporal dynamics of task-related activity remain unknown.

The validation assessed recovery at two complementary levels. First, we computed map-level similarity (i.e., Pearson correlation) to quantify whether STAMP-LME preserved the continuous spatiotemporal structure of the simulated effect, independently of any statistical threshold. Second, detection-level metrics quantified whether thresholded inference identified the correct vertices and time points. This distinction is important because a method may produce statistical maps that resemble the ground truth but fail to withstand multiple-comparisons correction. Alternatively, a method may detect significant effects while only partially capturing their spatial extent or temporal profile.

Across simulations, STAMP-LME successfully recovered the main spatiotemporal structure of the ground-truth effects (Fig. 5A). The resulting Z-score maps tracked the expected temporal profile, with the strongest similarity during the simulated response window and weaker outside the activation period (Fig. 6C). Spatially, this similarity peaked within the simulated activation patch and declined with distance from the center (Figs. 6B and D). This pattern indicates that STAMP-LME preserved the spatial and temporal organization of the simulated effect. As expected, similarity depended on the signal-to-noise ratio, reaching its maximum near the response peak and within the activation patch, where the simulated cortical effect contributed most reliably to the observed channel-level signals relative to the additive channel noise (Fig. 5C). Moreover, the successful recovery of effects across different cortical regions demonstrates that the method’s accuracy was not restricted to a single favorable geometry (Supp. Figs. 1 and 2).

Detection-based metrics provided complementary evidence. True-positive rates increased with effect strength and were concentrated around the simulated activation sites (Figs. 7B, D and F; Supp. Fig. 3A, C, and E). Importantly, false-positive rates remained low (Fig. 7C and E and Supp. Fig. 3B, D, G, and H). Furthermore, centroid-based localization estimates showed that the center of mass of detected STAMP-LME effects remained close to the ground-truth activation centers (Fig. 7G and Supp. Fig. 3F), providing a direct measure of spatial localization accuracy.

The extended simulations showed that STAMP-LME performance varied systematically with the quality and spatial consistency of the simulated effects (Supp Figs. 4 and 5). As expected, weaker effect sizes, reduced electrode sampling, and greater source-location jitter decreased detectability. However, when effects were recovered, the reconstructed activation patterns remained broadly consistent with the GT maps, as reflected by precision, MCC, map-level similarity, and centroid localization error.

Interestingly, the smoothing parameter σ had a limited influence on overall performance (Table 2). Detectability changed little across the tested kernels, and most performance metrics showed only modest differences between σ = 3, 5, and 7 mm. With greater source-location variability, the differences between σ = 3 and 5 mm decreased, while σ = 7 mm performed comparably in some challenging scenarios but did not consistently provide an advantage. Mean centroid localization error remained low across conditions (< 8 mm), indicating that larger smoothing kernels did not introduce substantial systematic shifts in the estimated effect location.

The clearest effect of σ was observed for the similarity between GT and STAMP-LME activation maps. The smallest kernel, σ = 3 mm, consistently produced the highest similarity, suggesting that smaller spatial smoothing better preserved the structure of the simulated effects. This result should be interpreted in light of the fact that the GT maps were not spatially smoothed. Together, these findings indicate that STAMP-LME is robust to moderate changes in σ, with smoothing mainly affecting the spatial fidelity of the recovered maps rather than the presence or absence of statistical detection.

Overall, the convergence of map-level similarity, detection-level performance, and centroid localization supports the conclusion that STAMP-LME recovers the underlying spatiotemporal structure of simulated cortical HFA effects, while also supporting reliable statistical detection of activation patterns.

The comparison of STAMP-LME results with classical channel-based statistics further illustrates the value of moving statistical inference to the cortical surface. Channel-level analyses operate on a sparse and heterogeneous sampling space, making it difficult to combine information across individuals when electrodes are anatomically variable. In addition, statistical power is limited because effects must first be detected at individual channels before they can contribute to group-level conclusions (Fig. 8A-B). In contrast, STAMP-LME integrates spatially distributed electrode contributions onto a common cortical surface prior to group-level modeling, allowing partially overlapping evidence from different participants to contribute to the same cortical estimate. This resulted in substantially improved recovery of simulated effects, particularly under lower SNR conditions, where channel-level statistics failed to detect reliable group-level differences (Fig. 8C). This comparison indicates that, by combining diffuse electrode-to-surface projection into weighted mixed-effects modeling, STAMP-LME is better suited to detect spatially distributed group-level activations from sparse iEEG recordings.

To adjust for multiple comparisons, we used both pointwise FDR correction and TFCE correction. Overall, TFCE was more sensitive than FDR in the STAMP-LME simulations. This concurs with a previous study (Groppe et al., 2011) and likely occurs because TFCE exploits the spatiotemporal structure of neural activations rather than evaluating each vertex-time point independently. Consequently, TFCE is particularly well-suited for detecting spatially and temporally extended effects. However, FDR remains a useful complementary approach. Because it corrects individual vertex–time samples without relying on a spatially or temporally coherent cluster, FDR may be preferable when focal, spatially fragmented, or temporally brief effects are expected. This distinction is important because cluster-based approaches, including TFCE, can be less sensitive to small or fragmented effects and may favor patterns with stronger spatial or temporal continuity (Huang & Zhang, 2017). Additionally, TFCE results should not be interpreted as providing precise inference about the onset, duration, or exact anatomical boundaries of effects (Sassenhagen & Draschkow, 2019), whereas FDR results allow so.

The simulations also provided a stringent test because the forward model, used to generate SEEG synthetic recordings from projected cortical activity, and the STAMP-LME reconstructions were deliberately asymmetric, which makes the validation non-trivial. These two operations are not mathematically equivalent, and STAMP-LME does not simply invert the simulation model. Specifically, STAMP-LME incorporates confidence weighting and distance penalization, whereas the forward simulation utilized a different weighting scheme. Furthermore, the ground-truth solution had complete data for every cortical vertex and included random slopes to model inter-subject variability in condition effects. By contrast, STAMP-LME operated with missing data and heterogeneous electrode coverage across participants, and utilized a simpler random-effects structure, making the validation more challenging.

Therefore, the ability to recover the simulated effects despite these constraints validates the inference strategy rather than merely reflecting a trivial inversion of the simulation model and suggests that STAMP-LME is robust to missing information and inter-subject variability typical of group-level iEEG studies.

Taken together, the simulation results supported the spatial similarity and statistical validity of the framework. The STAMP-LME successfully recovered the location, timing, and extent of known cortical effects, maintained low false-positive rates, and outperformed conventional channel-level group statistics in the tested scenarios. These findings provide an essential validation step prior to applying the method to empirical data, where the true cortical activation pattern cannot be directly observed.

### 4.3. Empirical application of STAMP-LME to auditory deviance detection

The empirical SEEG analysis demonstrates the feasibility of applying STAMP-LME to real intracranial recordings with heterogeneous electrode coverage. In this auditory deviance dataset, STAMP-LME revealed distributed HFA responses across auditory, associative, and frontal cortical regions, while also making explicit which regions were sufficiently sampled for group-level inference (Fig. 10A). These effects are consistent with prior intracranial studies implicating auditory and perisylvian cortices in auditory deviance detection and predictive processing (Dürschmid et al., 2016; Edwards et al., 2005; El Karoui et al., 2015; Eliades et al., 2014; Phillips et al., 2016). The smaller frontal effects may reflect additional higher-order predictive processes (Deouell, 2007; Dürschmid et al., 2016; Phillips et al., 2016).

TFCE- and FDR-corrected results showed highly similar detection patterns (Fig. 10B-C). This convergence supports the robustness of the observed auditory deviance effects: the main activations were detectable both as pointwise vertex–time effects after FDR correction and as coherent spatiotemporal clusters after TFCE correction.

The variation of the smoothing parameter σ produced the expected blurring effect. We excluded the analysis for σ = 3 mm due to the reduced and fragmented sampled areas available. Kernels of σ = 5 mm and σ = 7 mm produced similar but smoother activation patterns for the latter, suggesting that both spatial integration is useful for empirical group-level SEEG analysis (Supp. Fig. 7). In addition, the single-subject surface projections revealed some level of heterogeneous activation patterns across participants further supporting this interpretation (Supp. Fig. 6). At the same time, the similarity between the unthresholded STAMP-LME Z-score map and the surface-smoothed channel-level t-map provides a useful sanity check: both approaches preserved the broad spatial organization of the empirical response (Fig. 10A and E). The advantage of STAMP-LME is that it extends this surface representation into a formal group-level statistical inference.

Finally, the vertex-level time courses in core auditory and associative regions provided an informative summary of the temporal dynamics underlying the surface maps (Fig. 11). These time courses helped link the spatial distribution of significant effects to the expected temporal profile of auditory deviance responses, illustrating how STAMP-LME can be used not only to localize group-level effects, but also to characterize their dynamic evolution over time.

### 4.4. Practical considerations for applying STAMP-LME

The simulation and empirical results indicate that STAMP-LME is robust across a range of analysis conditions, but a few practical choices remain important when applying the framework. These include the spatial smoothing parameter, the minimum sampling criteria required for vertex-wise inference, and the correction method used for multiple comparisons. Among these, the smoothing parameter σ is particularly important because it determines the spatial scale over which electrode-derived evidence is integrated across the cortical surface.

As a general starting point, we recommend using a Gaussian smoothing kernel of σ = 5 mm, corresponding to a FWHM = 11.77 mm. This value provides a practical compromise between local spatial specificity and sufficient spatial support for group-level inference. It is also broadly consistent with previous surface-based iEEG work. Kadipasaoglu et al. (2014) used a kernel with FWHM = 7.6 mm for group-level ECoG analysis, Miller et al. (2007) used σ = 5 mm to represent ECoG responses on cortical surfaces, and Esposito et al. (2013) reported optimal iEEG–fMRI correspondence for FWHM values around 10–15 mm. Importantly, σ should not be interpreted as the physiological “listening zone” of the electrodes, which depends on electrode type, frequency band, tissue properties, and referencing scheme (McCarty et al., 2022). Rather, σ is an analysis parameter that defines the spatial scale of statistical integration on the cortical surface.

The empirical application illustrated why this parameter should be adapted to the available sampling. In that dataset, σ = 5 mm provided a useful compromise, preserving focal temporal activations while maintaining spatial continuity. The smaller kernel, σ = 3 mm, produced maps that looked more sparse because the available spatial coverage was insufficient to support stable inference at that spatial scale. Conversely, σ = 7 mm produced results broadly similar to σ = 5 mm without a clear additional benefit.

The reliability of surface-based estimates also depends on the number of participants and electrodes that contribute meaningful information to each vertex. As emphasized by Mercier et al. (2022), there is no universal minimum sample size for iEEG group-level analysis, because statistical power depends on multiple interacting factors, including signal-to-noise ratio, effect size, spatial specificity, sampling density, and patient-related variability. In STAMP-LME, subject and electrode coverage thresholds are therefore study-dependent and interact with σ.

Therefore, although σ = 5 mm can serve as a useful default, the optimal value should be tailored to the expected spatial variability of the effect, the density and distribution of electrode sampling, and the physiological process under investigation. Larger smoothing kernels can increase the number of electrodes contributing to a vertex, but at the cost of spatial specificity. In contrast, smaller kernels may preserve more focal spatial patterns but can leave fewer vertices with sufficient group-level coverage. In practice, we recommend the smallest smoothing kernel that accommodates the expected inter-participant spatial variability to yield a sufficiently sampled surface for stable inference.

Consequently, the optimal combination of smoothing and coverage thresholds should be selected based on the goals and constraints of each study. Strong and spatially consistent effects, such as early sensory responses, may be detectable with more limited sampling, whereas weaker, more variable, or spatially distributed cognitive effects may require larger samples and stricter coverage criteria. Researchers should report the smoothing kernel, the minimum subject and electrode inclusion criteria, and the resulting sampled surface. Regions failing to meet coverage criteria should be interpreted as insufficiently sampled, rather than as evidence of an absence of neural effects.

### 4.5. Limitations and methodological scope

As with any group-level iEEG approach, STAMP-LME is constrained by the sparse and non-uniform sampling imposed by clinical electrode implantation (Johnson et al., 2020; Parvizi & Kastner, 2018). Electrode placement is determined by clinical needs rather than experimental design, resulting in uneven coverage across cortical and subcortical regions and variable electrode counts across participants. Although the mixed-effects formulation mitigates overrepresentation of densely sampled participants, regions with sparse coverage necessarily yield less reliable estimates (Mercier et al., 2022).

A second limitation concerns anatomical reconstruction and normalization. The framework relies on accurate cortical surface reconstruction, electrode localization, and registration to a common surface template. Segmentation errors, co-registration inaccuracies, or imperfect surface-based alignment can reduce the anatomical precision of group-level maps (Blenkmann et al., 2017). This issue is particularly relevant in patients with lesions, resections, or atypical cortical morphology, where surface standardization may be less reliable. Careful quality control and individualized correction procedures are therefore essential.

The present framework implements a simplified modeling strategy that does not explicitly incorporate tissue-specific differences in electrical conductivity between gray matter, white matter, and cerebrospinal fluid, nor the anisotropic characteristics of white matter (von Ellenrieder et al., 2012). These factors can influence volume conduction, especially near sulci, where cerebrospinal fluid and opposing cortical banks may shape the recorded field. To reduce ambiguity, our analyses focused on HFA activity from electrodes located within or near cortical gray matter, utilizing a projection-distance threshold to restrict contributions from distant generators (McCarty et al., 2022; von Ellenrieder et al., 2021). While this constrained approach optimizes interpretability for surface-based group analysis, it inherently limits the inclusion of more distant or distributed sources.

The framework was developed and validated using cortical HFA, a widely used iEEG measure associated with local population firing and postsynaptic activity (Lei et al., 2026; Leszczyński et al., 2020; Ray & Maunsell, 2011). HFA also generally exhibits greater spatial specificity than lower-frequency electrophysiological activity (McCarty et al., 2022; Parvizi and Kastner, 2017; Mercier et al., 2022; Stolk et al., 2018; Lachaux et al., 2012; Johnson et al., 2020). The framework is expected to extend naturally to other frequency-band power measures, although its performance may require further evaluation for electrophysiological features with different spatial characteristics or polarity properties. This issue is particularly relevant for event-related potentials, whose polarity depends on the orientation of the underlying current generators relative to electrode position and the referencing scheme. Individual anatomical variability may therefore produce differences in response polarity and partial cancellation at the group level. Developing approaches that accommodate interindividual anatomical variability while preserving ERP polarity therefore remains an important challenge for future work.

The STAMP-LME framework can handle multiple main effects, but it does not accommodate complex fixed-effect structures, such as interaction effects. Although model validation was limited to random intercepts (to reduce computational load and guarantee model convergence), this does not rule out the possibility of incorporating random slopes into the structure.

Finally, while our simulations were designed to test realistic signal-to-noise and spatial configurations, they were not exhaustive. The computational cost of the surface-based permutation framework was substantial, which limited the number of parameter combinations that could be explored. Future validation could systematically sample a broader range of source spatiotemporal patterns, noise structures, random-effects configurations, and electrode sampling patterns. Similarly, although TFCE provides a useful threshold-free approach for detecting spatially or temporally extended effects, we did not optimize its parameters for iEEG data. The present analyses used values previously applied in EEG contexts, and future work should assess whether alternative TFCE settings are better suited to STAMP-LME inference.

### 4.6. Outlook and future extensions

The present study focused on cortical HFA responses, but the STAMP-LME framework is not restricted to this signal feature. The same general approach could be applied to time series from distinct frequency bands, such as alpha- or beta-band activity, or to single-trial measures derived from information-theoretic analyses (Fuhrer et al., 2022, 2023; Combrison et al., 2022). Such extensions will likely require signal-specific choices of projection distance and smoothing scale, given that different electrophysiological features may differ in spatial spread.

Although our focus here was primarily on cortical mapping, the approach can be extended to subcortical structures. This could involve applying LME models and permutation testing to anatomically defined ROIs. Moreover, there is potential in mapping complex structures, such as the hippocampus, into unfolded 2D representations (DeKraker et al., 2018). Integrating these representations with STAMP-LME would provide a unified analytical framework across both cortical and subcortical territories.

Future developments could also incorporate more detailed electromagnetic modeling. Realistic forward models, for example, based on finite-element methods (Dessert et al., 2023; Zhang et al., 2008), could estimate subject-specific current distributions while accounting for tissue-specific conductivities and complex sulcal geometry. These subject-level estimates could then be entered into the group-level STAMP-LME framework, potentially reducing spatial ambiguities and improving the treatment of distant or cross-sulcal sources.

Finally, a natural extension of the proposed framework is the evaluation of interaction effects. This can be achieved using permutation schemes based on reduced models, where residuals are permuted while preserving main effects (Anderson, 2001; König et al., 2024).

Incorporating such approaches would enable the investigation of higher-order dependencies in cortical responses and should be addressed in future work.

## Glossary of terms

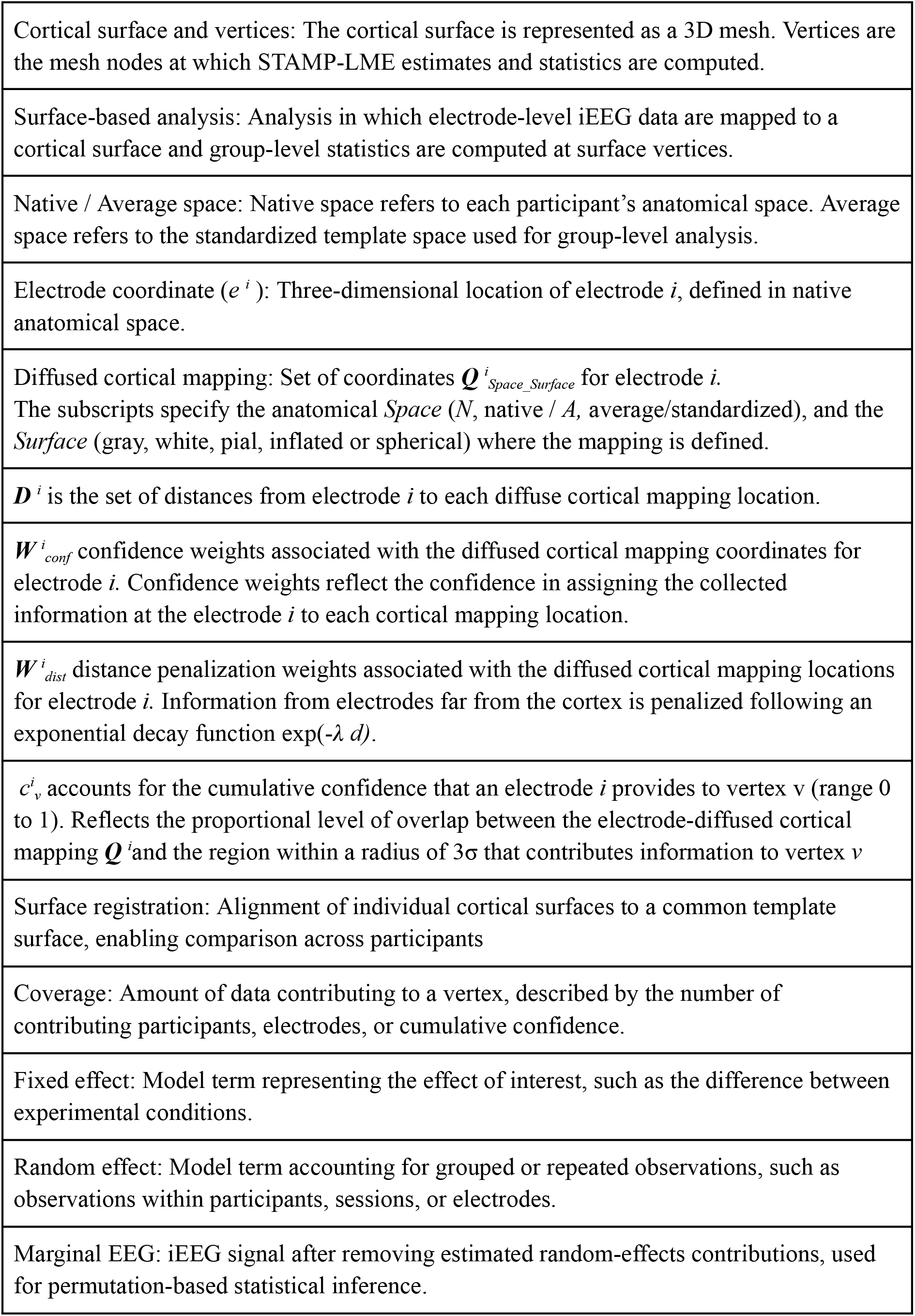
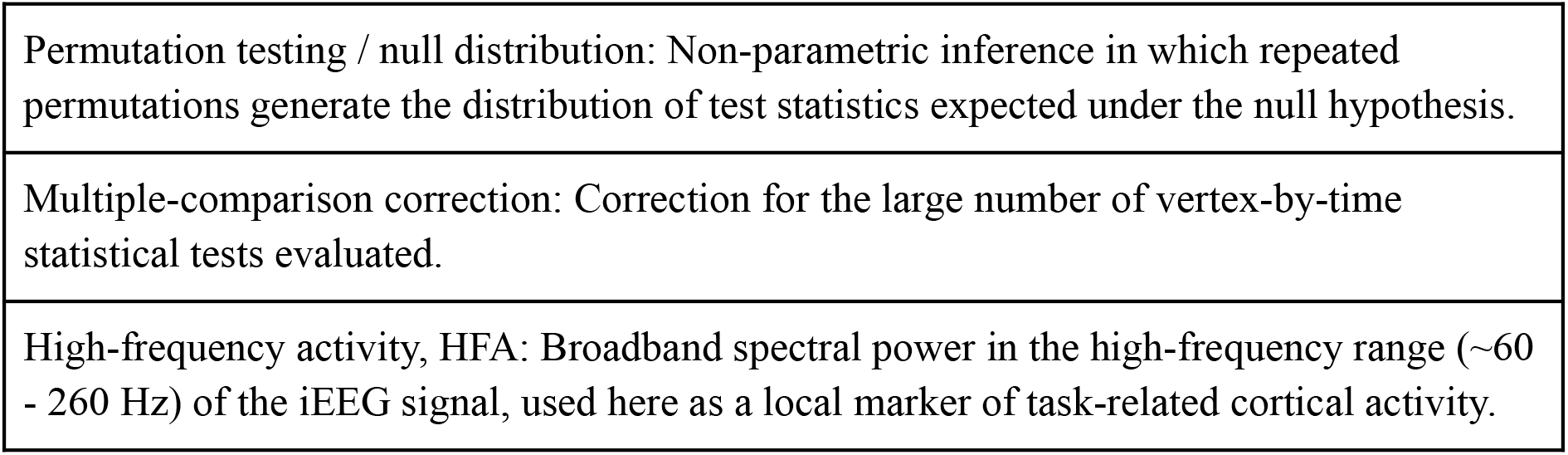

## 5. Data and Code availability

For transparency and reproducibility, the scripts used to generate synthetic data, along with the analysis scripts, the synthetic data, reconstructions, and statistical results from the STAMP-LME analysis are available in open repositories upon publication.

Due to the confidential nature of the patient data analyzed in this study, these data cannot be made publicly available. Our ethical approval conditions do not permit the public archiving of this study’s data. Readers seeking access to the data should contact the author, A. O. Blenkmann, Department of Psychology, University of Oslo.

## 6. Acknowledgements

We thank the patients for their participation in our study. We also express our gratitude to the clinicians and technicians at Oslo University Hospital-Rikshospitalet for their support during data collection. This work was supported in part by the Research Council of Norway (RCN) through its Centres of Excellence scheme, RITMO (project number 262762), and by RCN project “Neurophysiological Mechanisms of Human Auditory Predictions: From population- to single neuron recordings” (AudioPred; project number 314925).

## 7. CRediT statement

A.O.B. conceptualized the framework and developed the code. All authors evaluated the methodology and validation results. A.O.B. wrote the first manuscript. All authors reviewed and edited the manuscript. A.O.B. and A.K.S. provided the funding.

## References

Abou-Al-Shaar, H., Brock, A. A., Kundu, B., Englot, D. J., & Rolston, J. D. (2018). Increased nationwide use of stereoencephalography for intracranial epilepsy electroencephalography recordings. Journal of Clinical Neuroscience, 53, 132–134. 10.1016/j.jocn.2018.04.064

Allen, E. A., Erhardt, E. B., Wei, Y., Eichele, T., & Calhoun, V. D. (2012). Capturing inter-subject variability with group independent component analysis of fMRI data: A simulation study. NeuroImage, 59(4), 4141–4159. 10.1016/j.neuroimage.2011.10.010

Benjamini, Y., & Hochberg, Y. (1995a). Controlling the False Discovery Rate: A Practical and Powerful Approach to Multiple Testing. Journal of the Royal Statistical Society. Series B (Methodological), 57(1), 289–300.

Benjamini, Y., & Hochberg, Y. (1995b). Controlling the False Discovery Rate: A Practical and Powerful Approach to Multiple Testing. Journal of the Royal Statistical Society Series B: Statistical Methodology, 57(1), 289–300. 10.1111/j.2517-6161.1995.tb02031.x

Benjamini, Y., & Yekutieli, D. (2001). The control of the false discovery rate in multiple testing under dependency. The Annals of Statistics, 29(4). 10.1214/aos/1013699998

Blenkmann, A. (2012). *Localización de fuentes de actividad cerebral* [Doctor en Ingeniería, Universidad Nacional de La Plata]. 10.35537/10915/18159

Blenkmann, A., Beltrachini, L., von Ellenrieder, N., Kochen, S., & Muravchik, C. H. (2011). Evaluación de sLORETA para la resolución del problema inverso en ECoG utilizando modelos de poblaciones neuronales fisiológicamente plausibles. Reunión de Trabajo En Procesamiento de La Información y Control, RPIC2011.

Blenkmann, A. O., Collavini, S., Lubell, J., Llorens, A., Funderud, I., Ivanovic, J., Larsson, P. G., Meling, T. R., Bekinschtein, T., Kochen, S., Endestad, T., Knight, R. T., & Solbakk, A.-K. (2019). Auditory deviance detection in the human insula: An intracranial EEG study. Cortex, 121, 189–200. 10.1016/j.cortex.2019.09.002

Blenkmann, A. O., Leske, S. L., Llorens, A., Lin, J. J., Chang, E. F., Brunner, P., Schalk, G., Ivanovic, J., Larsson, P. G., Knight, R. T., Endestad, T., & Solbakk, A.-K. (2024). Anatomical registration of intracranial electrodes. Robust model-based localization and deformable smooth brain-shift compensation methods. Journal of Neuroscience Methods, 404, 110056. 10.1016/j.jneumeth.2024.110056

Blenkmann, A. O., Phillips, H. N., Princich, J. P., Rowe, J. B., Bekinschtein, T. A., Muravchik, C. H., & Kochen, S. (2017). iElectrodes: A Comprehensive Open-Source Toolbox for Depth and Subdural Grid Electrode Localization. Frontiers in Neuroinformatics, 11(March), 14. 10.3389/fninf.2017.00014

Blenkmann, A. O., Solbakk, A.-K., Ivanovic, J., Larsson, P. G., Knight, R. T., & Endestad, T. (2022). Modeling intracranial electrodes. A simulation platform for the evaluation of localization algorithms. Frontiers in Neuroinformatics, 16. 10.3389/fninf.2022.788685

Chicco, D., & Jurman, G. (2023). The Matthews correlation coefficient (MCC) should replace the ROC AUC as the standard metric for assessing binary classification. BioData Mining, 16(1), 4. 10.1186/s13040-023-00322-4

Coalson, T. S., Van Essen, D. C., & Glasser, M. F. (2018). The impact of traditional neuroimaging methods on the spatial localization of cortical areas. Proceedings of the National Academy of Sciences, 115(27), E6356–E6365. 10.1073/pnas.1801582115

Cohen, M. X. (2014). Analyzing Neural Time Series Data: Theory and Practice. MIT Press.

Combrisson, E., Allegra, M., Basanisi, R., Ince, R. A. A., Giordano, B. L., Bastin, J., & Brovelli, A. (2022). Group-level inference of information-based measures for the analyses of cognitive brain networks from neurophysiological data. NeuroImage, 258, 119347. 10.1016/j.neuroimage.2022.119347

Cosandier-Rimélé, D., Badier, J.-M., Chauvel, P., & Wendling, F. (2007). A physiologically plausible spatio-temporal model for EEG signals recorded with intracerebral electrodes in human partial epilepsy. IEEE Transactions on Bio-Medical Engineering, 54(3), 380–388. 10.1109/TBME.2006.890489

Cosandier-Rimélé, D., Merlet, I., Badier, J. M., Chauvel, P., & Wendling, F. (2008a). The neuronal sources of EEG: Modeling of simultaneous scalp and intracerebral recordings in epilepsy. NeuroImage, 42(1), 135–146. 10.1016/j.neuroimage.2008.04.185

Cosandier-Rimélé, D., Merlet, I., Badier, J. M., Chauvel, P., & Wendling, F. (2008b). The neuronal sources of EEG: modeling of simultaneous scalp and intracerebral recordings in epilepsy. NeuroImage, 42(1), 135–146. 10.1016/j.neuroimage.2008.04.185

Dale, A. M., Fischl, B., & Sereno, M. I. (1999). Cortical surface-based analysis. I. Segmentation and surface reconstruction. NeuroImage, 9(2), 179–194. 10.1006/nimg.1998.0395

Dasgupta, D., Miserocchi, A., McEvoy, A. W., & Duncan, J. S. (2022). Previous, current, and future stereotactic EEG techniques for localising epileptic foci. Expert Review of Medical Devices, 19(7), 571–580. 10.1080/17434440.2022.2114830

DeKraker, J., Ferko, K. M., Lau, J. C., Köhler, S., & Khan, A. R. (2018). Unfolding the hippocampus: An intrinsic coordinate system for subfield segmentations and quantitative mapping. NeuroImage, 167, 408–418. 10.1016/j.neuroimage.2017.11.054

Deouell, L. Y. (2007). The frontal generator of the mismatch negativity revisited. Journal of Psychophysiology, 21(3–4), 188–203. 10.1027/0269-8803.21.34.188

Depuydt, E., Criel, Y., De Letter, M., & van Mierlo, P. (2024). Investigating the effect of template head models on Event-Related Potential source localization: A simulation and real-data study. Frontiers in Neuroscience, 18. 10.3389/fnins.2024.1443752

Desikan, R. S., Ségonne, F., Fischl, B., Quinn, B. T., Dickerson, B. C., Blacker, D., Buckner, R. L., Dale, A. M., Maguire, R. P., Hyman, B. T., Albert, M. S., & Killiany, R. J. (2006). An automated labeling system for subdividing the human cerebral cortex on MRI scans into gyral based regions of interest. NeuroImage, 31(3), 968–980. 10.1016/j.neuroimage.2006.01.021

Dessert, G. E., Thio, B. J., & Grill, W. M. (2023). Optimization of patient-specific stereo-EEG recording sensitivity. Brain Communications, 5(6), fcad304. 10.1093/braincomms/fcad304

Destrieux, C., Fischl, B., Dale, A., & Halgren, E. (2010). Automatic parcellation of human cortical gyri and sulci using standard anatomical nomenclature. NeuroImage, 53(1), 1–15. 10.1016/j.neuroimage.2010.06.010

Dubarry, A.-S., Liégeois-Chauvel, C., Trébuchon, A., Bénar, C., & Alario, F.-X. (2022). An open-source toolbox for Multi-patient Intracranial EEG Analysis (MIA). NeuroImage, 257, 119251. 10.1016/j.neuroimage.2022.119251

Dürschmid, S., Edwards, E., Reichert, C., Dewar, C., Hinrichs, H., Heinze, H.-J., Kirsch, H. E., Dalal, S. S., Deouell, L. Y., & Knight, R. T. (2016). Hierarchy of prediction errors for auditory events in human temporal and frontal cortex. Proceedings of the National Academy of Sciences, 113(24), 201525030. 10.1073/pnas.1525030113

Dürschmid, S., Reichert, C., Hinrichs, H., Heinze, H.-J., Kirsch, H. E., Knight, R. T., & Deouell, L. Y. (2018). Direct Evidence for Prediction Signals in Frontal Cortex Independent of Prediction Error. Cerebral Cortex (New York, N.Y. : 1991), 1–9. 10.1093/cercor/bhy331

Edwards, E., Soltani, M., Deouell, L. Y., Berger, M. S., & Knight, R. T. (2005). High gamma activity in response to deviant auditory stimuli recorded directly from human cortex. Journal of Neurophysiology, 94(6), 4269–4280. 10.1152/jn.00324.2005

Eklund, A., Nichols, T. E., & Knutsson, H. (2016). Cluster failure: Why fMRI inferences for spatial extent have inflated false-positive rates. Proceedings of the National Academy of Sciences, 113(28), 7900–7905. 10.1073/pnas.1602413113

El Karoui, I., King, J.-R., Sitt, J., Meyniel, F., Van Gaal, S., Hasboun, D., Adam, C., Navarro, V., Baulac, M., Dehaene, S., Cohen, L., & Naccache, L. (2015). Event-Related Potential, Time-frequency, and Functional Connectivity Facets of Local and Global Auditory Novelty Processing: An Intracranial Study in Humans. Cerebral Cortex, 25(11), 4203–4212. 10.1093/cercor/bhu143

Eliades, S. J., Crone, N. E., Anderson, W. S., Ramadoss, D., Lenz, F. A., & Boatman-Reich, D. (2014). Adaptation of high-gamma responses in human auditory association cortex. Journal of Neurophysiology, 112(9), 2147–2163. 10.1152/jn.00207.2014

Erhardt, E. B., Allen, E. A., Wei, Y., Eichele, T., & Calhoun, V. D. (2012). SimTB, a simulation toolbox for fMRI data under a model of spatiotemporal separability. NeuroImage, 59(4), 4160–4167. 10.1016/j.neuroimage.2011.11.088

Esposito, F., Singer, N., Podlipsky, I., Fried, I., Hendler, T., & Goebel, R. (2013). Cortex-based inter-subject analysis of iEEG and fMRI data sets: Application to sustained task-related BOLD and gamma responses. NeuroImage, 66, 457–468. 10.1016/j.neuroimage.2012.10.080

Fang, Q., & Boas, D. A. (2009). Tetrahedral mesh generation from volumetric binary and grayscale images. Proceedings - 2009 IEEE International Symposium on Biomedical Imaging: From Nano to Macro, ISBI 2009, (Isbi), 1142–1145. 10.1109/ISBI.2009.5193259

Fischl, B. (2012). FreeSurfer. NeuroImage, 20 YEARS OF fMRI, 62(2), 774–781. 10.1016/j.neuroimage.2012.01.021

Fischl, B., Sereno, M. I., & Dale, A. M. (1999). Cortical surface-based analysis. II: Inflation, flattening, and a surface-based coordinate system. NeuroImage, 9(2), 195–207. 10.1006/nimg.1998.0396

Fischl, B., Sereno, M. I., Tootell, R. B. H., & Dale, A. M. (1999). High-resolution intersubject averaging and a coordinate system for the cortical surface. Human Brain Mapping, 8(4), 272–284. 10.1002/(SICI)1097-0193(1999)8:4%3C272::AID-HBM10%3E3.0.CO;2-4

Flinker, A., Chang, E. F., Kirsch, H. E., Barbaro, N. M., Crone, N. E., & Knight, R. T. (2010). Single-Trial Speech Suppression of Auditory Cortex Activity in Humans. Journal of Neuroscience, 30(49), 16643–16650. 10.1523/JNEUROSCI.1809-10.2010

Friston, K., Ashburner, J., Kiebel, S., Nichols, T., & Penny, W. (Eds.). (2007). Statistical Parametric Mapping. Academic Press. 10.1016/B978-0-12-372560-8.X5000-1

Fuhrer, J., Blenkmann, A., Endestad, T., Solbakk, A. K., & Glette, K. (2022). Complexity-based Encoded Information Quantification in Neurophysiological Recordings. Proceedings of the Annual International Conference of the IEEE Engineering in Medicine and Biology Society, EMBS, 2022-July(Mi), 2319–2323. 10.1109/EMBC48229.2022.9871501

Fuhrer, J., Glette, K., Ivanovic, J., Larsson, P. G., Bekinschtein, T., Kochen, S., Knight, R. T., Tørresen, J., Solbakk, A.-K., Endestad, T., & Blenkmann, A. (2025). Direct brain recordings reveal implicit encoding of structure in random auditory streams. Scientific Reports, 15(1), 14725. 10.1038/s41598-025-98865-5

Fuhrer, J., Glette, K., Llorens, A., Endestad, T., Solbakk, A.-K., & Blenkmann, A. O. (2023). Quantifying evoked responses through information-theoretical measures. Frontiers in Neuroinformatics, 17. https://www.frontiersin.org/articles/10.3389/fninf.2023.1128866

Gao, R., Van den Brink, R. L., Pfeffer, T., & Voytek, B. (2020). Neuronal timescales are functionally dynamic and shaped by cortical microarchitecture. eLife, 9, 1–44. 10.7554/ELIFE.61277

Gelman, A., & Hill, J. (2006). Data Analysis Using Regression and Multilevel/Hierarchical Models (1st ed.). Cambridge University Press. 10.1017/CBO9780511790942

Genovese, C. R., Lazar, N. A., & Nichols, T. (2002). Thresholding of statistical maps in functional neuroimaging using the false discovery rate. NeuroImage, 15(4), 870–878. 10.1006/nimg.2001.1037

Giroud, J., Trébuchon, A., Mercier, M., Davis, M. H., & Morillon, B. (2024). The human auditory cortex concurrently tracks syllabic and phonemic timescales via acoustic spectral flux. Science Advances, 10(51), eado8915. 10.1126/sciadv.ado8915

Glasser, M. F., Coalson, T. S., Robinson, E. C., Hacker, C. D., Harwell, J., Yacoub, E., Ugurbil, K., Andersson, J., Beckmann, C. F., Jenkinson, M., Smith, S. M., & Van Essen, D. C. (2016). A multi-modal parcellation of human cerebral cortex. Nature, 536(7615), 171–178. 10.1038/nature18933

Greve, D. N., & Fischl, B. (2018). False positive rates in surface-based anatomical analysis. NeuroImage, 171, 6–14. 10.1016/j.neuroimage.2017.12.072

Groppe, D. M., Urbach, T. P., & Kutas, M. (2011). Mass univariate analysis of event-related brain potentials/fields II: Simulation studies. Psychophysiology, 48(12), 1726–1737. 10.1111/j.1469-8986.2011.01272.x

Helfrich, R. F., Fiebelkorn, I. C., Szczepanski, S. M., Lin, J. J., Parvizi, J., Knight, R. T., & Kastner, S. (2018). Neural Mechanisms of Sustained Attention Are Rhythmic. Neuron, 99(4), 854–865.e5. 10.1016/j.neuron.2018.07.032

Hoy, C. W., Quiroga-Martinez, D. R., Sandoval, E., King-Stephens, D., Laxer, K. D., Weber, P., Lin, J. J., & Knight, R. T. (2023). Asymmetric coding of reward prediction errors in human insula and dorsomedial prefrontal cortex. Nature Communications, 14(1), 8520. 10.1038/s41467-023-44248-1

Huang, G., & Zhang, Z. (2017). Improving sensitivity of cluster-based permutation test for EEG/MEG data. 2017 8th International IEEE/EMBS Conference on Neural Engineering (NER), 9–12. 10.1109/NER.2017.8008279

Ince, R. A., Paton, A. T., Kay, J. W., & Schyns, P. G. (2021). Bayesian inference of population prevalence. eLife, 10, e62461. 10.7554/eLife.62461

Johnson, E. L., Kam, J. W. Y., Tzovara, A., & Knight, R. T. (2020). Insights into human cognition from intracranial EEG: A review of audition, memory, internal cognition, and causality. Journal of Neural Engineering, 17(5), 051001. 10.1088/1741-2552/ABB7A5

Joswig, H., Steven, D. A., Parrent, A. G., MacDougall, K. W., Mirsattari, S. M., McLachlan, R. S., Diosy, D. C., & Burneo, J. G. (2018). Intracranial Electroencephalographic Monitoring: From Subdural to Depth Electrodes. Canadian Journal of Neurological Sciences, 45(3), 336–338. 10.1017/cjn.2018.4

Kadipasaoglu, C. M., Baboyan, V. G., Conner, C. R., Chen, G., Saad, Z. S., & Tandon, N. (2014). Surface-based mixed effects multilevel analysis of grouped human electrocorticography. NeuroImage, 101, 215–224. 10.1016/j.neuroimage.2014.07.006

Kam, J. W. Y., Szczepanski, S. M., Canolty, R. T., Flinker, A., Auguste, K. I., Crone, N. E., Kirsch, H. E., Kuperman, R. A., Lin, J. J., Parvizi, J., & Knight, R. T. (2018). Differential Sources for 2 Neural Signatures of Target Detection: An Electrocorticography Study. Cerebral Cortex, 28(1), 9–20. 10.1093/cercor/bhw343

Kleiner M, Brainard D, P. D. (2007). What’s new in Psychtoolbox-3? Perception, 36.

König, S. D., Safo, S., Miller, K., Herman, A. B., & Darrow, D. P. (2024). Flexible multi-step hypothesis testing of human ECoG data using cluster-based permutation tests with GLMEs. NeuroImage, 290, 120557. 10.1016/j.neuroimage.2024.120557

Krol, L. R., Pawlitzki, J., Lotte, F., Gramann, K., & Zander, T. O. (2018). SEREEGA: Simulating event-related EEG activity. Journal of Neuroscience Methods, 309, 13–24. 10.1016/j.jneumeth.2018.08.001

Lachaux, J.-P., Axmacher, N., Mormann, F., Halgren, E., & Crone, N. E. (2012). High-frequency neural activity and human cognition: Past, present and possible future of intracranial EEG research. Progress in Neurobiology, 98(3), 279–301. 10.1016/j.pneurobio.2012.06.008

Laird, N. M., & Ware, J. H. (1982). Random-effects models for longitudinal data. Biometrics, 38(4), 963–974.

Lei, T., Scheid, M. R., Flint, R. D., Glaser, J. I., & Slutzky, M. W. (2026). Active dissociation of intracortical spiking and high gamma activity. Nature, 1–7. 10.1038/s41586-026-10331-y

Leszczyński, M., Barczak, A., Kajikawa, Y., Ulbert, I., Falchier, A. Y., Tal, I., Haegens, S., Melloni, L., Knight, R. T., & Schroeder, C. E. (2020). Dissociation of broadband high-frequency activity and neuronal firing in the neocortex. Science Advances, 6(33), 977–989. 10.1126/sciadv.abb0977

Lv, P., & Wang, L. (2023). How Can iEEG Data Be Analyzed via Multi-Level Models? In N. Axmacher (Ed.), Intracranial EEG: A Guide for Cognitive Neuroscientists (pp. 579–586). Springer International Publishing. 10.1007/978-3-031-20910-9_36

Magnotti, J. F., Wang, Z., & Beauchamp, M. S. (2020). RAVE: Comprehensive open-source software for reproducible analysis and visualization of intracranial EEG data. NeuroImage, 223, 117341. 10.1016/j.neuroimage.2020.117341

Maris, E., & Oostenveld, R. (2007). Nonparametric statistical testing of EEG- and MEG-data. Journal of Neuroscience Methods, 164(1), 177–190. 10.1016/j.jneumeth.2007.03.024

McCarthy, G., Puce, a, Belger, a, & Allison, T. (1999). Electrophysiological studies of human face perception. II: Response properties of face-specific potentials generated in occipitotemporal cortex. Cerebral Cortex (New York, N.Y. : 1991), 9(5), 431–444.

McCarty, M. J., Woolnough, O., Mosher, J. C., Seymour, J., & Tandon, N. (2022). The Listening Zone of Human Electrocorticographic Field Potential Recordings. eNeuro, 9(2). 10.1523/ENEURO.0492-21.2022

Mensen, A., & Khatami, R. (2013). Advanced EEG analysis using threshold-free cluster-enhancement and non-parametric statistics. NeuroImage, 67, 111–118. 10.1016/j.neuroimage.2012.10.027

Mercier, M. R., Bickel, S., Megevand, P., Groppe, D. M., Schroeder, C. E., Mehta, A. D., & Lado, F. A. (2017). Evaluation of cortical local field potential diffusion in stereotactic electro-encephalography recordings: A glimpse on white matter signal. NeuroImage, 147, 219–232. 10.1016/j.neuroimage.2016.08.037

Mercier, M. R., Dubarry, A.-S., Tadel, F., Avanzini, P., Axmacher, N., Cellier, D., Vecchio, M. D., Hamilton, L. S., Hermes, D., Kahana, M. J., Knight, R. T., Llorens, A., Megevand, P., Melloni, L., Miller, K. J., Piai, V., Puce, A., Ramsey, N. F., Schwiedrzik, C. M., … Oostenveld, R. (2022). Advances in human intracranial electroencephalography research, guidelines and good practices. NeuroImage, 260, 119438. 10.1016/j.neuroimage.2022.119438

Miller, K. J., Leuthardt, E. C., Schalk, G., Rao, R. P. N., Anderson, N. R., Moran, D. W., Miller, J. W., & Ojemann, J. G. (2007). Spectral Changes in Cortical Surface Potentials during Motor Movement. Journal of Neuroscience, 27(9), 2424–2432. 10.1523/JNEUROSCI.3886-06.2007

Mukamel, R. (2005). Coupling Between Neuronal Firing, Field Potentials, and fMRI in Human Auditory Cortex. Science, 309(5736), 951–954. 10.1126/science.1110913

Näätänen, R., Pakarinen, S., Rinne, T., & Takegata, R. (2004). The mismatch negativity (MMN): Towards the optimal paradigm. Clinical Neurophysiology, 115, 140–144. 10.1016/j.clinph.2003.04.001

Nichols, T. E., & Holmes, A. P. (2002). Nonparametric permutation tests for functional neuroimaging: A primer with examples. Human Brain Mapping, 15(1), 1–25. 10.1002/hbm.1058

Oehrn, C. (2023). What Are the Pros and Cons of ROI Versus Whole-Brain Analysis of iEEG Data? In N. Axmacher (Ed.), Intracranial EEG (pp. 475–486). Springer International Publishing. 10.1007/978-3-031-20910-9_29

Oostenveld, R., Fries, P., Maris, E., & Schoffelen, J.-M. (2011). FieldTrip: Open Source Software for Advanced Analysis of MEG, EEG, and Invasive Electrophysiological Data. Computational Intelligence and Neuroscience, 2011, 1–9. 10.1155/2011/156869

Parvizi, J., & Kastner, S. (2017). Human intracranial EEG: Promises and Limitations. Nature Neuroscience. 10.1038/s41593-018-0108-2

Pernet, C. R., Latinus, M., Nichols, T. E., & Rousselet, G. A. (2015). Cluster-based computational methods for mass univariate analyses of event-related brain potentials/fields: A simulation study. Journal of Neuroscience Methods, Cutting-Edge EEG Methods, 250, 85–93. 10.1016/j.jneumeth.2014.08.003

Phillips, H. N., Blenkmann, A., Hughes, L. E., Kochen, S., Bekinschtein, T. A., Cam-CAN, & Rowe, J. B. (2016). Convergent evidence for hierarchical prediction networks from human electrocorticography and magnetoencephalography. Cortex, 82, 192–205. 10.1016/j.cortex.2016.05.001

Pinheiro, Bates. (2000). Mixed-Effects Models in S and S-PLUS. Springer-Verlag. 10.1007/b98882

Ray, S., & Maunsell, J. H. R. (2011). Different Origins of Gamma Rhythm and High-Gamma Activity in Macaque Visual Cortex. 9(4). 10.1371/journal.pbio.1000610

Sassenhagen, J., & Draschkow, D. (2019). Cluster-based permutation tests of MEG/EEG data do not establish significance of effect latency or location. Psychophysiology, 56(6), 1–8. 10.1111/psyp.13335

Scheer, H. J., Sander, T., & Trahms, L. (2006). The influence of amplifier, interface and biological noise on signal quality in high-resolution EEG recordings. Physiological Measurement, 27(2), 109–117. 10.1088/0967-3334/27/2/002

Smith, S. M., & Nichols, T. E. (2009). Threshold-free cluster enhancement: Addressing problems of smoothing, threshold dependence and localisation in cluster inference. NeuroImage, 44(1), 83–98. 10.1016/j.neuroimage.2008.03.061

Stolk, A., Griffin, S., Van Der Meij, R., Dewar, C., Saez, I., Lin, J. J., Piantoni, G., Schoffelen, J. M., Knight, R. T., & Oostenveld, R. (2018). Integrated analysis of anatomical and electrophysiological human intracranial data. Nature Protocols, 13(7), 1699–1723. 10.1038/s41596-018-0009-6

Trotta, M. S., Cocjin, J., Whitehead, E., Damera, S., Wittig, J. H., Saad, Z. S., Inati, S. K., & Zaghloul, K. A. (2018). Surface based electrode localization and standardized regions of interest for intracranial EEG. Human Brain Mapping, 39(2), 709–721. 10.1002/hbm.23876

Visalli, A., Montefinese, M., Viviani, G., Finos, L., Vallesi, A., & Ambrosini, E. (2024a). lmeEEG: Mass linear mixed-effects modeling of EEG data with crossed random effects. Journal of Neuroscience Methods, 401, 109991. 10.1016/j.jneumeth.2023.109991

Visalli, A., Montefinese, M., Viviani, G., Finos, L., Vallesi, A., & Ambrosini, E. (2024b). lmeEEG: Mass linear mixed-effects modeling of EEG data with crossed random effects. Journal of Neuroscience Methods, 401, 109991. 10.1016/j.jneumeth.2023.109991

von Ellenrieder, N., Beltrachini, L., & Muravchik, C. H. (2012). Electrode and brain modeling in stereo-EEG. Clinical Neurophysiology, 123(9), 1745–1754. 10.1016/j.clinph.2012.01.019

von Ellenrieder, N., Khoo, H. M., Dubeau, F., & Gotman, J. (2021). What do intracerebral electrodes measure? Clinical Neurophysiology, 132(5), 1105–1115. 10.1016/j.clinph.2021.02.012

Weber, J., Solbakk, A.-K., Blenkmann, A. O., Llorens, A., Funderud, I., Leske, S., Larsson, P. G., Ivanovic, J., Knight, R. T., Endestad, T., & Helfrich, R. F. (2024). Ramping dynamics and theta oscillations reflect dissociable signatures during rule-guided human behavior. Nature Communications, 15(1), 637. 10.1038/s41467-023-44571-7

Winkler, A. M., Ridgway, G. R., Webster, M. A., Smith, S. M., & Nichols, T. E. (2014). Permutation inference for the general linear model. NeuroImage, 92, 381–397. 10.1016/j.neuroimage.2014.01.060

Woolnough, O., Donos, C., Murphy, E., Rollo, P. S., Roccaforte, Z. J., Dehaene, S., & Tandon, N. (2023). Spatiotemporally distributed frontotemporal networks for sentence reading. Proceedings of the National Academy of Sciences, 120(17), e2300252120. 10.1073/pnas.2300252120

World Medical Association. (2025). World Medical Association Declaration of Helsinki: Ethical Principles for Medical Research Involving Human Participants. JAMA, 333(1), 71–74. 10.1001/jama.2024.21972

Yeo, B. T. T., Krienen, F. M., Sepulcre, J., Sabuncu, M. R., Lashkari, D., Hollinshead, M., Roffman, J. L., Smoller, J. W., Zöllei, L., Polimeni, J. R., Fischl, B., Liu, H., & Buckner, R. L. (2011). The organization of the human cerebral cortex estimated by intrinsic functional connectivity. Journal of Neurophysiology, 106(3), 1125–1165. 10.1152/jn.00338.2011

Zhang, Y., van Drongelen, W., Kohrman, M., & He, B. (2008). Three-dimensional brain current source reconstruction from intra-cranial ECoG recordings. NeuroImage, 42(2), 683–695. 10.1016/j.neuroimage.2008.04.263

