## supplementary material for "STAMP-LME: A spatiotemporal surface-mapping framework for population-level iEEG"

by

Alejandro O. Blenkmann, Sabine L. Leske , Robert T. Knight, Anne-Kristin Solbakk and Robert Oostenveld

| Patient | Age | Sex | Hand. | Epilepsy debut (y) | MRI Lesion | Implanted electrodes | Recording Type | Hemisph. | Epileptogenic zone |
|---|---|---|---|---|---|---|---|---|---|
| 1 | 27 | M | R | 24 | Left Wernicke | 101 | SEEG | Bilateral | Next to left primary sensory cortex, extending to the trigone |
| 2' | 34 | F | R | 11 | Left parietal, Tumor | 165 | SEEG | Bilateral | Left parietal |
| 3 | 35 | M | R | 26 | Right temp, unspec | 110 | SEEG | Right | Right temporal |
| 4 | 50 | F | R | 27 | Right hippocampal resection | 147 | SEEG | Bilateral | Left Hippocampus |
| 5 | 33 | F | R | 14 | no | 211 | SEEG | Right | Not identified, suspected right temp. pole |
| 6 | 39 | M | R | 16 | no | 212 | SEEG | Right | Ventral temporal |
| 7 | 41 | F | R | 28 | Right Hippocampal formation | 193 | SEEG | Right | Right posterior superior temporal sulcus |
| 8 | 18 | M | R | 6 | Right Fronto-Parieta l | 247 | SEEG | Right | |
| 9 | 50 | F | R | 18 | Left Ventral Parietal | 162 | SEEG | Left | Left ventral parietal |

**Supplementary Table 1.** Patients' demographics

| Region | Description | Centroid (x, y, z) | | |
|---|---|---|---|---|
| A1 | Primary Auditory Cortex | 25 | 0 | -13 |
| AIP | Anterior Intraparietal Sulcus | 14 | -34 | 41 |
| IFG | Inferior Frontal Gyrus | 24 | 55 | -7 |
| PSL | PeriSylvian Language Area | 39 | -26 | 6 |
| MFG | Middle Frontal Gyrus | 7 | 79 | 17 |
| 8Av | Area 8A ventral | 9 | 53 | 36 |
| STGl | Lateral Superior Temporal Gyrus | 38 | -4 | -26 |
| STGa | Anterior Superior Temporal Gyrus | 27 | 28 | -53 |
| Motor | Primary Motor Cortex | 3 | 5 | 58 |

**Supplementary Table 2.** Simulated task-related source coordinates and anatomical description. Source coordinates are described in Freesurfer volumetric coordinate space.

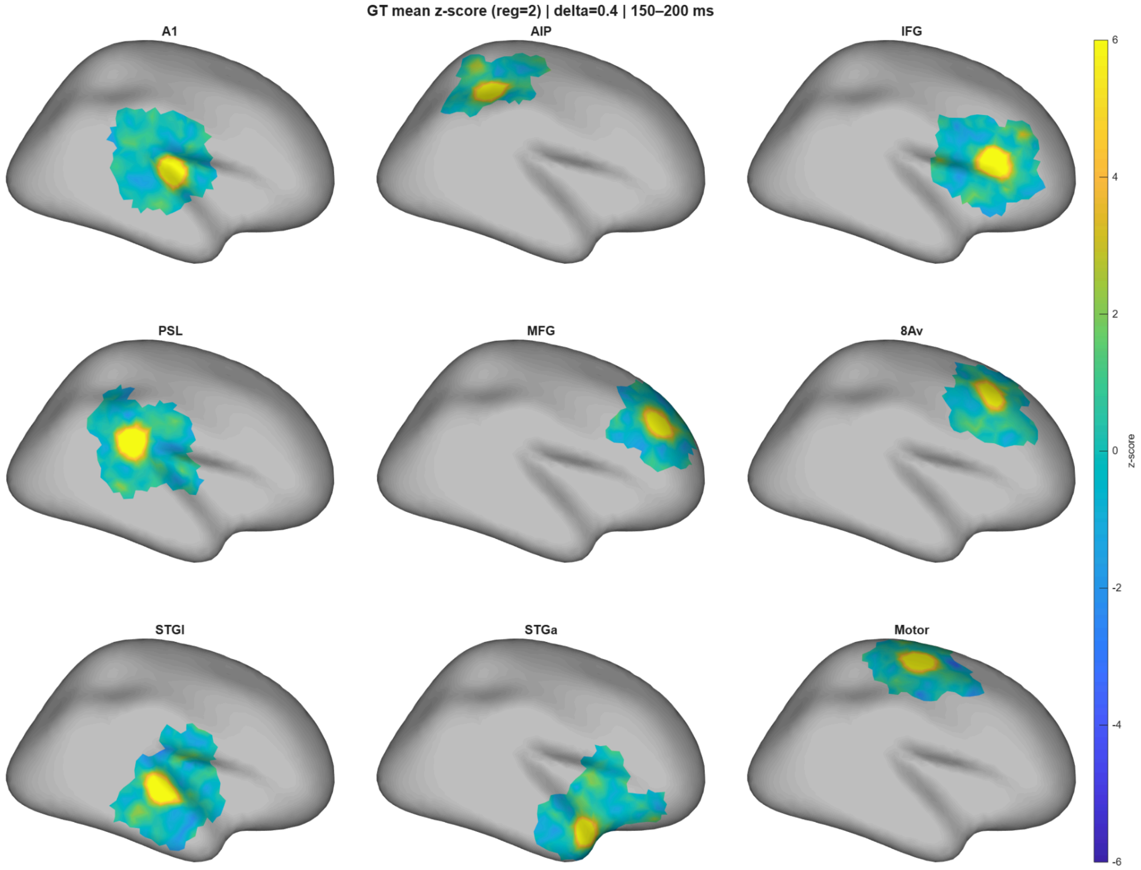


**Supplementary Figure 1. Ground-truth (GT) group-level effect maps across regions.**
For each region (A1, AIP, IFG, PSL, MFG, 8Av, STGl, STGa, Motor), the surface map shows the mean permutation-based z-score for the task-condition effect (B vs A) for effect size Δ = 0.4, averaged within the 150–200 ms post-stimulus window across participants. Z-scores were computed by standardizing the observed marginal t-statistic against the within-subject permutation null distribution at each vertex and time point. Z-scores were only computed for the vertices where STAMP-LME was later computed (supp Fig. 2). Non-sampled vertices are shown in gray color. The figure corresponds to the first simulation run. The analysis of the second run yielded similar results.

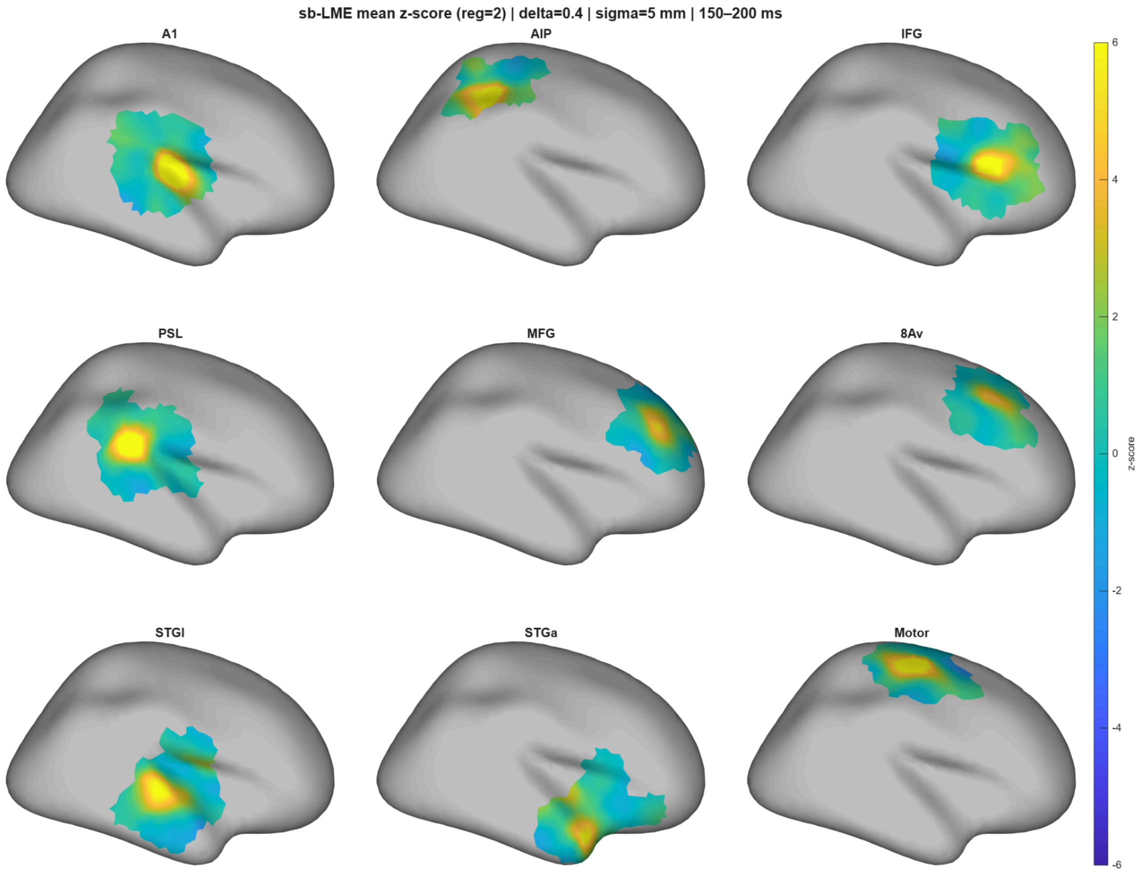


**Supplementary Figure 2. STAMP-LME group-level effect maps across regions.**
For each region of interest (A1, AIP, IFG, PSL, MFG, 8Av, STGl, STGa, Motor), the surface map shows the mean permutation-based z-score for the task-condition effect (B vs A) for effect size $\Delta = 0.4$, averaged within the 150–200 ms post-stimulus window. STAMP-LME was computed using a group-level spatial smoothing $\sigma = 5$ mm.
Z-scores were obtained by standardizing the observed marginal t-statistic at each vertex and time point against the corresponding within-subject permutation null distribution. Only vertices with valid electrode mapping are displayed; non-sampled vertices are shown in a gray color.
The figure corresponds to the first simulation run. The analysis of the second run yielded similar results.

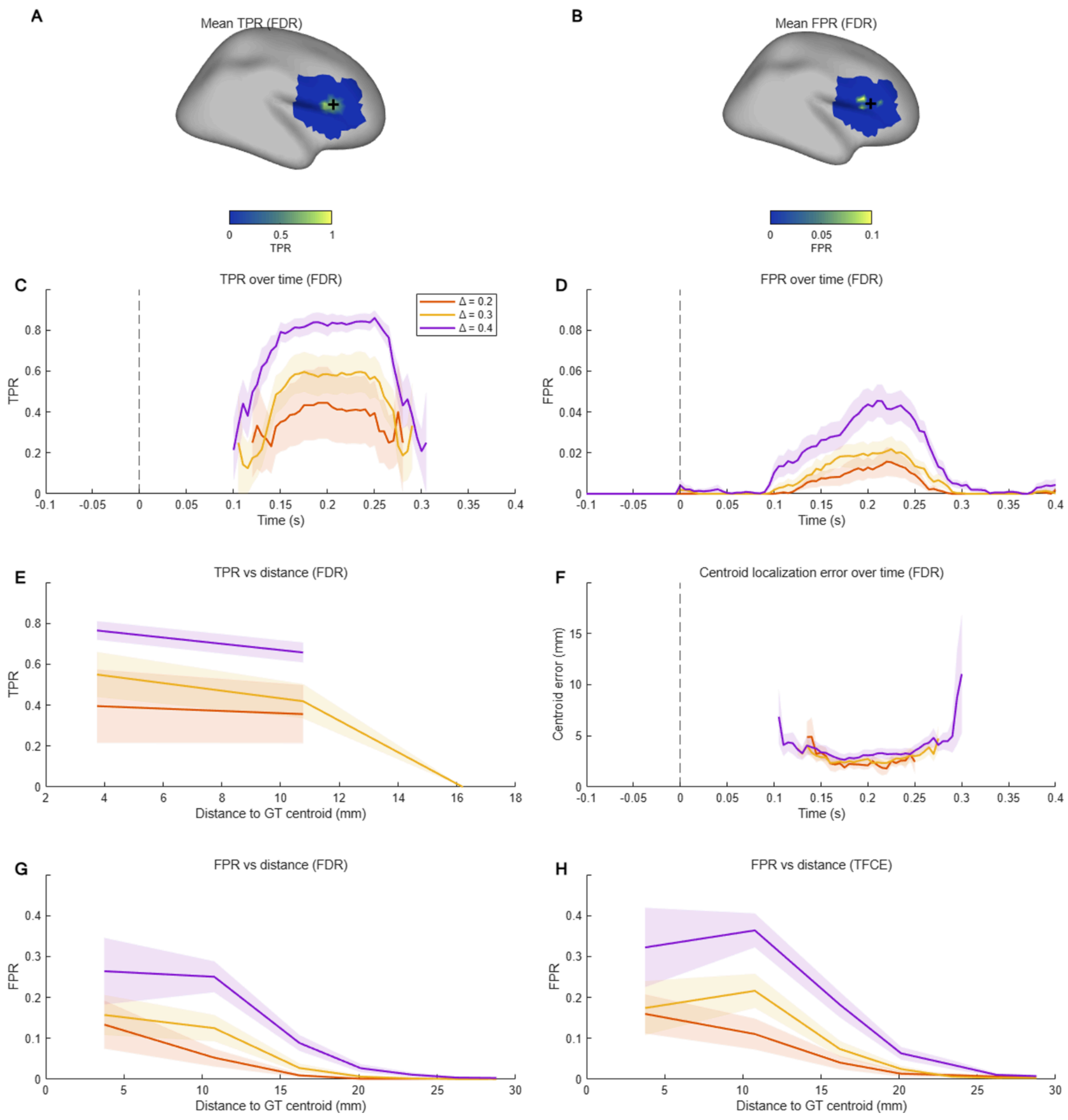


**Supplementary Figure 3. Detection performance of STAMP-LME relative to ground truth.**

A. Vertex-wise true positive rate (TPR) averaged across 0–400 ms for an example simulation in the inferior frontal gyrus (IFG) with an effect size of Δ = 0.3 μV (Fig 7A). Values indicate the fraction of time points at which STAMP-LME detections overlapped with ground truth (GT) detections after FDR correction. The black cross marks the GT centroid.

B. Vertex-wise false positive rate (FPR) averaged across 0–400 ms for the same simulation example. Values indicate the fraction of time points at which STAMP-LME detections occurred outside the GT detections.

C. TPR over time for different simulated effect sizes Δ, averaged across all simulations. Shaded bands denote ±SE.

D. FPR over time for different simulated effect sizes Δ, averaged across all simulations. Shaded bands denote ±SE.

E. TPR as a function of Euclidean distance from the GT centroid for different simulated effect sizes Δ. Shaded bands denote ±SE.

F. Euclidean distance between the centroids of STAMP-LME and GT detections over time for different simulated effect sizes Δ. Shaded bands denote ±SE.

G. FPR as a function of Euclidean distance from the GT centroid for different simulated effect sizes Δ using FDR correction. Curves are averaged across all simulations. Shaded bands denote ±SE.

H. FPR as a function of Euclidean distance from the GT centroid for different simulated effect sizes Δ using TFCE correction. Curves are averaged across all simulations. Shaded bands denote ±SE.

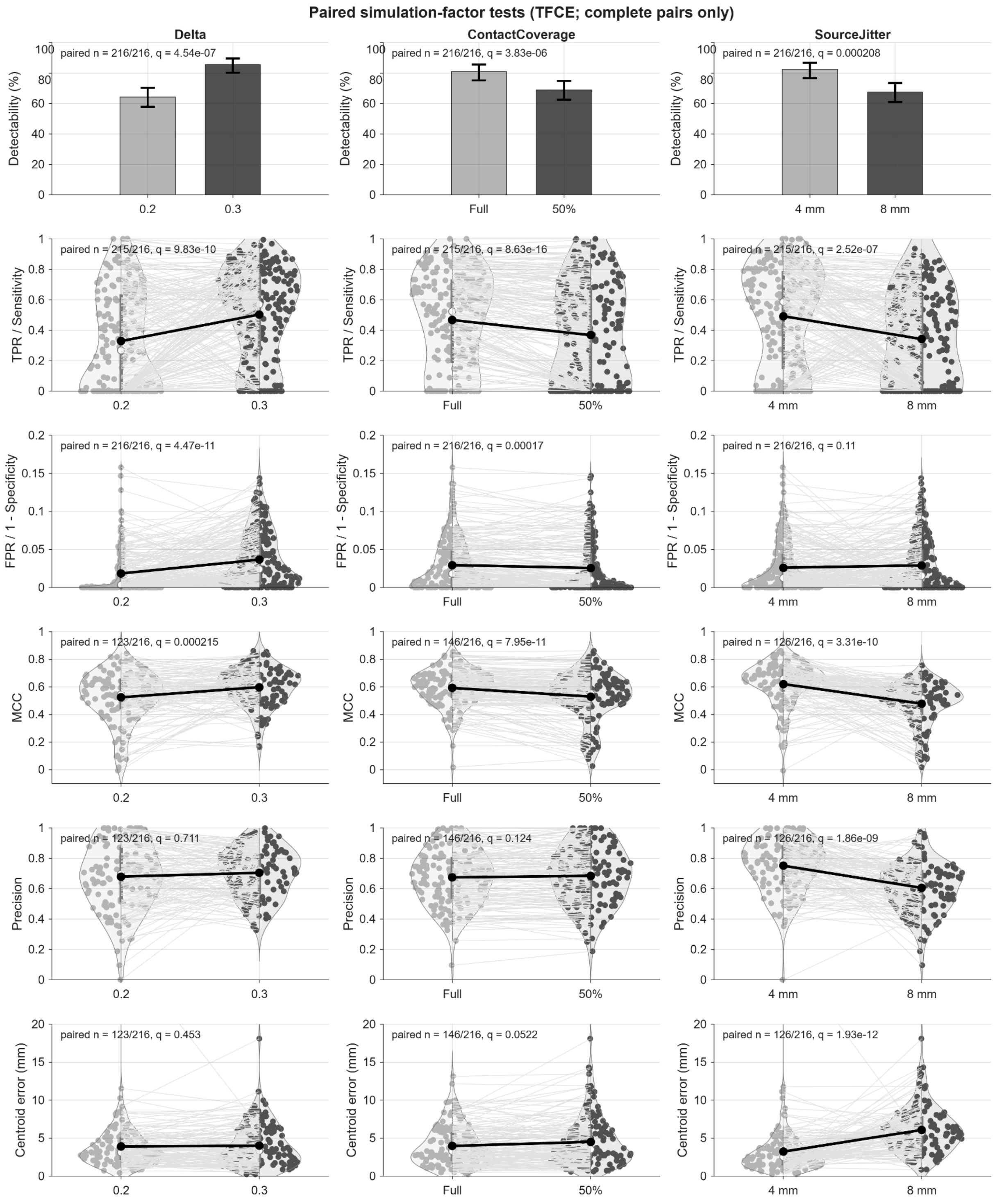


**Supplementary Figure 4. Effects of simulation parameters on the performance of TFCE-corrected STAMP-LME.**

Detectability, spatiotemporal true-positive rate (TPR; sensitivity), false-positive rate (FPR; 1 − specificity), Matthews correlation coefficient (MCC), precision, and centroid localization error are compared as a function of simulated effect size, electrode sampling, and source-location jitter. All metrics were evaluated within the 150–250 ms temporal window. Detectability was defined at the simulation-case level as the presence of at least one TFCE-corrected STAMP-LME detection within the evaluation window.

For vertex–time metrics, TP denotes vertex–time samples detected by both STAMP-LME and GT; FN denotes GT-positive vertex–time samples not detected by STAMP-LME; FP denotes vertex–time samples detected by STAMP-LME but not by GT; and TN denotes vertex–time samples detected by neither STAMP-LME nor GT. TPR was defined as TP/(TP + FN), FPR as FP/(FP + TN), precision as TP/(TP + FP), and MCC as $(TP \times TN - FP \times FN) / \sqrt{[(TP + FP)(TP + FN)(TN + FP)(TN + FN)]}$. TP, FN, FP, and TN were pooled across all valid vertices and time samples within the 150–250 ms interval. Centroid localization error was defined as the Euclidean distance between the centroids of the STAMP-LME and GT detections within the same interval.

Detectability is shown as proportions, with error bars denoting 95% Wilson confidence intervals. Continuous measures are shown as violin plots; dots show complete paired observations, thin lines connect matched simulation pairs, and thick black lines connect group means. Comparisons retained all balanced smoothing levels and remaining simulation conditions without averaging. Effect sizes of $\Delta = 0.2$ and 0.3 were paired within the same run, region, electrode-coverage condition, and smoothing level; full and 50% electrode sampling were paired within the same run, region, effect size, and smoothing level; and source-location jitter of 4 and 8 mm was paired between different runs while matching region, effect size, electrode coverage, and smoothing level.

Detectability was assessed using exact McNemar tests, and continuous measures were assessed using paired Wilcoxon signed-rank tests. Values shown as q are Benjamini–Hochberg FDR-adjusted p-values across the three simulation-factor comparisons within each outcome; n denotes the number of complete pairs evaluated. The number of complete pairs was lower for metrics that were undefined when the required detections were absent, particularly MCC, precision, and centroid localization error.

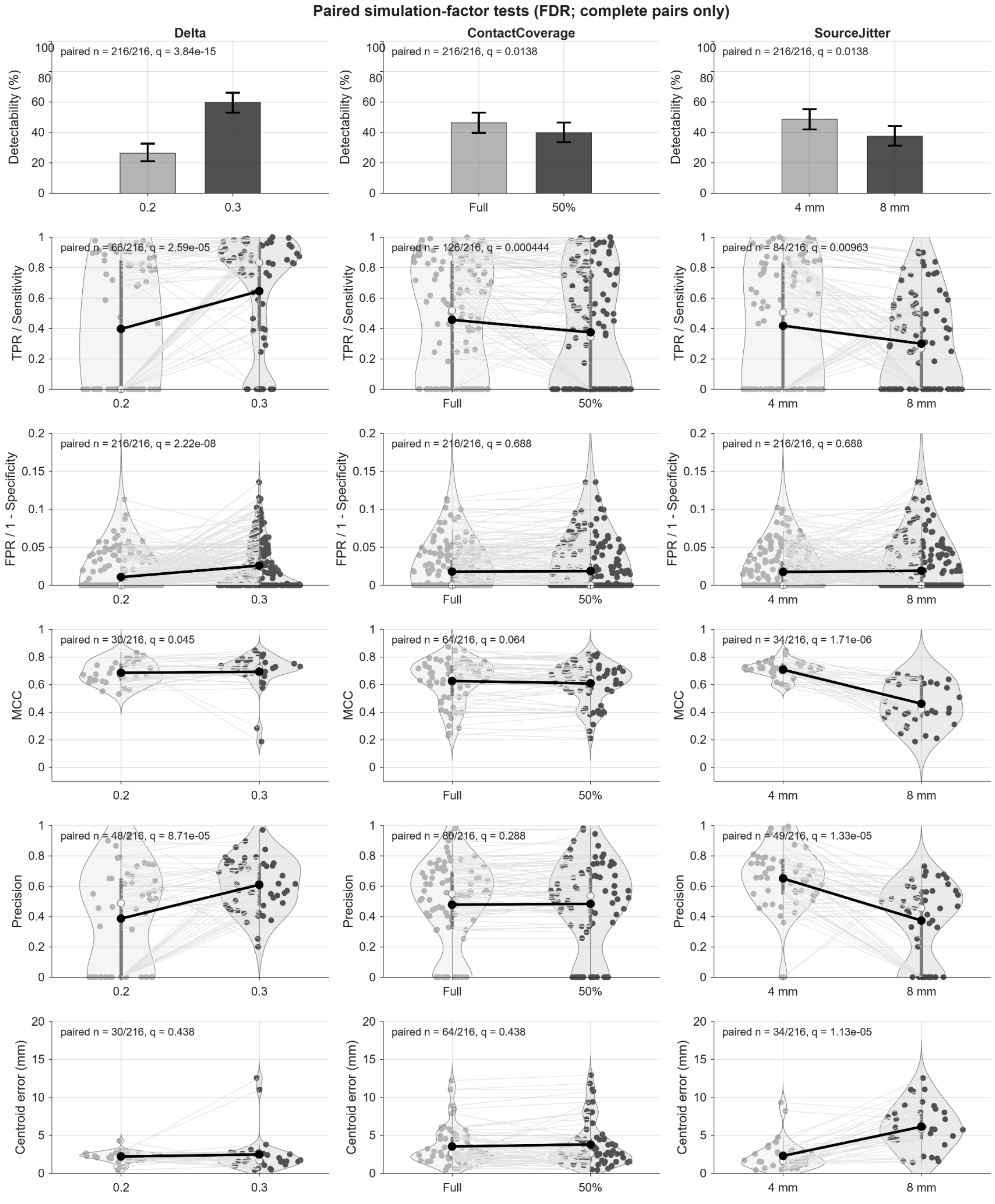


**Supplementary Figure 5. Effects of simulation parameters on the performance of FDR-corrected STAMP-LME.** Same conventions, outcome definitions, pairing scheme, and statistical tests as in Supplementary Figure 4, but using FDR-corrected detections instead of TFCE-corrected detections.

| Scenario | | | Detectability [%] | | | | Precision [%] | | | | MCC [%] | | | | Similarity [%] | | | | Centroid Error [mm] | | | |
|---|---|---|---|---|---|---|---|---|---|---|---|---|---|---|---|---|---|---|---|---|---|---|
| Source Jitter [mm] | Ch | Δ effect size | σ = 3 | σ = 5 | σ = 7 | Best σ | σ = 3 | σ = 5 | σ = 7 | Best σ | σ = 3 | σ = 5 | σ = 7 | Best σ | σ = 3 | σ = 5 | σ = 7 | Best σ | σ = 3 | σ = 5 | σ = 7 | Best σ |
| 4 | Full | 0.2 | 50.0 | 33.3 | 22.2 | 3/5/7 | 48.4 | 36.9 | 45.9 | 3/5/7 | 75.2 | 69.8 | 61.7 | 3/5/7 | 66.1 | 56.2 | 53.7 | 3 | 1.86 | 1.94 | 2.28 | 3/5 |
| | | 0.3 | 77.8 | 72.2 | 55.6 | 3/5/7 | 75.1 | 68.2 | 53.5 | 3 | 70.8 | 70.8 | 64.1 | 3/5 | 75.1 | 66.6 | 63.2 | 3 | 2.23 | 2.45 | 3.61 | 3/5 |
| | 50% | 0.2 | 44.4 | 33.3 | 22.2 | 3/5/7 | 55.2 | 50.1 | 32.6 | 3/5/7 | 70.0 | 66.0 | 58.6 | 3/5/7 | 63.2 | 52.6 | 48.9 | 3 | 1.97 | 2.66 | 2.50 | 3/5 |
| | | 0.3 | 72.2 | 55.6 | 44.4 | 3/5/7 | 76.8 | 60.2 | 53.0 | 3 | 69.5 | 65.8 | 65.8 | 3/5/7 | 72.4 | 63.8 | 60.1 | 3 | 2.36 | 3.30 | 3.07 | 3/5/7 |
| 8 | Full | 0.2 | 22.2 | 22.2 | 16.7 | 3/5/7 | 18.3 | 18.5 | 22.7 | 3/5/7 | 73.7 | 79.0 | 75.8 | 3/5/7 | 57.4 | 47.7 | 46.8 | 3 | 3.25 | 2.71 | 3.03 | 3/5/7 |
| | | 0.3 | 66.7 | 61.1 | 55.6 | 3/5/7 | 47.7 | 41.6 | 35.9 | 3 | 43.6 | 46.9 | 49.4 | 3/5/7 | 61.1 | 54.6 | 53.5 | 3 | 5.35 | 6.44 | 6.80 | 3/5 |
| | 50% | 0.2 | 27.8 | 16.7 | 5.6 | 3/5/7 | 13.6 | 0.0 | 0.0 | 3/5/7 | 53.8 | n.a. | n.a. | 3/5/7 | 53.5 | 44.4 | 41.9 | 3 | 3.89 | n.a. | n.a. | 3/5/7 |
| | | 0.3 | 61.1 | 44.4 | 50.0 | 3/5/7 | 41.7 | 40.1 | 33.8 | 3 | 44.1 | 52.5 | 46.3 | 5 | 58.1 | 53.0 | 50.5 | 3 | 5.91 | 5.83 | 7.33 | 3/5/7 |

**Supplementary Table 3. Performance of STAMP-LME with FDR correction across spatial smoothing kernels.** Detectability, precision, Matthews correlation coefficient (MCC), similarity to the ground truth and centroid-localization error are shown for each combination of source-location jitter, electrode coverage, and simulated effect size. “Best σ” indicates the smoothing kernel or kernels that were not significantly inferior in paired comparisons (corrected for multiple testing). Each table entry was computed after 18 simulations. Note that similarity is independent of the multiple-comparisons correction method used.

Zeros indicate valid metric values equal to zero, whereas n.a. cells indicate undefined metrics. For example, precision is zero when STAMP-LME detections do not overlap with GT detections. Centroid error is instead computed only when both GT and STAMP-LME have significant vertex-time samples within the evaluation window.

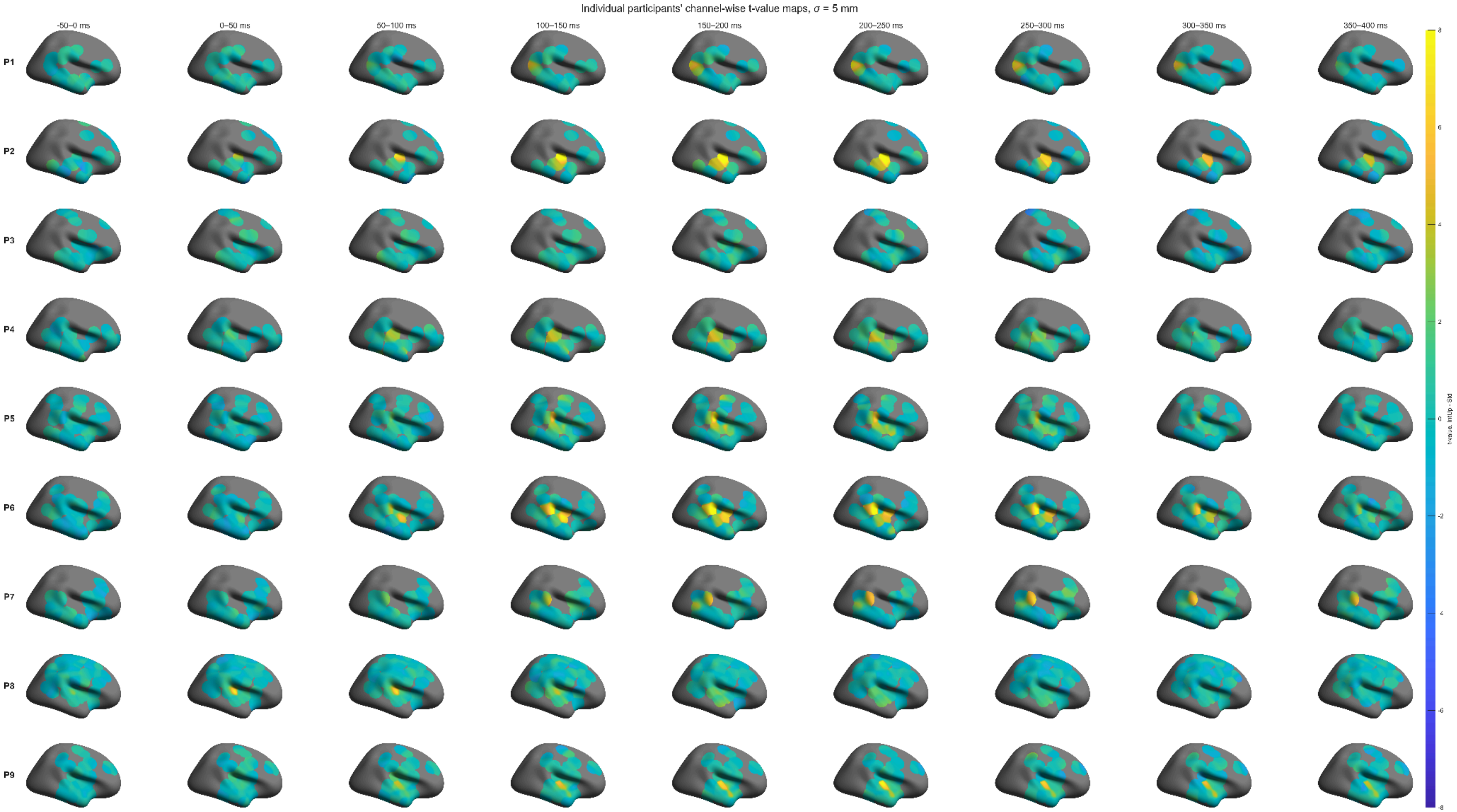
Individual participants' channel-wise t-value maps, σ = 5 mm
-50–0 ms
0–50 ms
50–100 ms
100–150 ms
150–200 ms
200–250 ms
250–300 ms
300–350 ms
350–400 ms
P1
P2
P3
P4
P5
P6
P7
P8
P9
8
6
4
2
0
-2
-4
-6
-8
t-value, IntUp - Std

**Supplementary Figure 6. Individual participants' t-value maps.**

Channel-wise t-values were computed separately for each participant by contrasting IntUp deviant trials with Std trials. The resulting t-values were projected to the fsaverage cortical surface and smoothed with a Gaussian spatial smoothing kernel (standard deviation = 5 mm) to generate surface-level maps. Rows correspond to individual participants (1-9), and columns show 50 ms time-window averages.

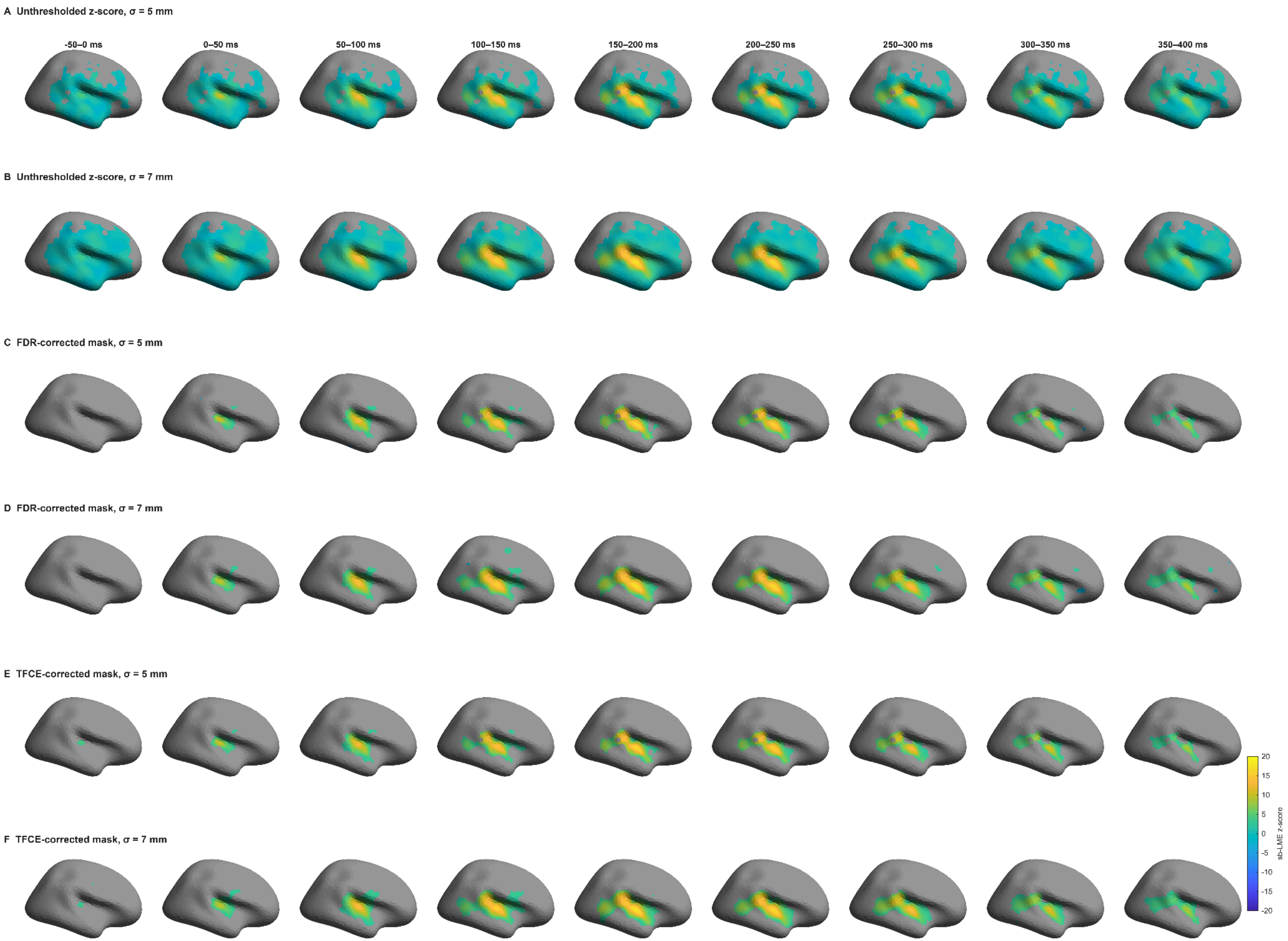
A Unthresholded z-score, σ = 5 mm
-50–0 ms
0–50 ms
50–100 ms
100–150 ms
150–200 ms
200–250 ms
250–300 ms
300–350 ms
350–400 ms
B Unthresholded z-score, σ = 7 mm
C FDR-corrected mask, σ = 5 mm
D FDR-corrected mask, σ = 7 mm
E TFCE-corrected mask, σ = 5 mm
F TFCE-corrected mask, σ = 7 mm
20
15
10
5
0
-5
-10
-15
-20
sb-LME z-score

**Supplementary Figure 7. Effect of spatial smoothing on empirical STAMP-LME results.**

STAMP-LME results for the IntUp versus Std contrast shown for two group-level smoothing kernels. Maps summarize the mean statistic (z-score) in 50 ms time windows.
A, B. Unthresholded z-scored STAMP-LME statistics for group-level spatial smoothing $\sigma = 5$ mm and $\sigma = 7$ mm, respectively.
C, D. Same maps as in (A, B), masked by significant permutation-derived p-values after FDR correction.
E, F. Same maps as in (A, B), masked by TFCE-corrected permutation statistics.
Across smoothing levels, the main activation pattern was preserved, with prominent effects over superior temporal and adjacent perisylvian regions. However, the spatial extent and continuity of the maps depend on $\sigma$. The $\sigma = 5$ mm spatial kernel yields a slightly more focal and spatially coherent pattern, whereas $\sigma = 7$ mm yields smoother, more spatially continuous maps, with a greater tendency to extend or blur focal activations.

Gray vertices indicate locations where STAMP-LME was not evaluated or where the statistical threshold was not met in the corrected maps. For FDR-corrected and TFCE-corrected panels, vertices are color-coded when more than 50% of the time window was significant and otherwise plotted in gray.
